# A cortical basis for perception of internal gut sensations

**DOI:** 10.64898/2026.02.11.705298

**Authors:** Omer Rafael, Stav Shtiglitz, Juliet Miller, Yael Prilutski, Itay Talpir, Ayal Lavi, Yoav Livneh

## Abstract

Interoception, the sensing of internal bodily signals, is essential for brain-body interactions and shapes emotion, cognition, and behavior^1–5^. Subconscious internal signals, including heartbeats or stomach fullness, can rise to conscious awareness, and this process can improve with practice. Disrupted interoception is emerging as a common deficit in diverse psychiatric disorders^1,3,6,7^. Nevertheless, we still lack a fundamental understanding of the neurobiological basis of perception and conscious reporting of internal sensations. Here, we combine genetic access to specific interoceptive primary afferents, with non-invasive optogenetics in mice, to establish a quantitative framework for studying internal perception. Specifically, we developed a behavioral task in which mice report detecting non-invasive optogenetic activation of vagal mechanosensory neurons, establishing “interoceptive psychophysics”. We combine this approach with cellular-resolution imaging and manipulations to reveal the neuronal basis for perception of these internal sensations in the interoceptive insular cortex. While representations of sensory stimuli were consistently observed in insular cortex across different tasks, perceptual reports were only encoded during a more difficult psychophysics task, but not during simple detection. Accordingly, manipulations of insular cortex activity revealed that, although it is generally required for behavioral reports of internal signals, it plays an additional role in supporting perception under conditions of uncertainty. These findings reveal a neural basis for perception of internal sensations and provide a blueprint for future quantitative exploration of other interoceptive modalities.

## Main

Brain-body communication is essential for our physical and mental well-being. Interoception, the sensing of internal bodily signals, plays a central role in these bidirectional interactions^1–5^. Interoception involves both consciously perceived sensations, as well as subconscious processing^3,6,8^. However, it is still unknown how internal sensory signals eventually become conscious percepts that can guide behavior.

Humans can volitionally focus their attention on specific interoceptive sensations, and the capacity to consciously perceive and report them can improve through practice. For example, focusing on one’s heartbeats can improve their detection^3,6–11^ (but see, e.g., ref. 12). Additionally, focusing on gustatory and visceral sensations of fullness during a meal can affect subsequent food consumption^13–15^. The importance of the ability to consciously perceive gastric sensations is also evident in toilet training in early infancy. Compromised sensing of internal bodily changes is a prominent feature in many psychiatric conditions, from obesity and eating disorders, to anxiety, depression, and drug addiction^1,3,6,7^. Nevertheless, the neurobiological basis of conscious and unconscious internal sensing remains poorly understood^8,16^.

Studies of perception of external sensory stimuli have used quantitative behavioral approaches, such as psychophysics (i.e., the quantitative relationship between physical stimuli and their perception), to study the perceptual limits in each sense, as well as the neural basis for conscious vs. unconscious perception^17–22^. The use of such quantitative approaches in interoception has been impeded by the limited ability to quantitatively vary internal stimuli and concomitantly record and manipulate the relevant neural activity^23^. The recent development of ingestible vibrating capsules for clinical applications also paved the way for a new wave of approaches for studying gut interoception in humans, overcoming the limitations of earlier invasive methods such as gastric balloons^24,25^. While such approaches in humans provide invaluable insights, they cannot be easily combined with cellular-resolution neuronal recordings and manipulations, especially not those that benefit from the wide array of genetic and molecular tools that enable mechanistic dissection in animal models, such as mice and flies^23^.

Foundational neuroanatomical and physiological studies established that gastrointestinal mechanosensory signals are conveyed by specialized vagal afferents, and are relayed centrally through the nucleus tractus solitarius (NTS)^26–26^. More recent work in mice has revealed different genetically defined vagal sensory neuronal subtypes in the nodose ganglion, each conveying distinct sensory signals from different organs, including gastrointestinal mechanosensory signals^30–37^ (e.g., aortic blood pressure, gastric stretch, intestinal nutrient sensing, lung chemosensing). The NTS preserves and further organizes organ-and modality-specific visceral representations, including responses to gastric distension^37–36^.

Beyond brainstem pathways, gastric signals engage a distributed cortical network. Gastric distension and stimulation activate insular and somatosensory cortices together with cingulate and posteromedial regions, including precuneus and retrosplenial cortex, in humans and rodents^40–42^. Complementary electrophysiological and fMRI studies of the intrinsic gastric rhythm identified a broad “gastric network” whose activity is coupled to ongoing stomach dynamics and encompasses sensorimotor, visual, insular, cingulate, retrosplenial and dorsal precuneus territories, highlighting prominent posterior-medial cortical involvement in stomach-brain communication^43,44^.

To begin to explore the higher-order cortical processing of interoception, we focused here on the insular cortex (InsCtx), a central node in the brain-wide network for interoception^45,46^. InsCtx, initially identified as the “vagal receptive cortex”^47–50^, was later shown to be the main cortical site that receives gustatory and interoceptive signals from multiple sensory pathways^46,51,52^. Posterior and mid-InsCtx are more closely associated with visceral and somatosensory processing, while anterior InsCtx integrates bodily signals with affective and cognitive information^45,46,51,53,54^. A broadly similar organization is supported by human neuroimaging studies^55–57^. InsCtx thus links external sensations with taste and other internal sensory systems^45,46,46,58–70^. It is therefore thought to mediate the interoceptive aspects of numerous behaviors, from feeding and drinking to emotional regulation, decision-making, and social behaviors^45,46,71–82^.

Here, we sought to investigate how internal bodily signals become sensory percepts and subsequent decisions by focusing on the InsCtx. To do so, we combined non-invasive optogenetic activation of putative vagal gut mechanosensory neurons with cellular-resolution imaging and manipulations of mid-posterior InsCtx. We chose mechanosensation for studying internal perception, as it is a relatively temporally confined stimulus, and also to allow comparison with recent studies in humans^23,24,83^. We used this approach to establish different behavioral assays, including quantitative internal psychophysics. By linking task-dependent perceptual readouts with cellular-scale activity and causal perturbations, our findings advance our understanding of the cortical computations by which internal bodily signals are transformed into decision-relevant representations.

### Non-invasive optogenetic activation of vagal mechanosensory neurons

In order to train mice to report activation of vagal mechanosensory neurons, we developed an experimental system that allows their non-invasive activation using optogenetics. We targeted oxytocin receptor (OxtR) expressing vagal sensory neurons located in the nodose ganglia (hereafter referred to as NG-OxtR^+^ neurons), which have been previously identified as gut mechanosensory neurons^30^. We used the highly sensitive red-shifted opsin, ChRmine, as it has been previously used for non-invasive activation of brain and heart cells^84,85^. We injected a retrograde virus (retroAAV-EF1a-DIO-ChRmine-mScarlet) into the brainstem nucleus tractus solitarius (NTS) of OxtR-t2a-Cre mice to virally express ChRmine in NG-OxtR^+^ neurons and their axons (**Fig. 1a**). This retrograde approach circumvented the need for extensive neck surgery to expose the vagus nerve for virus injections. We confirmed efficient expression in the nodose ganglion (**Fig. 1b**), but not in OxtR^+^ vagal motor neurons (**Fig. 1c**). Moreover, retrogradely labeled NG-OxtR^+^ neurons projected to previously described subregions of the NTS^30^ (**Fig. 1c**). Consistent with previous work identifying NG-OxtR^+^ neurons as gut mechanosensory neurons that innervate the entire intestine^30^, we observed labeled intraganglionic laminar endings (IGLEs) innervating the intestine, which are known to underlie gut mechanosensing^26,27^ (**Fig. 1d; Extended Data Fig. 1a**). We used vagus nerve electrophysiology to validate efficient activation of retrogradely-labeled NG-OxtR^+^ neurons using optogenetic activation of ChRmine via red light illumination of the neck (**Fig. 1e**).

**Figure 1:**
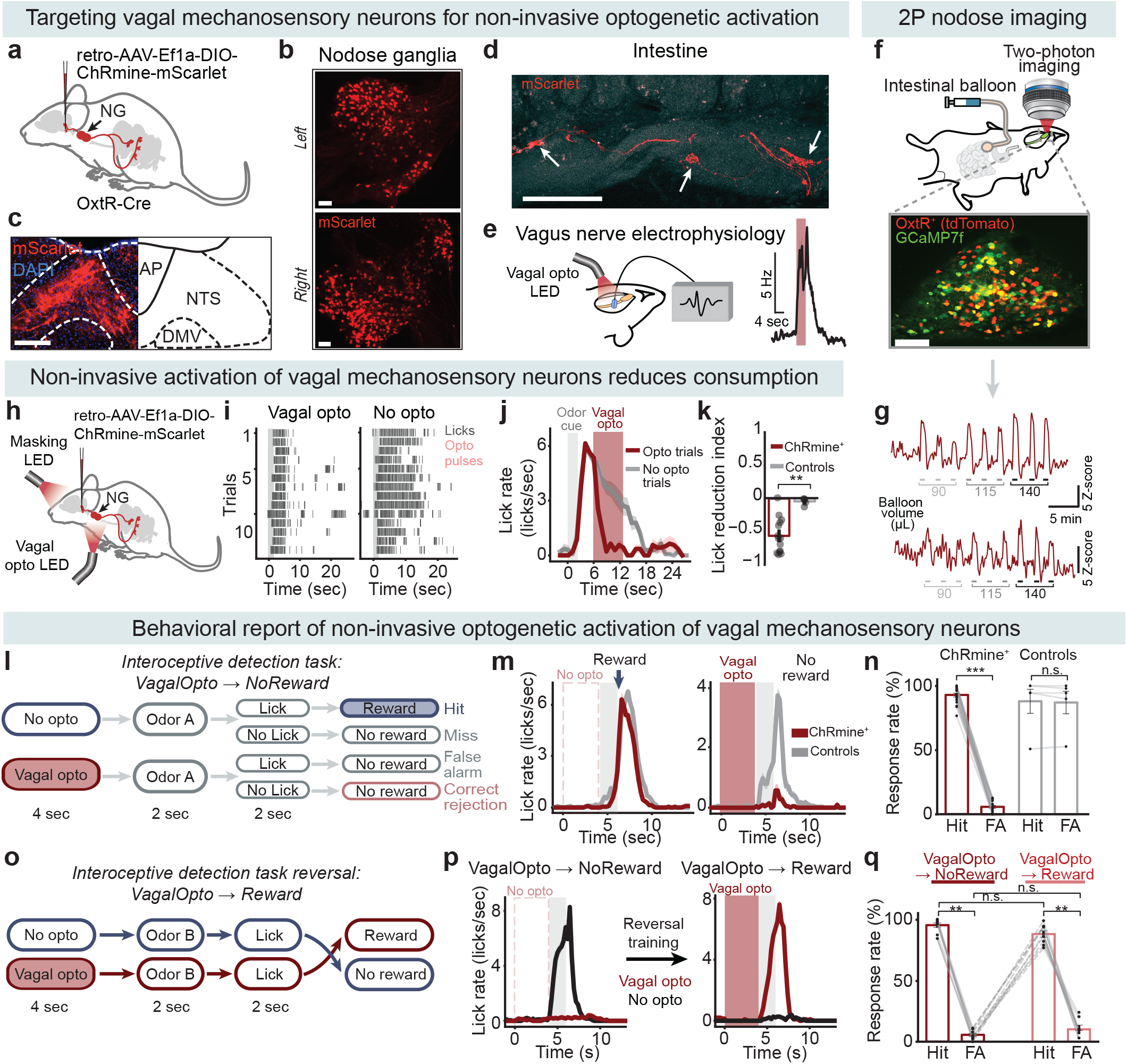
Behavioral report of non-invasive optogenetic activation of vagal mechanosensory neurons. (a) Schematic of the retrograde viral strategy for expressing ChRmine-mScarlet in NG-OxtR^+^ neurons. (b) Example of the right and left NG of an OxtR-Cre mouse expressing Cre-dependent ChRmine-mScarlet following retrograde brainstem injections. Scale bar = 200 µm. (c) NG-OxtR^+^ terminals expressing ChRmine-mScarlet innervating the NTS. Scale bar = 200 µm. (d) NG-OxtR^+^ terminals expressing ChRmine-mScarlet innervating the intestine and forming intra ganglionic laminar endings (IGLEs; labeled with white arrows). Scale bar = 100 µm. (e) Validation of optogenetic activation of retrogradely labeled NG-OxtR^+^ neurons. *Left*: Illustration of vagal electrophysiological recordings during optogenetic activation of retrogradely labeled NG-OxtR^+^. *Right*: Example vagal nerve responses to optogenetic stimulation (red shaded rectangle). Values are mean ± SEM across 9 repetitions. (f-g) Validation of gut mechanosensory responses of NG-OxtR^+^ neurons. (f) *Top*: schematic of 2-photon calcium imaging from vagal sensory NG neurons in OxtR-Cre mice during intestinal balloon inflation used to evoke activation of intestinal mechanosensors. *Bottom*: example field of view of the NG showing OxtR^+^ tdTomato-labeled neurons (red), GCaMP7f-expressing neurons (green), and colocalized OxtR^+^ neurons expressing both tdTomato and GCaMP7f (yellow). Scale bar = 100 µm. (g) Example traces from two NG-OxtR^+^ neurons showing time-locked responses to intestinal balloon inflations. Each inflation trial (indicated in grey) lasted 50 sec, with 3 volumes tested (90 µL, 115 µL, 140 µL), each repeated three times. (h) Illustration of a mouse during non-invasive activation of NG-OxtR^+^ neurons. To avoid visual detection of the optogenetic illumination, we used a masking red light directed towards the mouse’s face. (i) Example lick rasters showing licking behavior of a food restricted mouse during the closed-loop sucrose consumption task. The grey rectangle marks the 2-sec odor cue presentation, and grey ticks mark individual licks. Each red tick represents an optogenetic activation pulse delivered in response to a lick. Vagal opto and no-opto trials were interleaved and are separated here for presentation purposes only. (j) Average lick rate (of the raster in panel ‘i’) during the sucrose consumption task. The grey rectangle marks odor-cue presentation, and the red rectangle marks the optogenetic stimulation time window in optogenetic trials. (k) Quantification of lick reduction during the sucrose consumption task in food restricted mice. Bars show mean ± SEM across mice (ChRmine^+^ n = 14, controls n = 5). ChRmine^+^ mice reduced licking to sucrose during vagal optogenetic stimulation while control mice did not. p = 0.003 (Mann-Whitney U test). Effect size (rank-biserial correlation): r = 0.86. (l) Schematic of the interoceptive detection task: VagalOpto→NoReward. When the odor cue appears without preceding vagal optogenetic stimulation, mice lick operantly and receive a sugar-water reward (∼5µL, 600mM sucrose). When vagal optogenetic stimulation (15Hz) precedes the odor cue, no reward is delivered regardless of licking. Each trial starts with a 1 sec masking light that remains on until the end of the trial and turns off during the inter-trial interval. Mice performed ∼180-200 randomly interleaved trials per session. (m) Example average lick rates of a ChRmine^+^ mouse and a control mouse, both showing similar lick rates in trials with no optogenetic stimulation (no opto; *left*). The control tdTomato^+^ mouse had high lick rates for the VagalOpto→NoReward trial type, whereas the ChRmine^+^ mouse had low lick rates (vagal opto; *right*). The grey rectangles mark odor cue presentation, and the red rectangle marks the optogenetic stimulation in vagal opto trials. (n) Quantification of response rates across mice in the interoceptive VagalOpto→NoReward task, bars show mean ± SEM across mice. All ChRmine^+^ mice (n = 19) reported the detection of non-invasive NG-OxtR^+^ optogenetic activation. They showed high hit rates (licking in trials without optogenetic stimulation, receiving sugar-water reward) and low FA rates (withholding licking in optogenetic stimulation trials), p = 3.6×10^-21^ (paired t-test), Cohen’s d = -12.08. Control mice (n = 5) did not distinguish between opto vs. no opto trials, and showed similar licking responses to both, p = 0.42 (paired t-test). (o) Schematic of the interoceptive detection task reversal, VagalOpto→Reward, in which vagal optogenetic activation (15Hz) before the odor cue predicts sugar-water reward upon licking (∼5µL 600mM sucrose). A different odor cue was used for the VagalOpto→Reward and VagalOpto→NoReward tasks. If the odor was not preceded by the vagal optogenetic stimulation, no reward was delivered regardless of licking. (p) Example average lick rates of the same mouse performing the VagalOpto→NoReward version of the task (*left*), and after reversal training, the VagalOpto→Reward version of the task (*right*). Grey rectangles mark odor-cue presentation, and the red rectangles mark the optogenetic stimulation in vagal opto trials. *Left*: in the VagalOpto→NoReward version, the mouse had high licking rate in no-opto trials (black), and low licking rate for vagal optogenetic stimulation trials (red). *Right*: in the VagalOpto→Reward version, the mouse had high licking rate in the vagal opto trials (red), and low licking rate for the no opto trials (black). (q) Quantification of response rates of the same mice (n=9) during the VagalOpto→NoReward task (*left*) and VagalOpto→Reward task (*right*). Bars show mean ± SEM across mice. Mice showed comparable hit rates (adj.P = 0.099, Wilcoxon with FDR correction) and false-alarm rates (adj.P = 0.164, Wilcoxon with FDR correction) between tasks. Within both the VagalOpto→NoReward and VagalOpto→Reward conditions, hit and false-alarm rates differed, consistent with task learning (VagalOpto→NoReward: adj.P = 0.0078, Effect size (rank-biserial correlation): r = -0.87; VagalOpto→Reward: adj.P = 0.0078, Effect size (rank-biserial correlation): r = -0.87 Wilcoxon test with FDR correction).

Previous work has provided strong experimental support for the gut-related mechanosensory function of NG-OxtR^+^ neurons^30^; yet, they have not been shown to directly respond to gut mechanosensory stimuli. To test this, we performed in vivo two-photon calcium imaging of the nodose ganglion while inflating balloons in the proximal small intestine to evoke gut distention^33,86^ (**Fig. 1f-g**). Retrogradely labeled NG-OxtR^+^ neurons expressed tdTomato in addition to the calcium indicator GCaMP7f, enabling identification and activity monitoring. We found that ∼23% of NG-OxtR^+^ neurons responded to inflation of the intestinal balloon (45/168 neurons from 4 mice). We note that this is likely an underestimation as NG-OxtR^+^ neurons innervate the entire small intestine^30^, while our mechanosensory stimulation covered only a small fraction of the entire small intestine length (∼3-6%, see Methods). These data, together with previous work^30^, directly demonstrate that the retrogradely-labeled NG-OxtR^+^ neurons respond to gut mechanosensory stimulation.

Previous work has demonstrated that activation of NG-OxtR^+^ reduces food consumption in food-restricted mice^30^. We therefore tested whether non-invasive optogenetic activation of these neurons affected consumption. We targeted the somata of ChRmine-expressing OxtR^+^ neurons in the nodose ganglia by pointing red LEDs at their necks (see additional control experiments in the next section). To eliminate potential visual responses to the optogenetic stimulation light, we deployed a masking red light positioned above the animal’s head throughout all trials (**Fig. 1h**). We trained food-restricted mice to operantly lick for a sucrose solution presented in a trial-structured manner, and randomly interleaved trials with non-invasive closed-loop optogenetic activation of NG-OxtR^+^ that was triggered by licking. We observed a clear reduction in licking (i.e., reduced consumption) during optogenetic activation trials compared to control trials (**Fig. 1i-j**), quantitatively confirmed by a significant decrease in the lick-reduction index (**Fig. 1k**). We did not observe this effect in wildtype (C57BL/6) mice or OxtR-t2a-Cre mice injected with a control virus (retroAAV-CAG-FLEX-tdTomato; **Fig. 1k**). We observed no differences between males and females (**Extended Data Fig. 1b**), and consistent with previous work, we observed a similar lick-reduction in water-restricted mice^30^ (**Extended Data Fig. 1c**).

We next conducted control analyses and experiments to test whether the observed reduction in licking during non-invasive optogenetic activation of NG-OxtR^+^ was driven by aversion. First, during the sucrose consumption test, mice consistently resumed licking during subsequent trials without optogenetic activation (**Fig. 1i-k, Extended Data Fig. 1d**), suggesting that the lick reduction effect is acute and does not persist once the optogenetic stimulus stops. To test this quantitatively, we separately compared interleaved trials with vs. without optogenetic stimulation in ChRmine-expressing and control mice. We found that licking was significantly reduced in ChRmine-expressing mice during optogenetic stimulation trials, but did not differ between groups during no-stimulation trials (**Extended Data Fig. 1e**). We therefore reason that this transient suppression of licking likely reflects a reflexive response to vagal stimulation rather than an aversive state. Second, we performed a head-fixed conditioned taste preference task by pairing one type of juice reward with NG-OxtR^+^ activation (**Extended Data Fig. 1f**). Consistent with the immediate lick reduction described above, we observed a marked reduction in licking for the paired juice during stimulation days. However, we only observed a very small change in preference on the subsequent test day without optogenetic stimulation (∼0.5 during conditioning vs. ∼0.06 post-conditioning; **Extended Data Fig. 1f**). Moreover, as shown below, NG-OxtR^+^ activation can itself be learned as an appetitive reward-predicting cue (**Fig. 1o-q**), further suggesting that NG-OxtR^+^ activation does not cause strong aversion. Nevertheless, we cannot rule out that NG-OxtR^+^ stimulation is mildly aversive. These results are also in line with previous work demonstrating that optogenetic activation of NG-OxtR^+^ neurons does not affect place preference to suggest it is not aversive^30^.

To assess how the magnitude of vagal activation evoked by our optogenetic stimulation compares with that produced by physiological intestinal distention, we first estimated the change in intestinal diameter following feeding. Small intestine diameter increased from 2.06 ± 0.33 mm in fasted mice to 3.25 ± 0.23 mm in refed mice allowed access to food for 2 h following fasting, corresponding to a ∼1.5-fold increase (mean ± SEM; n = 4 mice per group). We therefore compared vagus nerve activity evoked by a ∼1.5-fold intestinal distention with that evoked by our (15 Hz) optogenetic stimulation of NG-OxtR^+^ neurons. Although the magnitude of the response varied across mice, the two stimuli produced overall comparable changes in vagal firing rate. This suggests that our optogenetic stimulation induced a change in vagal activity that does not exceed the natural firing rates evoked by intestinal distention (**Extended Data Fig. 1g**). We note, however, that our optogenetic stimulus does not recapitulate the natural pattern of mechanosensory recruitment, in which different intestinal segments are engaged sequentially rather than simultaneously.

### Behavioral report of optogenetic activation of vagal mechanosensory neurons

Recent work has shown that humans can report experimentally induced gut mechanosensations, establishing a powerful approach to study behavioral report of interoception^24^. We used our new experimental approach to test whether mice could be trained to behaviorally report optogenetically induced internal mechanosensations. To this end, we developed Go/NoGo behavioral tasks in which head-fixed mice operantly licked following an odor cue to receive a sugar-water reward, that was predicted by the presence or absence of the optogenetic stimulus prior to the odor cue. Specifically, in one version of this task, optogenetic activation (at 15 Hz) served as a ‘NoGo cue’, indicating no reward or punishment would be delivered upon licking (we refer to this task as “VagalOpto→NoReward”; **Fig. 1l**). The absence of optogenetic activation before the odor cue indicated that licking following the odor will be rewarded by drop of sugar water. Importantly, the odor was the same in both conditions; the only difference between them was the presence or absence of optogenetic activation of NG-OxtR^+^ neurons before the odor (**Fig. 1l**). As described above, we ensured that visual detection of the optogenetic illumination did not bias behavior by using a masking red light throughout all trials as well as light-blocking neck covers (see Methods).

Mice expressing ChRmine in NG-OxtR^+^ neurons successfully performed this task, exhibiting high licking rates in the no-optogenetic (“no-opto”) stimulation trials (‘hits’, receiving reward) and very low licking rates in the optogenetic (“opto”) vagal stimulation trials (‘false alarms’, no reward; **Fig. 1m-n**). In contrast, control mice expressing tdTomato in NG-OxtR^+^ neurons and wildtype (C57BL/6) mice were unable to differentiate the trial types and displayed similar rates of hits and false alarms (**Fig. 1m-n**). Mice learned the VagalOpto→NoReward task rapidly within the first session and performance remained high thereafter (**Extended Data Fig. 1h-j**). We thus tested whether this learning rate depended on the strength of the stimulus. To test this, we trained an additional cohort of mice using 1 Hz optogenetic stimulation instead of 15 Hz in the same VagalOpto→NoReward task. Compared with mice trained using 15 Hz stimulation, learning with 1 Hz stimulation emerged more gradually across training days, consistent with a weaker vagal signal. Nevertheless, mice learned to discriminate opto and no-opto trials, as reflected by increased discriminability index (d’), consistently high hit rates on no-opto rewarded trials, and progressively lower false alarm rates on unrewarded opto trials (**Extended Data Fig. 1k-m**).

The high performance in the VagalOpto→NoReward task (**Fig. 1n**), together with lick reduction in the sucrose consumption test (**Fig. 1h-k**), could potentially be explained by activation of reflexive lick-suppression signal, rather than using this vagal optogenetic activation flexibly as a learned cue. To test this, we used a reversed version of the task (**Fig. 1o**). In this reversed version, when optogenetic stimulation preceded the odor cue, it served as a ‘Go signal’, indicating that licking would result in sugar water reward delivery (‘hit’). Conversely, in the absence of optogenetic stimulation, licking in the presence of the same odor cue did not result in a reward (‘false alarm’). We refer to this task as “VagalOpto→Reward” (**Fig. 1o**). We note that we used a different odor from the one used in the VagalOpto→NoReward task. Importantly, the optogenetic stimulation parameters (duration, intensity) were identical for all trials in both the VagalOpto→NoReward and VagalOpto→Reward task versions, and the only difference between trials was the assigned behavioral meaning of the optogenetic stimulation. We successfully trained the same mice to perform both VagalOpto→NoReward and the subsequent reversal to the VagalOpto→Reward task (**Fig. 1p**). Mice demonstrated comparable high hit rates and low false alarm rates when performing both tasks (**Fig. 1q**). This indicates that mice could flexibly associate the optogenetic stimulation as predicting the presence or absence of reward following an odor cue. Furthermore, behavioral performance of mice expressing ChRmine in NG-OxtR^+^ neurons (quantified using d’) improved across training days in the VagalOpto→NoReward task, and remained similarly high during the reversal VagalOpto→Reward task (**Extended Data Fig. 1h**). We note that in Go/NoGo tasks such as ours, performance can be influenced not only by sensory discriminability but also by shifts in decision criterion or response propensity^22^. This is especially relevant for our stimulus as it naturally suppresses consumption (**Fig. 1i-k**). Nevertheless, this concern is mitigated by the reversal experiments, which show that the same vagal stimulus successfully supported both action withholding and action execution depending on its learned contingency. This argues against a fixed stimulus-evoked suppression bias. Importantly, control tdTomato-expressing mice and wildtype (C57BL/6) mice lacking ChRmine, could not successfully perform the task and this did not improve across multiple training sessions (**Extended Data Fig. 1h**). We note that since our optogenetic stimulus does not recapitulate the natural pattern of sequential mechanosensory recruitment, we cannot generalize perceptual performance to the full phenomenology of visceroception.

We next asked whether mice relied on the specific odor cue’s identity or could generalize the learned vagal optogenetic stimulation contingency across different odor cues. After animals acquired the task, we replaced their known odor cue with a new odor, and observed that it did not impair performance. Mice continued to show high discriminability, with high hit rates and low false alarm rates (**Extended Data Fig. 1n**). These results indicate that mice generalized the learned association between vagal optogenetic activation and reward availability across odor identities, rather than relying on an association with a specific odor cue. Conversely, switching the behavioral contingency during a session from VagalOpto→NoReward to VagalOpto→Reward disrupted performance and altered behavioral responses (**Extended Data Fig. 1o**). This indicated that behavioral responses depended on the learned contingencies of vagal optogenetic stimulation, and required gradual relearning (compare to **Fig. 1p,q**). To characterize the temporal constraints of the vagal optogenetic percept, we next varied either the duration of stimulation or the delay between vagal optogenetic stimulation and the odor cue.

First, we varied the duration of the optogenetic stimulus, i.e., the number of optogenetic pulses delivered before the odor cue. In mice initially trained on the original task with 4 sec stimuli, we randomly interleaved different optogenetic stimulus durations (all at 15 Hz): the original 4 s (60-pulse) 1 s (15 pulses), ∼267 ms (4 pulses), or a single ∼6.7 ms pulse (**Extended Data Fig. 1p-q**). Mice showed the highest performance to 60 pulses, and performance progressively declined as stimulation was shortened (**Extended Data Fig. 1p**). This impairment was primarily reflected by an increase in false alarm responses during vagal opto trials, while hit rates on no-opto rewarded trials remained high, suggesting reduced detection of shorter stimuli (**Extended Data Fig. 1q**). Thus, brief activation of NG-OxtR+ vagal neurons was sufficient to influence behavior, but reliable behavioral use of the signal depended on stimulation duration, indicating that the evoked interoceptive signal is integrated over time rather than acting as an all-or-none trigger.

Second, we asked how long the vagal optogenetic signal remained behaviorally accessible after optogenetic stimulation ended. To test this, we initially trained mice on the original task with no delay between the optogenetic stimulus offset and odor cue onset. Then, we randomly interleaved different delays between the end of vagal optogenetic stimulation and the odor cue (2, 4, 6, and 8 s delay). Behavioral performance declined with longer delays (**Extended Data Fig. 1r**). As in the variable duration experiment above, this decline was driven mainly by increased false alarm responses on vagal opto trials, whereas hit rates on no-opto rewarded trials remained high (**Extended Data Fig. 1s**). These results indicate that the evoked internal mechanosensory signal has a limited temporal persistence: it can guide behavior when the report odor cue follows shortly after stimulation, but its influence decays over a few seconds.

Finally, although we verified that retrogradely labeled vagal OxtR^+^ neurons respond to gut mechanosensory stimulation, these retrograde injections could potentially label other non-vagal OxtR^+^ neurons in the brainstem. To rule out the contribution from other neurons, we injected retrograde ChRmine to the NTS as described above, and also ablated NG-OxtR^+^ neurons directly by bilaterally injecting a Cre-dependent Caspase-expressing virus in the nodose ganglion (**Extended Data Fig. 2a,b**). Mice with near-complete ablation of NG-OxtR^+^ did not successfully learn the VagalOpto→NoReward task, even after several training days (**Extended Data Fig. 2c**). This verified that NG-OxtR^+^ neurons were necessary for performing the behavioral task.

Previous work using a different OxtR-Cre mouse line (BAC transgenic mice) reported that chemogenetic activation of NG-OxtR^+^ neurons induced significant reductions in body temperature, resulting in torpor (also accompanied by changes in blood pressure and heart rate)^87^. In contrast, previous work using the OxtR-t2a-Cre knock-in mouse line we use here, observed no change in body temperature and blood pressure^30^. We therefore also tested this in our preparation. We conducted heart rate measurements in anesthetized mice, previously trained on both the VagalOpto→NoReward and VagalOpto→Reward tasks. We observed heart rate reductions in response to optogenetic activation in some mice but not in others. Most importantly, mice with or without a change in heart rate showed similar high performance in the VagalOpto→NoReward and VagalOpto→Reward tasks (**Extended Data Fig. 2d)**. We also measured body temperature during performance of the VagalOpto→NoReward task, and did not observe a reduction in body temperature upon NG-OxtR^+^ stimulation, nor other evidence of torpor (**Extended Data Fig. 2e**).

Together with our anatomical identification of NG-OxtR^+^ intestinal IGLEs (**Fig. 1d; Extended Data Fig. 1a**), the responses of NG-OxtR^+^ neurons to intestinal distention (**Fig. 1f-g**), and the acute reduction in consumption induced by their activation (**Fig. 1i-k**), these findings support a substantial contribution of gut-innervating NG-OxtR^+^ neurons to the detected signal. However, NG-OxtR^+^ neurons are heterogeneous in their peripheral targets, with subsets reported to innervate the aortic arch and pancreas^37,87^. Thus, our optogenetic stimulation could potentially co-activate afferents from multiple organs. To test whether aortic arch input is required for behavioral detection, we performed aortic depressor nerve denervation, eliminating vagal sensory innervation of the aortic arch, previously reported to be a major source of NG-OxtR^+^ cardiovascular innervation^87^ (**Extended Data Fig. 2f-g**). Mice with verified denervation readily performed the optogenetic detection task (**Extended Data Fig. 2h**), demonstrating that aortic arch input is not necessary for behavioral report of NG-OxtR^+^ activation. Pancreatic sensory vagal afferents have been implicated in conveying metabolic signals related to glucose homeostasis^88^, and their contribution to the sensory signal evoked by our stimulation cannot be excluded. However, such signals are expected to operate on slower timescales than the sub-second behavioral reports observed here, further supporting a likely significant contribution of gut-related vagal signals to behavioral detection.

### Cortical activity during internal mechanosensory psychophysics

Psychophysical approaches in external senses have long been used to link changes in stimulus intensity with perception and neural activity^17–22^. We adapted this approach to interoception and extended our vagal optogenetic task to enable psychophysical testing of internal sensations. This requires systematically varying stimulus intensity and examining behavioral report. We therefore modified the strength of the optogenetic stimulus by varying pulse frequency, from 0.5 Hz to 15 Hz (randomly interleaved), keeping the duration fixed at 4 seconds (**Fig. 2a**). We chose this approach instead of modifying illumination intensity directly because different light intensities can alter absorption in the illuminated neck tissue (skin, fat, muscles, salivary glands, etc.), precluding direct quantitative comparison across intensities. Instead, we leveraged the fact that the slow kinetics of ChRmine^86^ summate pulses across the four-second stimulation time-window, to effectively stimulate NG-OxtR^+^ neurons with different intensities.

**Figure 2:**
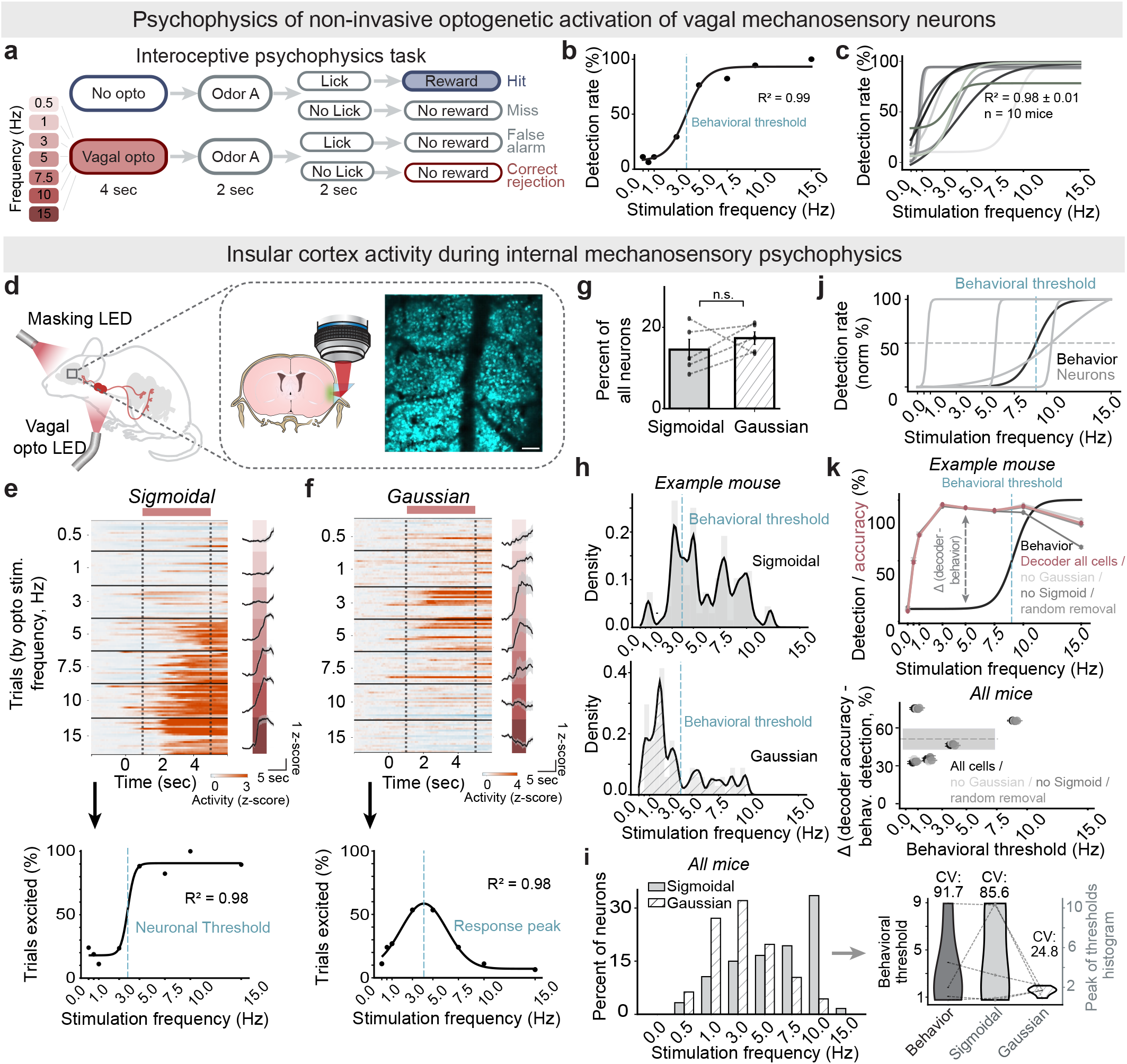
InsCtx activity during internal mechanosensory psychophysics. (a) Schematic of the interoceptive psychophysics task. (b) Example of behavioral psychometric function from one mouse. Dots represent detection rate (%) at each stimulation frequency; the black line shows a sigmoidal fit. The blue dashed line indicates the behavioral threshold, defined as the frequency at 50% detection rate. R² was used to quantify goodness of fit of the sigmoidal function. (c) Behavioral psychometric functions across all mice (n = 10). R² was used to quantify goodness of fit of the sigmoidal function (mean ± SEM across mice). (d) Schematic of the experimental approach: non-invasive optogenetic activation of NG-OxtR^+^ neurons combined with two-photon calcium imaging of InsCtx through a microprism. Schematic coronal brain section further illustrates the approach, also with an example field-of-view. Scale bar: 100 µm. (e–f) Example neurons with Sigmoidal (e) or Gaussian (f) frequency–response functions. *Top*: trial-by-trial z-scored activity sorted by stimulation frequency. Pink bar indicates the optogenetic vagal stimulation period. *Bottom*: black dots represent the observed fraction of trials in which the neuron was significantly excited at each stimulation frequency; the solid black line shows the fitted curve. R² was used to quantify goodness of fit. Dashed vertical line: neuronal threshold (e) or response peak (f). (g) Fraction of neurons classified as Sigmoidal vs. Gaussian out of all recorded neurons per mouse. Bars show mean ± SEM across mice (n = 5); dots indicate individual mice. n.s.: not significant, p =0.41(paired t-test). (h) Distributions of neuronal thresholds (*Top*: Sigmoidal, n=116 neurons) and response peaks (*Bottom*: Gaussian, n=71 neurons) from one example mouse (gray bars: histogram bins, black line: fit). The blue dashed line indicates the behavioral threshold, defined as the frequency at 50% detection rate. (i) *Left*: Distribution of neuronal thresholds (Sigmoidal) and response peaks (Gaussian), pooled across all recorded neurons from all mice (Sigmoidal: n=367 neurons, 73.4 ± 32.1 neurons per mouse; Gaussian: n = 461 neurons, 92.2 ± 42.2 neurons per mouse; from 5 mice). *Right*: Coefficient of Variation (CV) of the peak values from the neuronal threshold (Sigmoidal) and response peak (Gaussian) histograms shown in ‘h’, calculated separately for each mouse. The behavioral CV was computed from the distribution of behavioral thresholds across mice. Note that Sigmoidal thresholds are broadly distributed across stimulation frequencies, whereas Gaussian peaks are concentrated at lower frequencies (1 and 3 Hz), independent of each mouse’s behavioral threshold. (j) Example from one mouse showing the behavioral psychometric fit (black) alongside several Sigmoidal neuronal fits (gray). The example neurons illustrate thresholds lower than the behavioral threshold (more sensitive) and thresholds higher than the behavioral threshold (less sensitive). The blue dashed line marks the behavioral threshold, defined as the frequency at 50% detection rate. (k) Comparison of behavioral detection and neuronal population decoding. *Top*: example mouse showing the behavioral psychometric fit (black) and neuronal SVM decoder accuracy using all recorded neurons (red). Additional decoder curves show performance after removal of different neuronal groups: Gaussian neurons, Sigmoidal neurons, or a randomly selected matched number of neurons. The blue dashed line marks the behavioral threshold, defined as the frequency at 50% detection rate. The gray arrow indicates the frequency used for the calculation of the difference between decoder accuracy and behavioral detection at the bottom (chosen as two frequency steps below the behavioral threshold). *Bottom*: Analyses of all mice. Difference in neuronal decoding accuracy (y-axis) at the chosen frequency below each mouse’s behavioral detection threshold (x-axis, see Methods). The difference (Δ) was calculated as neuronal decoding accuracy minus behavioral detection rate at the indicated frequency. Note positive differences in all mice, demonstrating that neuronal population decoding is more sensitive than behavioral report.

To test the behavioral sensitivity to different frequencies when training only on the highest frequency, we first trained mice on the VagalOpto→NoReward task using the strongest stimulus (15 Hz). Training lasted until mice reached stable high performance, indicating that they had learned the task well (performance on the last day before psychophysics testing: opto correct rejection rate = 62.8 ± 2.6%, d′ = 3.1 ± 0.3; mean ± SEM, n = 10 mice). We then conducted 1-2 separate psychophysics testing days in which optogenetic stimuli were randomly drawn from the entire frequency range, using the same odor cue (i.e., the other frequencies were introduced without prior training). Importantly, we based the psychophysics task on the VagalOpto→NoReward task, in which the absence of optogenetic stimulation signaled reward. We chose this task so that the number of rewards mice received did not depend on their behavioral detection performance (50% reward trials). On the psychophysics day, mice exhibited classic sigmoidal detection curves, similar to detection of external sensory stimuli, and reached near-perfect detection performance at the strongest stimulus (15 Hz, maximal correct rejection rate = 68.5 ± 1.5 %; mean ± SEM, n = 10 mice; sigmoidal fit R² = 0.68 ± 0.01; **Fig. 2b-c; Extended Data Fig.3a**). Together with our vagal recordings (**Extended Data Fig. 1g**), the behavioral thresholds we observed (mostly ∼1-3 Hz), are potentially ∼5-20% of the post-meal firing of vagal mechanosensory neurons. To test whether performance was influenced by recent reward history, we repeated the analysis separately for trials following rewarded and non-rewarded trials and found very similar behavioral thresholds, indicating that the psychometric curve shape and detection threshold were not affected by reward history (**Extended Data Fig. 3b**). These results demonstrate that non-invasive optogenetic activation of vagal mechanosensory neurons can be used to quantitatively study mouse interoception.

Having established a robust psychophysics framework, we next sought to elucidate the neural basis of perception of internal mechanosensations, focusing on the interoceptive InsCtx, initially described as the “vagal receptive cortex”^46,50,60,61^. InsCtx has been shown to respond to gut mechanosensations and to regulate gut motility^52,53,62,63^. Moreover, in human subjects, consciously focusing attention on interoceptive signals engages the InsCtx^57,64^.

To verify that InsCtx responses to the vagal optogenetic stimulus depended on NG-OxtR^+^ neurons, we measured bulk InsCtx calcium activity using fiber photometry together with the same caspase-based ablation approach described above (**Extended Data Fig. 3c**). Vagal optogenetic stimulation evoked robust InsCtx responses in sham-ablation animals, whereas these responses were completely absent in NG-OxtR^+^ ablated animals (**Extended Data Fig.3d**). To examine the representation of these signals with single-neuron resolution during behavior, we imaged the activity of GCaMP7f expressing layer 2/3 mid-posterior InsCtx neurons via a microprism^65,66^ while animals performed the psychophysics task (**Fig. 2d**). We initially focused only on responses to the vagal optogenetic stimulus, and not on the subsequent reward responses (but see **Fig. 4** below).

We quantified neural responses using auROC analysis, comparing neural activity during baseline and vagal optogenetic stimulation epochs, and determined response significance for each trial using permutation tests. We thus identified InsCtx neurons responsive to vagal optogenetic stimulation and then characterized the temporal decay of their response. We examined the deconvolved responses and found that they decayed to two-thirds of their peak amplitude within 0.85 ± 0.47 s, reaching 35.6 ± 8.6% of their peak response amplitude by the offset of the optogenetic stimulus (onset of the odor cue, mean ± SEM, **Extended Data Fig. 3e**). This vagal optogenetic signal decay time-course is consistent with the behavioral data above examining variable delays between the optogenetic stimulus offset and the odor cue onset (**Extended Data Fig. 1r,s**). Additionally, these results preclude the possibility that the vagal optogenetic stimulus induces a longer-term change in InsCtx activity that persists beyond the current trial.

To characterize InsCtx neuronal response profiles to vagal optogenetic stimulation, we fitted both Sigmoid and Gaussian models for each neuron, and chose the model with the best fit (highest R²). We only considered neurons whose fits were substantially better than shuffled data fits (see Methods). We found that 14.5±2.6% neurons had classic Sigmoidal “neurometric” response patterns^21^, in which neuronal responses increased as stimulus intensity increased (**Fig. 2e; Extended Data Fig. 3f**; see also **Extended Data Fig. 3g-h** for the less prevalent suppression responses). Other neurons were more tuned to specific stimulus frequencies and exhibited Gaussian shaped tuning curves (17.4±1.6%; **Fig. 2f; Extended Data Fig. 3f-i**). Together, these two populations accounted for 32±3% of neurons across mice, and were similarly prevalent (**Fig. 2g**). These response patterns were not explained by licking behavior, as mice did not lick during the vagal optogenetic stimulation in the psychophysics task (**Extended Data Fig. 3j**). They were also not explained by trial history, as we observed similar neuronal tuning when separately analyzing trials following rewarded and non-rewarded trials (**Extended Data Fig. 3k**). Most Sigmoidal neurons response thresholds (i.e., 50% of max response) were centered around the behavioral threshold (**Fig. 2h, top**). In contrast, the peak response frequency of Gaussian neurons were mostly independent of the behavioral threshold, suggesting they were more tuned to physical stimulus properties (**Fig. 2h, bottom**). Across mice, the peak responses of Gaussian neurons were more concentrated at lower frequencies (**Fig. 2i**, **left**), regardless of the individual mouse behavioral threshold (**Fig. 2i, right**). In contrast, Sigmoidal neurons’ thresholds were more widely distributed (**Fig. 2i, right**), and this difference was not explained by reward history (**Extended Data Fig. 3l**). Sigmoidal neurons and Gaussian neurons were spatially intermingled with no consistent spatial organization (**Extended Data Fig. 3m,n**).

Since some neurons were substantially more sensitive than the behavioral detection threshold (**Fig. 2j**), we tested whether population activity could more accurately detect the stimulus than behavior itself. We trained a population decoder on neural activity and found that we could indeed decode behaviorally undetected stimuli, suggesting that the neural population was more sensitive than the behavioral report (**Fig. 2k, top**). We then retrained the decoder after removing different groups of neurons to test how strongly decoding depended on specific response classes. Decoder performance was largely preserved after removing Gaussian cells, Sigmoidal cells, or a randomly selected matched subset of neurons, with only a modest reduction at 15 Hz after removal of Sigmoidal cells (**Fig. 2k**). To quantify this across mice, we sought to train the decoder on sufficiently high frequencies that could activate neurons, while ensuring that the animal could not reliably detect them behaviorally. We trained the decoder for each mouse on a lower frequency than its behavioral report threshold, calculated the difference between them, and found that the decoder had higher accuracy than behavioral report in all mice, regardless of their specific behavioral report threshold (all had positive values; **Fig. 2k, bottom**, see Methods). Moreover, the decoder trained on the lowest frequency (0.5 Hz) still exhibited above-chance performance, even though 0.5 Hz was well below the detection threshold of all mice (**Extended Data Fig. 3o**). When training the decoder on the highest frequency (15 Hz), it could not decode lower frequencies, consistent with the distinct frequency tuning profiles of different neurons that we observed (**Extended Data Fig. 3o**). Decoder performance remained largely preserved after removing different cell classes, although removing Sigmoidal neurons decreased decoding accuracy at lower stimulation frequencies for the 15-Hz-trained decoder (**Extended Data Fig. 3o**).

### A neural basis for detection of internal mechanosensations

We next focused on elucidating the neural basis of detected versus undetected stimuli (i.e., “correct rejections” and “false alarms”). We observed neurons that had differential responses on detected vs. undetected trials during the vagal optogenetic stimulation, prior to the odor cue and to the behavioral response (**Fig. 3a**). To quantify this at the level of individual neurons, we used an auROC-based analysis, often used for single-neuron decoding of behavioral report, termed “choice probability”^67^. We compared detected vs. undetected vagal optogenetic stimulation trials, calculated choice probability, and performed permutation tests within and across conditions to determine statistical significance (**Fig. 3b**; see Methods). We found that 15.4±4.0% of neurons exhibited significant decoding of detected vs. undetected trials during vagal optogenetic stimulation in the psychophysics task (**Fig. 3b**; **Extended Data Fig. 4a**). Surprisingly, when we performed the same analysis for the 1-frequency VagalOpto→NoReward task (**Fig. 1l**), we found very little single-neuron decoding of detected vs. undetected stimuli (6.2±2.4% with a false positive rate of 5%; **Fig. 3c**), and this was independent of specific statistical thresholds (**Extended Data Fig. 4b**). Both Gaussian and Sigmoidal neurons showed robust single-neuron decoding in the psychophysics task (**Extended Data Fig. 4c**). In some mice (3/5) we observed a spatial organization of neurons with similar choice probability values (**Extended Data Fig. 4d-e**).

**Figure 3:**
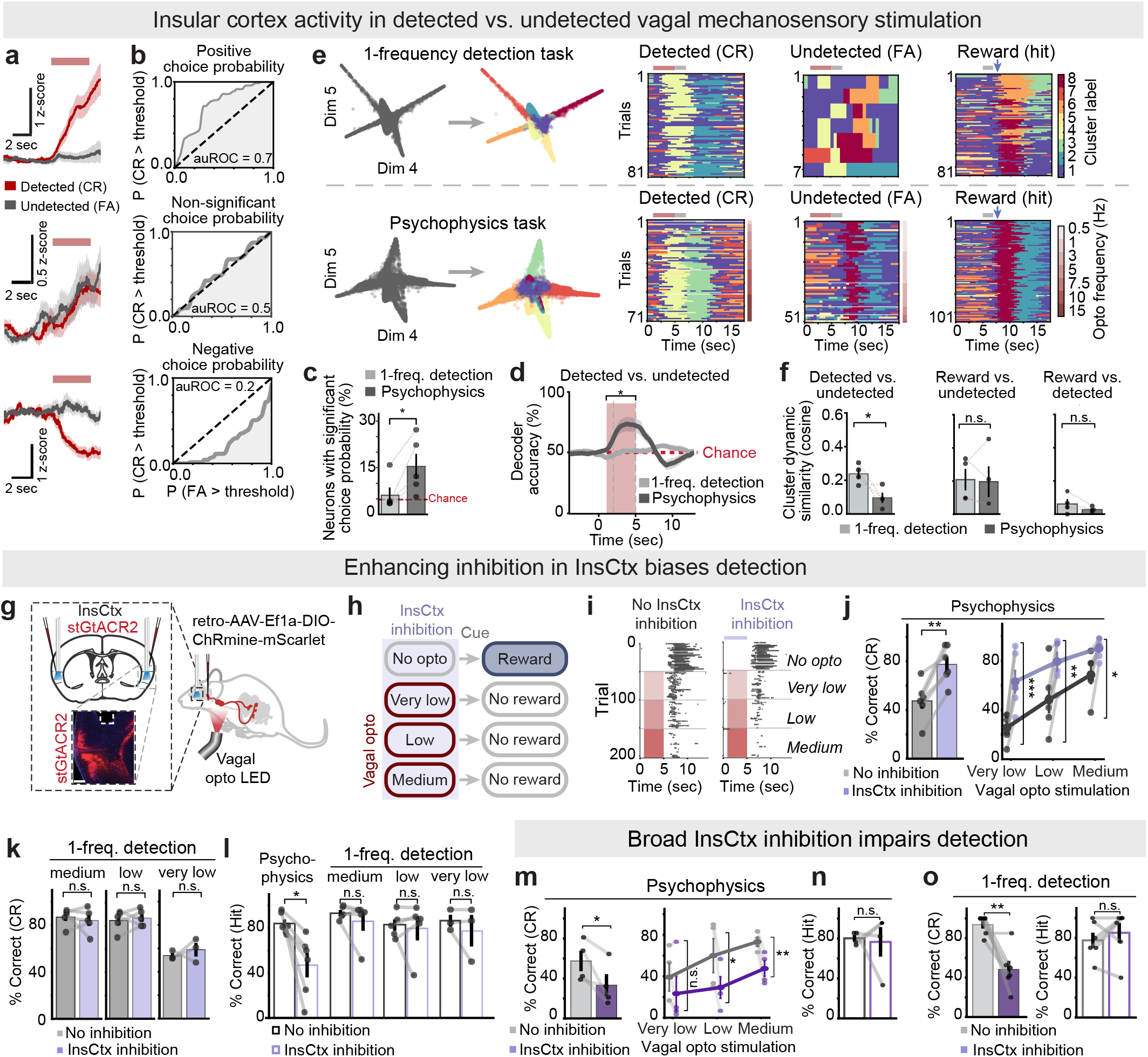
InsCtx activity basis for detection of internal mechanosensations. (a) Example InsCtx neuronal responses during detected trials (CR, red) and undetected trials (FA, black). Pink bars indicate the vagal optogenetic stimulation period. Values are mean ± SEM across trials. (b) Single neuron decoding of detected vs. undetected trials (choice probability): area under the receiver operating characteristic (auROC) for the neurons shown in ‘a’. For each neuron, activity on detected (CR) vs. undetected (FA) trials was compared by sweeping a decision criterion from the minimum to the maximum observed response. At each criterion, we plotted the true positive rate, P(activity_CR > criterion), against the false positive rate, P(activity_FA > criterion; see Methods). The area under this curve quantifies choice probability. *Top*: positive choice probability (auROC > 0.5). *Middle*: non-significant (auROC ≈ 0.5). *Bottom*: negative (auROC < 0.5). Dashed diagonal line: chance. (c) Fraction of neurons with significant choice probability (positive or negative) in each task, out of all recorded neurons. Bars show mean ± SEM across mice (n = 5); dots indicate individual mice. Red dashed line: 5% chance (false positive rate). p = 0.028 (paired t-test), Cohen’s d: 1.5. (d) Time course of SVM decoder accuracy for detected vs. undetected trials (mean ± SEM, n = 5 mice). Red dashed line: chance (50%). Pink bar: vagal opto stimulation time window. Gray dashed vertical line: time period used to train the decoder. Comparison of decoding accuracy between the two tasks during the vagal opto stimulation time window: p=0.043 (paired t-test), Cohen’s d: 1.3. (e) *Left*: Representative planes of dimensionality-reduced population activity over time for the 1-frequency detection task (*top*) and the psychophysics task (*bottom*). Each point represents the population activity pattern in a 0.5 s time bin. Colors indicate manifold-cluster identity assigned by the automated clustering procedure (see Methods). *Right*: Activity dynamics, shown separately for detected, undetected, and reward trials. Note that cluster numbers and colors are assigned arbitrarily. Also note that in undetected trials (FAs) of the 1-frequency detection task both vagal opto and reward clusters are present, whereas in the psychophysics task the vagal opto cluster is absent. (f) Similarity of activity manifold cluster dynamics (cosine similarity). Bars show mean ± SEM across mice (n = 5); dots indicate individual animals. Higher values indicate greater similarity; lower values indicate greater separation. Comparisons: detected (CR) vs. undetected (FA), p = 0.04, Cohen’s d:-1.3; reward (hit) vs. undetected (FA), p = 0.42, Cohen’s d:-0.11; reward (hit) vs. detected (CR), p = 0.1 (paired 1 tail t-test), Cohen’s d: -0.8. (g) Schematic of bi-lateral InsCtx inhibition using stGtACR2 during non-invasive optogenetic activation of NG-OxtR^+^ neurons. (h) Schematic of the partial psychophysics task with optogenetic inhibition of the InsCtx. InsCtx inhibition occurred in a subset of trials for 5 seconds before the odor cue begins. In vagal opto trials, InsCtx inhibition began 1 sec before vagal stimulation and continued throughout its 4 sec duration (total of 5 seconds), ending before odor presentation. Vagal optogenetic stimulation included medium, low, and very low frequencies (per mouse, levels selected from that mouse’s psychometric curve; see Methods). Trials were randomly interleaved. (i) Example lick raster from one mouse for each trial type without (*left*) or with (*right*) InsCtx optogenetic inhibition. Each block of 50 trials (rows) indicates a different trial type, as labeled on the *right*: No opto, very low, low, and medium. Pink bars: vagal opto stimulation time window. Each gray tick indicates a lick. Trials were randomly interleaved and are presented here separately for clarity. (j) *Left*: average percentage of correct rejection trials across all vagal opto frequencies (very low, low, medium), without (gray) and with (purple) InsCtx optogenetic inhibition p = 0.002 (paired t-test), Cohen’s d = 2.42. Bars show mean ± SEM across mice (n=6). Dots indicate individual animals. *Right*: Percent of correct rejection for each vagal opto frequency without (gray) and with (purple) InsCtx inhibition: very low p = 0.0009 (paired t-test), Cohen’s d = 2.86, low p = 0.007 (paired t-test), Cohen’s d = 1.81, medium p = 0.013 (paired t-test), Cohen’s d = 1.54. (k) Percent of correct rejections in the single-frequency VagalOpto→NoReward task without (gray) or with (purple) InsCtx optogenetic inhibition. Bars show mean ± SEM across mice; dots indicate individual animals. *Left*: medium vagal opto stimulation frequency p = 0.61 (paired t-test, n=5). *Middle*: low vagal opto stimulation frequency p = 0.61 (paired t-test, n=5). *Right*: very low vagal opto stimulation frequency p = 0.59 (paired t-test, n=3). (l) Percent of hit trials (i.e., licking correctly and receiving a reward) with vs. without InsCtx optogenetic inhibition, in the psychophysics task (n=6 mice, p = 0.03, Effect size (rank-biserial correlation): r = -0.86, Wilcoxon test) and in the 1-frequency VagalOpto→NoReward task at medium (p = 0.47, Wilcoxon test, n=5 mice), low (p = 0.81, Wilcoxon test, n=5 mice) or very low (p = 0.75, Wilcoxon test, n=3 mice) vagal opto stimulation frequency. (m-o) Broad pharmacological inhibition of InsCtx during the psychophysics task (m,n) and the 1-frequency detection task (o). (m) Psychophysics task. *Left*: Average percent of correct rejection trials across all vagal opto frequencies (very low, low, medium), without (gray) and with (purple) InsCtx pharmacological inhibition p = 0.03 (one tailed paired t-test), Cohen’s d = -1.42. Bars show mean ± SEM across mice (n=4). Dots indicate individual animals. *Right*: percent of correct rejection for each vagal opto frequency without (gray) and with (purple) InsCtx pharmacological inhibition: very low p = 0.12 (one tailed paired t-test), low p = 0.049 (one tailed paired t-test), Cohen’s d = 1.19, medium p = 0.01 (one tailed paired t-test) Cohen’s d = 2.21. (n) Psychophysics task. Average percent of hit trials across all no opto trials without (white) and with (purple) InsCtx pharmacological inhibition p = 0.85 (paired t-test). Bars show mean ± SEM across mice (n=4). Dots indicate individual animals. (o) 1-frequency detection task. *Left*: percent of correct rejections during vagal opto trials p = 0.009 (one tailed paired t-test), Cohen’s d = 1.7. *Right*: percent of hits during no opto trials, p = 0.41 (paired t-test). Bars show mean ± SEM across mice; dots indicate individual animals (n=6).

We wondered whether decoding at the population level could discriminate detected vs. undetected vagal optogenetic stimulation trials in both the psychophysics task and the 1-frequency detection task. To test this, we trained two different population decoders on the vagal optogenetic stimulation time-window, while eliminating trial history effects by balancing after-reward and not-after-reward trials between conditions (see Methods). Consistent with the single neuron results, we found that we could readily decode detected vs. undetected trials in the psychophysics task. In contrast, we could not perform such successful population decoding in the 1-frequency detection task (**Fig. 3d; Extended Data Fig. 4f**). Importantly, we could accurately decode the presence or absence of vagal optogenetic stimulation in both tasks similarly (reward trials vs. vagal opto trials; **Extended Data Fig. 4g**). Decoding detected vs. undetected trials in the psychophysics task could potentially be explained by decoding of strong (usually detected) vs. weak (usually undetected) optogenetic stimuli. To test this, we limited the decoder to a subset of frequencies with a comparable number of detected and undetected trials (see Methods) and found similar effects (**Extended Data Fig. 4h**). This showed that decoding detected vs. undetected vagal optogenetic stimulation in the psychophysics task was not due to stimulus intensity.

To go beyond binary population decoding, we sought to explore InsCtx population activity pattern dynamics using an unsupervised approach. We used our previously established method for characterizing the InsCtx population activity manifold (i.e., the repertoire of population activity patterns) and dynamics within it^68^. We used non-linear dimensionality reduction, followed by automated clustering, which allowed us to describe activity dynamics as transitions between different subspaces (clusters) of the activity manifold (**Fig. 3e**). Using this approach, we found that in the 1-frequency detection task, undetected trials (false alarms) appeared as a simple combination of vagal optogenetic stimulation and reward anticipation (correct rejections and hits, respectively; **Fig. 3e, top row**). Remarkably, in the psychophysics task, undetected trials (false alarms) did not have a clear representation of the vagal optogenetic stimulus, regardless of stimulus frequency. Thus, it appeared to reflect reward anticipation, rather than reflecting the actual optogenetic stimulus (**Fig. 3e, bottom row**). We quantified this across mice using cluster dynamics similarity (**Fig. 3f**). This analysis showed that detected and undetected trials were less similar in the psychophysics task as compared to the 1-frequency detection task (**Fig. 3f, left**). Undetected trials (false alarms) resembled reward trials in both the 1-frequency detection and psychophysics tasks, likely reflecting (incorrect) reward anticipation in both tasks (**Fig. 3f, middle**). These results are consistent with the single-neuron choice probability analysis and the population decoding results above.

The different representations of detected vs. undetected stimuli in the different tasks suggest that InsCtx may contribute differently to internal sensory perception depending on task demands. To test this, we used optogenetic manipulations to alter InsCtx activity in a temporally precise manner during vagal optogenetic stimulation. Our rationale was that mild increases or decreases in InsCtx neural activity during the vagal optogenetic stimulus would bias detection, particularly if the stimulus was near perceptual threshold. However, our pilot experiments showed that even mild optogenetic excitation of InsCtx could cause seizures, similar to reports in the adjacent piriform cortex^66^, and we therefore focused on optogenetic inhibition. We expressed the light-gated chloride channel, stGtACR2^100^, in excitatory InsCtx neurons by injecting AAV1-CaMKIIa-stGtACR2-FusionRed, and verified the efficacy of inhibition using bulk fiber photometry of axonal projections of excitatory InsCtx neurons (**Extended Data Fig. 4i**). The magnitude of inhibition was qualitatively comparable to natural fluctuations in InsCtx activity, but was highly reliable across repetitions and time-locked to light onset and offset (**Extended Data Fig. 4i, right**). We quantified AAV infection and estimated that 47±3% (mean±STD) of neurons expressed stGtACR2.

We delivered precisely timed InsCtx optogenetic inhibition in the psychophysics task, time-locked to the vagal optogenetic stimulus period in 50% of vagal optogenetic trials and 20% of rewarded trials, randomly interleaved (see Methods). InsCtx inhibition (5 sec) started 1 sec before the vagal optogenetic stimulus epoch and ended together with the vagal optogenetic stimulus right before the odor cue. To have sufficient trial numbers to compare trials with vs. without optogenetic inhibition, we used 3 different vagal optogenetic frequencies instead of 7 (**Fig. 3g-h**). To avoid floor or ceiling effects (0% or 100% false alarms), we chose 3 frequencies for each mouse: ‘medium’, ‘low’, and ‘very low’ (typically 5 Hz, 3 Hz, and 1 Hz, respectively; see Methods). Optogenetic inhibition increased the number of detected trials across all three frequencies, which was seen as a reduction in the false alarm rate (**Fig. 3i-j**). In contrast, optogenetic inhibition of InsCtx during performance of 1-frequency detection tasks using only one frequency at a time (either ‘medium’, ‘low, or ‘very low’) had no effect on performance (**Fig. 3k**). Thus, even for very weak, near-threshold stimulus, the effect of partial inhibition depended on the variable multi-stimulus context causing higher perceptual uncertainty. It was not observed when the same weak stimulus was presented within a reliable 1-frequency context. Moreover, behavioral responses in the non-optogenetics trials (i.e., Hits) were also reduced by InsCtx inhibition in the psychophysics task, but not in the 1-frequency detection task, suggesting that mice mistakenly reported detecting a stimulus in the psychophysics task during no-stimulus trials (**Fig. 3l**). This suggests that partial InsCtx inhibition was sufficient to bias detection to the “stimulus present” report.

Importantly, we further tested whether the optogenetic inhibition effects were specific to the psychophysics task by also examining other behavioral contexts. In an odor-only Reward/NoReward task (no vagal optogenetics), pre-cue inhibition during what would have been the vagal optogenetic stimulation window, had no detectable impact (**Extended Data Fig. 4j**). This suggests that InsCtx inhibition did not affect odor cue discrimination and did not exert a general motivational effect. In the vagal optogenetic detection task at the maximum frequency (15 Hz), inhibition did not have significant effects in both VagalOpto→Reward (vagal opto→licking) and VagalOpto→NoReward (vagal opto→lick suppression) versions of the task (**Extended Data Fig. 4k**). This suggests that the InsCtx inhibition effects were not due to inhibition of licking nor to a general effect on motivation.

The InsCtx optogenetic inhibition results suggest that changing its activity patterns with partial inhibition is sufficient to bias detection. We therefore wondered whether a relatively mild manipulation of network activity targeting a small subset of neurons would be sufficient to cause similar effects. We performed an in-silico perturbation analysis on our InsCtx imaging data. We ranked neurons according to their activity during the vagal optogenetic stimulation window on detected trials (correct rejections) and selected the 20% most suppressed neurons for perturbation, leaving the remaining 80% unperturbed. We determined this proportion in an exploratory analysis, finding that decoding accuracy plateaued once the top 20% most suppressed neurons were included (**Extended Data Fig. 4l**). We note that suppressed neurons have similar sensory response properties to the vagal optogenetic sensory stimulus as excited neurons (e.g., Sigmoidal and Gaussian; **Extended Data Fig. 3g-h**). We trained a decoder on unperturbed population activity to distinguish detected vagal optogenetic trials (correct rejections) from no-vagal-opto trials (hits), and then tested how artificially reducing neuronal activity during the vagal optogenetic window of undetected trials (false alarms) affected their classification. This shifted undetected trials toward the detected class and improved their decoding, which quantitatively matched the improved behavioral detection with optogenetic inhibition across different frequencies (**Extended Data Fig. 4m,n**). The correspondence with behavior was strongest at moderate inhibition strength (**Extended Data Fig. 4m,n**) and was reduced with stronger inhibition (**Extended Data Fig. 4o,p**, see Methods). In contrast, adding moderate inhibition to the entire population significantly reduced decoding, as did applying moderate inhibition to the 20% most activated neurons (**Extended Data Fig. 4q,r**). These results suggest that optogenetic inhibition potentially modified activity patterns, but did not cause a broad silencing of InsCtx activity.

To test whether broad inhibition would have different behavioral effects that are distinct from partial optogenetic inhibition, we pharmacologically inhibited InsCtx using GABA-A and GABA-B receptor agonists^101^ (**Extended Data Fig. 4s**). In the psychophysics task, pharmacological inhibition reduced the detection of the vagal optogenetic stimulus, as evident by reduced correct rejection rates (**Fig. 3m**) and unchanged high hit rates (**Fig. 3n**). Moreover, unlike the task-specific effects of optogenetic inhibition, pharmacological inhibition produced a similar effect in both the psychophysics and the 1-frequency detection tasks (**Fig. 3m-o**). Taken together, these experiments suggest that partial optogenetic inhibition biases report under conditions of higher perceptual uncertainty in the psychophysics task. In contrast, the effects of broader InsCtx pharmacological inhibition across both tasks indicate a more general requirement for InsCtx in the behavioral report of internal signals.

### Distinct stable representations of internal mechanosensations and rewards

Reward expectation and consumption are prominent features of InsCtx activity^60,78,76,65,66,68^. We therefore next compared the representations of these two salient aspects of our behavioral tasks – vagal optogenetic stimulation and reward (**Fig. 4a**). At the single neuron level, we used the auROC-based analysis of single-neuron responses to identify neurons responsive to each event relative to baseline with either activation or suppression (see Methods). We found that most InsCtx neurons responded to either vagal optogenetic stimulation or reward, with some responding to both (**Fig. 4b; Extended Data Fig. 5a**). In the co-responsive neurons, we examined the relationship between opto and reward responses. Due to the potential contribution of opto-inhibited neurons described above, we restricted our analysis to neurons that were suppressed during the vagal optogenetic stimulation window on detected trials. We found a significant inverse relationship in every mouse, with consistently negative Pearson correlation coefficients across animals (mean r = −0.22, range = −0.36 to −0.16; all mice p ≤ 0.004; n = 5 mice; **Extended Data Fig. 5b**). This relationship supports the idea that vagal signals are embedded within a heterogeneous InsCtx population that also represents other sensory events, sometimes with opposing activity patterns.

**Figure 4:**
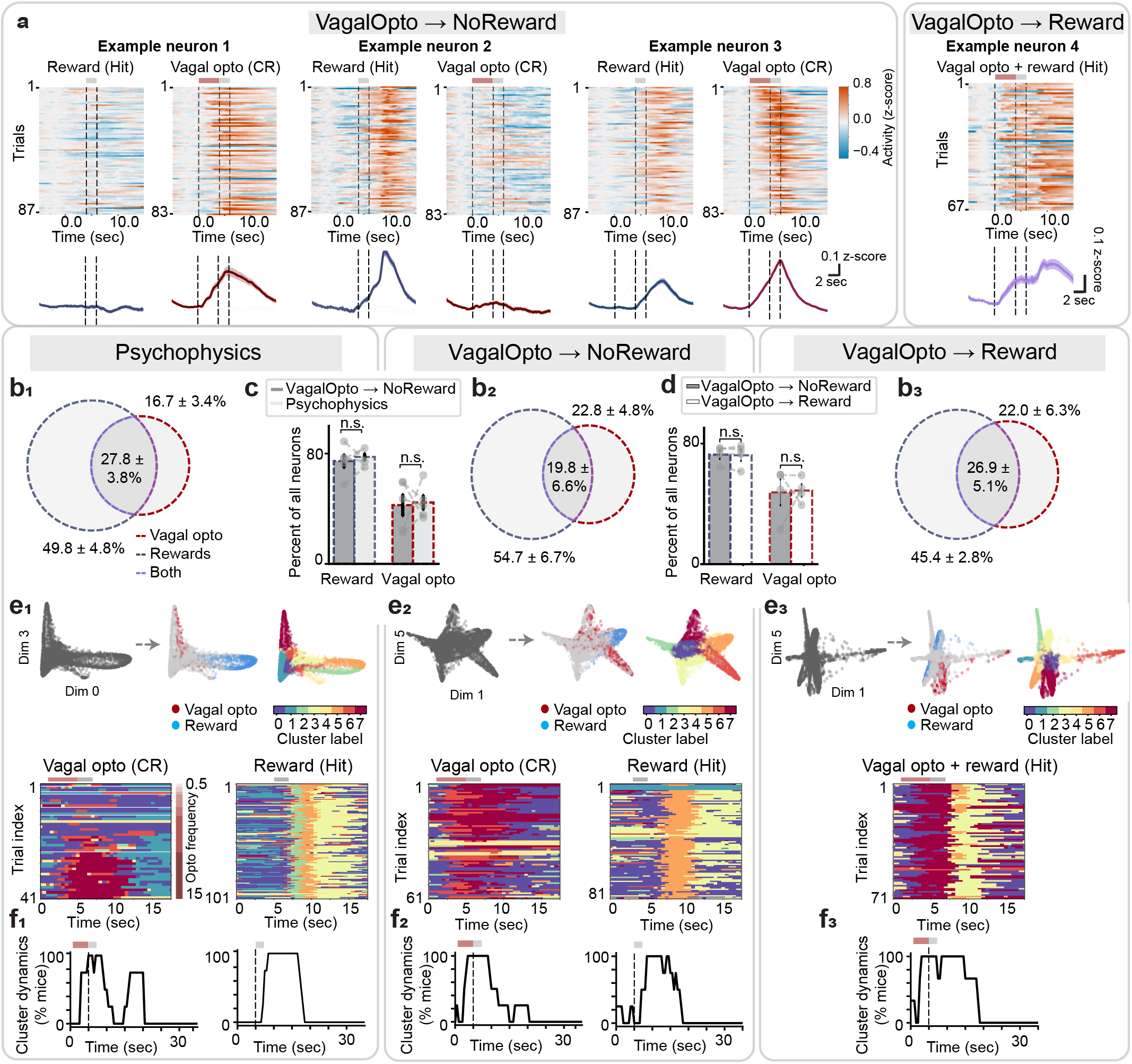
InsCtx representations of internal mechanosensations and rewards. (a) Example single-neuron responses. *Top*: Heatmaps in each subpanel show trial-by-trial activity aligned to vagal optogenetic stimulation. Pink bar marks the vagal opto stimulation period; gray bar marks the odor cue period. *Bottom*: average responses across trials (mean ± SEM). (b) Fraction of neurons significantly responsive to vagal opto stimulation only, reward only, or both, out of all recorded neurons per mouse. Values are mean ± SEM across mice (n = 5). (c) Comparison of psychophysics vs. 1-frequency detection tasks. Bars: fraction of recorded neurons per mouse that were significantly responsive to reward or to vagal optogenetic stimulation (mean ± SEM across mice, n = 5); dots indicate individual mice. Task effect: Reward p = 0.63, vagal opto p = 0.81; Wilcoxon signed-rank tests. n.s.: not significant. (d) Same as ‘c’, but comparing the VagalOpto→NoReward and VagalOpto→Reward tasks. Mean ± SEM across mice (n = 5); dots indicate individual mice. Reward p = 0.72, vagal opto p = 0.9; Paired t-tests. n.s.: not significant. (e) Representative planes of dimensionality-reduced population activity over time for psychophysics (e_1_), VagalOpto→NoReward (e_2_), and VagalOpto→Reward (e_3_) tasks. Each point is the population activity pattern in a 0.5 s bin. Colors indicate manifold-cluster assignment. Pink bar: vagal opto stimulation; gray bar: odor cue. Cluster numbers and colors are assigned arbitrarily. Note summation of vagal opto and reward clusters in the VagalOpto→Reward task (e_3_). (f) Activity manifold cluster dynamics over time for psychophysics (f_1_), VagalOpto→NoReward (f_2_), and VagalOpto→Reward (f_3_) tasks. Curves show the fraction of mice for each timepoint in which activity occupied at least one cluster (see Methods). Values are mean ± SEM across mice (n = 4). Pink bars: vagal opto stimulation; gray bars: odor cue.

Focusing specifically on Gaussian and Sigmoidal neurons, we found that they were also responsive to reward (**Extended Data Fig. 5c**). Interestingly, the fraction of neurons responding to vagal optogenetic mechanosensory stimulation and reward were quite similar between the psychophysics task (**Fig. 4b_1_**) and the 1-frequency VagalOpto→NoReward detection task using 15 Hz stimulation (**Fig. 4b_2_**, **Fig. 4c**). Additionally, the fraction of neurons responding similarly or oppositely (e.g., excitation to one and suppression to the other) to vagal optogenetic mechanosensory stimulation and reward were also similar across the two tasks (**Fig. 4b_1_-b_2_**, **Extended Data Fig. 5a**). However, fewer neurons responded overall to the 15 Hz stimulation frequency during the psychophysics task, suggesting dynamic shifts in frequency tuning between the tasks (**Extended Data Fig. 5d-e**). At the population level, the single-trial representations of vagal optogenetic mechanosensory stimulation and reward (i.e., correct rejections and hits) were similarly separable across both tasks (**Fig. 4e_1_-e_2_; Fig. 4 f_1_-f_2_**).

We wondered whether the difference in representations of vagal optogenetic stimulation and reward were due to differences in the task contingencies. That is, perhaps the optogenetic stimulus acquired a learned “no reward” or “lick inhibition” response that is naturally distinct from the lick-reward response. To test this, we trained mice on the reversed VagalOpto→Reward task (**Fig. 1o**) and imaged their InsCtx activity. The fraction of neurons responding to vagal optogenetic mechanosensory stimulation and reward were similar between the VagalOpto→NoReward and VagalOpto→Reward tasks (**Fig. 4b_2_-b_3_; Fig. 4d; Extended Data Fig. 5a**). We found similar separability of vagal optogenetic mechanosensory stimulation and reward representations in the VagalOpto→Reward task such that optogenetic hit trials now appeared as a simple sequential combination of distinct vagal optogenetics and reward representations (**Fig. 4e_3_; Fig. 4f_3_**). These results further support the conclusion that the responses to vagal optogenetic mechanosensory stimulation reflect in part their sensory features, and are largely independent of task contingencies and subsequent behavioral responses. We further tested whether the distinct representations of vagal optogenetic mechanosensory stimulation and reward would be spatially organized in InsCtx. Although we observed a small yet significant spatial organization of these distinct representations in some mice, it was not readily apparent visually. Additionally, this was not consistent across the different behavioral tasks (e.g., 4/5 showed clustering for the VagalOpto→NoReward task, but 1/5 for the psychophysics task; **Extended Data Fig. 5f,g**).

In summary, our results demonstrate that InsCtx representations of internal sensory stimuli and their perceptual report are distinct. Internal sensory representations are stable regardless of behavioral context or contingencies, whereas perceptual representations are only apparent in a difficult behavioral context with high uncertainty and play a causal role for perception in this context.

## Discussion

We developed a behavioral task in which mice report detecting non-invasive optogenetic activation of vagal mechanosensory neurons, establishing quantitative interoceptive psychophysics. This new approach paves the way for further mechanistic studies of internal perception in animal models. Combining this approach with cellular-resolution imaging and manipulations, we found that representations of sensory stimuli were consistent across different tasks, whereas perceptual reports were only encoded during a more difficult psychophysics task. Manipulations of insular cortex activity revealed that it is generally required for behavioral reports of internal mechanosensory signals in both task contexts. Furthermore, InsCtx plays an additional role in supporting perception only under conditions of uncertainty in the psychophysics task.

Based on these findings, we propose a framework in which InsCtx transforms graded vagal sensory representations into behavioral reports according to the reliability of the available sensory evidence, often referred to as “precision” in Bayesian and predictive coding frameworks^8,102–104^. Within this framework, sensory tuning reflects the representation of visceral sensory signals in insular populations, whereas choice probability-related representations only in the psychophysics task reflects trial-by-trial variation in how this sensory evidence is read out for behavioral report under perceptual uncertainty. Accordingly, manipulation of insular cortex activity with partial optogenetic inhibition biases report only in the psychophysics task. In contrast, broader pharmacological inhibition abolishes stimulus-dependent behavioral responses in both tasks. Our results reveal a cellular representation of interoceptive stimuli, and suggest flexible engagement of the interoceptive cortex depending on cognitive demands.

Human studies of interoception and specifically of interoceptive awareness, have advanced our understanding of these processes, as well as revealed potential neural mechanisms^3,6,16,105,106^. However, because such studies usually relied on the ability to track naturally fluctuating stimuli such as heartbeats, they lacked the ability to directly manipulate specific interoceptive sensory stimuli in a quantitative controlled manner. This constraint has hindered direct tests of how specific interoceptive stimuli are represented in the brain. Recent pioneering studies in humans using ingestible vibrating capsules and pressure-controlled breathing apparatuses have begun to perform such controlled studies of interoception^24,83,107^. Complementing these human studies with animal model studies that utilize powerful cellular-level neural recordings and manipulations, could provide detailed mechanistic insights into the neural basis of interoception^23^.

Animal models of interoception have mostly lacked the ability to capture perception of internal sensations through behavioral report^23^. Inspired by the recent quantitative psychophysics approaches to interoception in humans^107–106^, we developed a complementary approach in mice that enabled us to leverage the technological advantages of genetically and optically accessible mouse models. Our quantitative approach to interoception in mouse models opens up exciting opportunities to build specific bridges between work in animal models and in humans, which are much needed in the field of interoception and brain-body interactions^23,65,102^. These bridges now allow direct comparisons and bidirectional influences in parallel work using analogous protocols for conscious report of interoceptive sensations across humans and animal models. For example, future work using brief optogenetic manipulations of genetically identified baroreceptors to manipulate blood pressure^110^, cardiac pacemakers to control heart rate^85^, or lung mechanosensors to control breathing^34^, could establish different behavioral tasks in which mice report detecting such interoceptive signals. These would then allow comparisons with related work in humans on cardiovascular and breathing-related interoception.

The mid-posterior interoceptive InsCtx has been previously implicated in interoception awareness and attention^111^. Previous studies suggest that posterior insula regions primarily represent visceral, somatosensory, and interoceptive signals, while more anterior regions increasingly integrate these bodily signals with affective, motivational, contextual, and cognitive information^4,56,112^. Importantly, this organization is graded and bidirectional rather than strictly hierarchical, with overlapping connectivity and predictive-processing mechanisms across the InsCtx^8,113,114^. Indeed, predictive-processing accounts further propose that anterior agranular InsCtx regions generate interoceptive predictions and allostatic commands, while more posterior granular regions represent incoming sensory evidence and prediction errors, making processing fundamentally bidirectional rather than purely bottom-up^8^.

Previous neuroimaging work in humans has suggested that consciously focusing attention on interoceptive signals engages InsCtx^57,64^. Moreover, patients with different psychiatric conditions show changes in mid-InsCtx activation during interoception tasks, suggesting that InsCtx-based bodily representations are important for emotional regulation, and could be used as a neural marker of psychopathology^115^. We found that mid-posterior InsCtx represented vagal stimulus presence and reward-related information in both the psychophysics and 1-frequency detection tasks. In contrast, perception of internal mechanosensations was represented only during a difficult psychophysics task (where variable stimulus strength reduced the reliability of sensory evidence and made stimulus presence uncertain), but not during a simple detection task with one stimulus intensity, even a weak one. This suggests that InsCtx is not simply the primary detector of interoceptive inputs. Rather, it is an interoceptive readout system that converts graded visceral sensory representations into reportable perceptual decisions based on the reliability of the sensory evidence. Interestingly, the finding that partial optogenetic inhibition of InsCtx enhances detection of internal mechanosensations in the psychophysics task, while leaving performance in the 1-frequency detection tasks unaffected, suggests that the behavioral effect depends on the demands of perceptual judgment. Difficult psychophysical decisions may require task-relevant vagal evidence to be extracted from a heterogeneous InsCtx population that also represents spontaneous bodily-state fluctuations and other sensory signals. We speculate that partial optogenetic inhibition may alter the balance among these population components, thereby changing how InsCtx activity in response to ambiguous stimuli is mapped onto a perceptual report, biasing its behavior to the “stimulus-present” report.

Aberrant interoception plays a central role in pathological conditions such as eating disorders, anxiety, depression, and drug addiction^1,3,6,7,83,116^, yet the underlying etiology and neural mechanisms are poorly understood. More precise and controlled animal models offer the possibility to close these gaps. For example, does decreased interoceptive sensitivity reflect compromised peripheral afferent sensing (e.g., of gut mechanosensations for satiety) or aberrant central processing? If central processing is implicated, what are the neural circuits involved – brainstem sensing circuits, affective processing circuits, or action planning and executive function circuits? New mouse models, such as the one we present here, are ideally suited to provide detailed answers to these questions with cellular and molecular resolution. Such answers can then be used to rationally design molecular or circuit-level therapeutic interventions.

## Acknowledgements

We thank Assaf Deutsch for technical advice, and Daniel Goldian for technical assistance with electronics. We thank Laura Rupprecht and Diego Bohorquez for advice and assistance with vagal electrophysiology. We thank Diba Borgmann and Leonie Cabot for advice on nodose ganglia extraction and injection. We thank Matt Gardner and Guillermo Esber for advice on rodent behavioral training. We thank Henning Fenselau, Sahib Khalsa, Mark Andermann, Ilan Lampl, Sasha Devor, and members of the Livneh lab for comments on the manuscript and fruitful discussions. This work was supported by research grants from the European Research Council (ERC StG #101036145), Israel Science Foundation (ISF #860/21), the Peter and Patricia Gruber Awards, Sieratzki Institute for Advances in Neuroscience, Irene and Jared M. Drescher Center for Research on Mental and Emotional Health, Rolando Uziel, and the Center for New Scientists of the Weizmann Institute of Science.

## Methods

### Animals

We conducted all mouse care and experiments in accordance with protocols approved by the Institutional Animal Care and Use Committee at the Weizmann Institute of Science. We used 8-24-week-old mice for all experiments. We used the following mouse lines: OxtR-t2a-Cre (Jackson Laboratories stock #031303) and C57BL6/OlaHsd (Envigo). Sample sizes were chosen to reliably measure experimental parameters while keeping with standards in the relevant fields, and remaining in compliance with ethical guidelines to minimize the number of animals used. Experiments did not involve experimenter-blinding, but randomization was used with respect to trial order and data collection.

## Surgical procedures

### Brainstem injections

We anesthetized mice in an induction chamber with 3% isoflurane and maintained anesthesia at 1-1.5% via a nose cone. Once anesthetized, we secured each mouse in a stereotaxic frame (WPI). After removing fur, we made a midline scalp incision to expose the skull, and identified bregma as the reference point for stereotaxic targeting. Using a micro-drill (WPI), we performed craniotomies at the following coordinates for bilateral NTS injections: AP: -7.5, ML±0.4, DV: -5.2 to -4.8. We injected a retrograde AAV vector (retro-AAV-EF1a-DIO-ChRmine-mScarlet; plasmid #130668 from Addgene, produced by the Hebrew University ELSC Virus Core Facility) into the NTS of OxtR-Cre mice to retrogradely label NG-OxtR^+^ neurons. For controls, we either used OxtR-Cre mice, injected bilaterally with retro-AAV-CAG-FLEX-tdTomato (#28306, Addgene) into the NTS, or C57BL6 mice. Specifically for visualizing vagal innervation of the aortic arch, mice were injected with AAV-Retro6-CBA-mScarlet3 (produced by the Hebrew University ELSC Virus Core Facility). After the injections we sealed the craniotomy sites with bone wax and affixed a custom-designed head-post to the skull using Metabond (CGB Metabond® Quick Adhesive Cement System, # S380, Parkell).

Throughout all procedures, we continuously monitored respiration rate and anesthetic depth, including regular assessments of reflexive responses (e.g., toe-pinch). We lubricated the eyes with eye ointment (Duratears, Alcon) and provided analgesia using carprofen (5 mg/kg) and slow-release buprenorphine (3.25 mg/kg). We also closely monitored mice during recovery.

### Microprism assembly and surgery

Glass microprism implants were fabricated using 2 mm microprisms (#MCPH-2.0; Tower Optical) coated with aluminum along their hypotenuse. Microprisms were attached to a cover glass (#1 thickness), both along the hypotenuse (to prevent scratching of the reflective surface) and at the side of the prism that faces InsCtx, using Norland Optical Adhesive 81 cured using ultraviolet light. The glued prism assembly was then secured to a cannula using Kwik-Cast (WPI), allowing controlled insertion of the prism into the craniotomy.

Mice were anesthetized and prepared for surgery as described above. Using aseptic technique, a custom-made headpost was secured using CGB Metabond (Parkell). The mouse’s head was then rotated ∼70°. A ∼2.2×2.2 mm^2^ craniotomy was drilled over the left InsCtx, with the lower edge just above the squamosal plate. AAV8-hSyn-GCaMP7f (Addgene, #104488) mixed with FastGreen (Sigma; for visualization of viral spread) was injected directly into the exposed brain tissue at a depth of 0.2-0.6 mm in 4 adjacent locations (250 nL each) using a picospritzer with minimal injection pressure and long injection duration (PV850, WPI). The brain surface was kept hydrated throughout the procedure using saline-soaked gelfoam. The mouse’s head was then rotated back and the 2×2 mm^2^ microprism, attached to the cannula, was slowly lowered into the craniotomy until it made contact with the InsCtx. It was then advanced further while being slightly shifted medially (∼100 µm), ensuring that the lower edge of the microprism extended below the squamosal plate and contacted the medial wall of the craniotomy. Once properly positioned, the edges of the window were sealed to the skull with Vetbond (3M), followed by reinforcement with CGB Metabond (Parkell). The cannula was gently removed, and additional Metabond was applied to form a permanent seal. Mice were allowed to recover for at least six weeks prior to the start of behavioral training to allow for sufficient viral expression.

### InsCtx optogenetic inhibition

To validate the inhibition of excitatory InsCtx neurons using stGtACR2, we combined fiber photometry recordings of InsCtx → CeA projections with direct inhibition of InsCtx neurons in C57Bl6 mice. Specifically, we injected both AAV1-CaMKIIa-stGtACR2-FusionRed (#105666, Addgene) and AAV6-CaMKIIα-jGCaMP8m (#176751, Addgene) into the InsCtx (stGtACR2 bilaterally; jGCaMP8m to the right InsCtx) at the following coordinates: AP: 0 (bregma), DV: -4 to -3.8, ML: ±4mm (0.25 mm medial from the lateral edge of the skull). After the injection, we implanted optic fibers (Ø1.25 mm Ceramic Ferrule, 400 µm Core, 0.50NA; #R-FOC-BL400C-50NA, RWD Life Science) above the InsCtx (DV: -3.3mm) and another optic fiber (CFML15L05 -Fiber Optic Cannula, Ø1.25 mm SS Ferrule, Ø400 µm Core, 0.50 NA, L=5 mm, Thorlabs) above the CeA (AP:-1.4, ML:-2.6, DV:-4.5) for fiber photometry recordings. We secured all fibers and a custom-designed head post to the skull using Metabond (CGB Metabond® Quick Adhesive Cement System, # S380, Parkell).

Mice recovered and were then food restricted prior to any photometry recordings. We recorded axon fiber photometry signals of InsCtx excitatory neurons innervating the CeA using a pyPhotometry board^117^ with dedicated optics, together with 465 nm and 405 nm LEDs (Doric Lenses). The session began with a 2 minute baseline period without inhibition. Then, the protocol included 10 trials each of 5 seconds InsCtx inhibition (∼1mW blue light). We used a 3 second ITI with an additional randomized delay of 0-1 seconds to prevent temporal predictability. We analyzed the data by first applying a low-pass filter to denoise the signal. To remove brief artifacts occurring at the onset and offset of inhibition bouts, we excluded points that deviated more than three STD from local mean (computed with a sliding window) and interpolated the missing values from neighboring samples. We corrected photobleaching by fitting an exponential decay function to the signal and subtracting it^117^. We then removed motion-related artifacts by linearly fitting the isosbestic control signal to the GCaMP signal and subtracting the fitted motion component. Finally, we Z-scored the signal using the mean and STD of the baseline data.

For inhibition of excitatory InsCtx neurons during interoceptive detection tasks (described below), we used OxtR-Cre mice. We first performed brainstem injections of retro-AAV-EF1a-DIO-ChRmine-mScarlet (as described above) to express ChRmine in NG-OxtR^+^ neurons. During the same surgery, we bilaterally injected AAV1-CaMKIIa-stGtACR2-FusionRed (#105666, Addgene) into the InsCtx at the following coordinates: AP: 0 (bregma), DV: -4 to -3.8, ML: ∼±4mm (0.25 mm medial from the lateral edge of the skull). Following the injections, we implanted optic fibers (Ø1.25 mm Ceramic Ferrule, 400 µm Core, 0.50NA; #R-FOC-BL400C-50NA, RWD Life Science) above the InsCtx (DV: -3.3mm) and secured them with Metabond (CGB Metabond® Quick Adhesive Cement System, # S380, Parkell). Finally, we affixed a custom-designed head-post to the skull using Metabond. Throughout all procedures, we monitored anesthetic depth, reflexes, applied eye lubrication, and administered analgesia as described above.

### InsCtx pharmacological inhibition

For pharmacological inhibition of the InsCtx during the interoceptive detection tasks described below, we used OxtR-Cre mice. We first performed brainstem injections of retro-AAV-EF1a-DIO-ChRmine-mScarlet, as described above, to express ChRmine in NG-OxtR^+^ neurons. During the same surgery, we bilaterally implanted guide cannulas (26G, 0.48-mm diameter, M3.5; RWD Life Science, 62003) above the InsCtx and inserted dummy cannulas (0.3-mm diameter, M3.5; RWD Life Science, 6210238MM). The guide cannulas were 2.5 mm long, and the dummy cannulas extended 0.2 mm beyond the guides, such that the dummy cannula tips reached a dorsoventral depth of -2.7 mm. We positioned the cannula at AP: 0 mm relative to bregma, DV: -2.7, and ML: ∼±4mm (0.25 mm medial from the lateral edge of the skull). For later pharmacological infusions, we used 4-mm infusion cannulas (0.3-mm diameter, M3.5; RWD Life Science 62203), which extended 1.5 mm beyond the tips of the 2.5-mm guide cannulas, resulting in a final infusion depth of 4 mm below the skull surface. We secured the guide cannulas to the skull using Metabond (CGB Metabond® Quick Adhesive Cement System, # S380, Parkell) and affixed a custom-designed head-post to the skull using the same adhesive cement. Throughout surgery, we monitored anesthetic depth and reflexes, applied ophthalmic lubricant and administered analgesia as described above.

Insular cortex pharmacological inhibition was performed by infusing mice with 0.3 µL of muscimol and baclofen solution (0.05 µg / µL and 0.02 µg / µL, respectively) or saline at a rate of 0.1 µL / min to both hemispheres. One minute after each infusion, the injector cannula was replaced with the dummy cannula. Behavioral testing started 15 min later, following previously established protocols^66,101^.

### Vagal electrophysiological recordings

To enable optogenetic stimulation of NG-OxtR^+^ neurons, we performed stereotaxic surgery as described above to induce ChRmine expression. We conducted vagal electrophysiology approximately three months after viral injection, aligning with the maximal expression time used in the behavioral experiments.

Vagal electrophysiology was conducted as previously described^118^. Briefly, we induced anesthesia in a chamber using 3% isoflurane and maintained it at 1-1.5% via a nose cone. We made a small midline incision at the ventral neck to expose the cervical vagus nerve. Using a mouse retractor set (InterFocus #18200-20), we gently separated the salivary glands and masseter muscles laterally, and the esophagus medially, to reveal the carotid artery and adjacent vagus nerve. After carefully lifting the nerve with a spinal cord hook (FST #10162-12) and clearing the surrounding connective tissue with fine forceps, we proceeded with electrode placement. Prior to surgery, we had soldered two platinum-iridium wires (WPI #PTT0110), fashioned into hook-shaped “cuff electrodes”, to thicker platinum-iridium leads (WPI #PPT0502). This assembly was connected to a headstage (A-M Systems Connector Pins #521200 and #521000) and mounted on a right-handed micromanipulator (WPI M3301R) for stable positioning. Once the setup was in place, we gently positioned the cuff electrodes around the exposed vagus nerve.

We amplified neural signals 1000x and band-pass filtered between 300Hz and 5kHz using a differential amplifier (A-M Systems Microelectrode Amplifier). We then digitized the signals with a USB-6001 DAQ (National Instruments) and acquired them in real-time using MATLAB Analog Input Recorder app (MathWorks).

Following stable baseline recording, we delivered optogenetic stimulation using a 630 nm LED (UHP-F-5-630 with LLG-5, Prizmatix) which emitted 102 mW/mm^2^ light pulses at 15 Hz (each pulse 6.67 msec) for 4 seconds to the exposed neck overlaying the nerve. We manually controlled the stimulation using an Arduino Uno. For small intestine distension, we inserted a 20G gavage needle through the pyloric sphincter into the proximal small intestine and secured it in place with a suture. We clamped the distal small intestine with a bulldog clamp to prevent fluid outflow. To induce distension, we infused saline though the gavage needle until the outer intestinal diameter increased approximately 1.5-fold relative to its pre-distention diameter, corresponding to the range of intestinal distension observed following refeeding (see below).

At the end of the experiment, we euthanized mice using sodium pentobarbital (200mg/ml, 0.1 ml per 10g body weight, IP), and collected the NG and brains for histological analysis, as described below.

### Vagal electrophysiological data analysis

We analyzed data using custom MATLAB scripts. First, we band-pass filtered the signals between 80-2000 Hz. We identified and removed breathing and cardiac artifacts based on their characteristic amplitude and frequency profiles. We then detected spikes by applying an amplitude threshold to the filtered signal and used 200 ms bins to calculate spike rates.

### Measurement of intestinal diameter following fasting and refeeding

To determine the extent of small intestine distension induced by food consumption, we divided mice into two groups (n = 4 mice per group). Before measurements, we habituated mice three times to an overnight fast, followed by free access to food. Following habituation, we fasted both groups overnight. We euthanized mice in the fasted group at the end of the overnight fast. We provided mice in the refed group with ad libitum access to food for 2 hours following the overnight fast and then euthanized them. Immediately after euthanasia, we exposed the gastrointestinal tract and used a caliper to measure the outer diameter at the widest point of the small intestine.

### ECG measurements

We performed stereotaxic viral injections to express ChRmine in NG-OxtR^+^ neurons, as described above. To ensure that mice could perceive and report the optogenetic stimulation, we selected animals that had previously completed both the interoceptive detection VagalOpto→NoReward task and VagalOpto→Reward task (see below). We then recorded ECG signals under anesthesia while optogenetically stimulating NG-OxtR^+^ neurons using the same protocol employed in the awake behavioral task (e.g., 4-second light pulses at 15 Hz; 102 mW/mm^2^, each pulse 6.67 msec).

Before recording, we connected three 27G stainless steel needles to platinum-iridium leads (WPI #PPT0502) and connected them to a differential amplifier headstage (A-M Systems Microelectrode Amplifier). We induced anesthesia with 3% isoflurane in an induction chamber and maintained it at 1-1.5% via a nose cone. Once the mice were fully anesthetized, we attached the needles to the paws: two for signal acquisition and one for ground. We sampled ECG signals at 2 kHz and applied a 200 Hz low-pass filter during acquisition.

### Nodose ganglia two-photon imaging

To image NG-OxtR^+^ neuronal activity, we used OxtR-Cre mice and injected retro-AAV-hSyn-GCaMP7f (#104488, Addgene) into the NTS to drive GCaMP7f expression with no genetic specificity in NG neurons. To specifically identify retrogradely labeled NG-OxtR^+^ neurons, we also injected Cre-dependent retro-AAV-CAG-FLEX-tdTomato (#28306, Addgene) into the NTS. This approach allowed us to image calcium dynamics across the NG population while distinguishing the OxtR^+^ subset based on tdTomato fluorescence.

4-6 weeks after the injections, we performed a two-photon calcium imaging experiment during intestinal balloon inflation to evoke mechanosensory responses. We anesthetized mice with an intraperitoneal injection of pentobarbital (100 mg/kg). After confirming full anesthesia, we removed fur from the ventral neck and upper abdomen. We made a small incision above the anatomical location of the stomach, gently retracted the overlying tissue, and identified the stomach and proximal duodenum. We carefully externalized the duodenum, made a small incision in the intestinal wall, and inserted a latex balloon (Braintree Scientific #73–3478) that we had pre-attached to a plastic gavage needle (Instech ITH-FTP-22-25-50). We secured the balloon with sutures, and intermittently irrigated the exposed tissues with sterile saline. For some mice, we used 2 adjacent balloons in the small intestine, inflated separately during the experiment. After balloon placement, we returned the intestine to the abdominal cavity and placed saline soaked gauze above, to avoid tissue dehydration. Following the abdominal procedure, we made a midline incision at the ventral neck to expose the cervical vagus nerve. Using a mouse retractor set (InterFocus #18200-20), we gently separated the salivary glands and masseter muscles laterally and the esophagus medially to expose the carotid artery and adjacent vagus nerve. We carefully lifted the nerve with a spinal cord hook (FST #10162-12) and removed surrounding connective tissue using fine forceps. We exposed the NG and positioned it on a microscope slide, covered with a coverslip for stabilization. We filled both the neck cavity and the space between the slide and the coverslip with sterile saline to maintain hydration.

We placed the mouse on a heating pad to maintain body temperature throughout the surgery and imaging, and positioned the two-photon objective vertically above the exposed nodose ganglion. We connected the balloon to a tube filled with water and attached it to a syringe mounted on a programmable syringe pump (KFTechnology #NE-1000). Intestinal mechano-stimulation was achieved by volume-controlled inflation of the implanted gastric balloons. Each stimulus lasted 50 seconds, including 10 seconds of inflation and 10 seconds of deflation.

We imaged NG activity using a two-photon microscope (Thorlabs Bergamo), with 620 nm excitation (Toptica FFUltra 620). We also collected red fluorescence emitted with 620 nm excitation to identify NG-OxtR^+^ neurons expressing tdTomato. Imaging was performed at a frame rate of 15 Hz.

### Direct ablation of NG-OxtR^+^ neurons

For specific ablation of NG-OxtR^+^ neurons using Caspase 3, we directly injected AAV6-flex-taCasp3-TEVp (plasmid #45580 from Addgene, produced by the Hebrew University ELSC Virus Core Facility) bilaterally into the NG. Briefly, we exposed the NG as described above, and used a pulled glass pipette to inject the viral solution directly into the ganglion. To aid visualization and confirm proper targeting, we mixed the virus with fast green dye (Sigma) prior to injection. These surgeries were performed in two separate procedures, aimed to improve recovery and overall survival rates. The first surgery included the injection of retro-AAV-EF1a-DIO-ChRmine-mScarlet into the NTS of OxtR-Cre mice and headpost implementation as described above. Additionally we directly injected of AAV6-flex-taCasp3-TEVp into the right NG. Following sufficient recovery (∼3-4 weeks after the first surgery), we performed the second surgery, where we injected AAV6-flex-taCasp3-TEVp into the left NG.

### InsCtx fiber photometry recordings with and without NG-OxtR^+^ ablation

For fiber photometry recordings of InsCtx population activity, we injected AAV8-hSyn-GCaMP7f (#104488-AAV8, Addgene) into the left InsCtx (AP: 0 mm relative to bregma; ML: approximately -4 mm, 0.25 mm medial to the lateral edge of the skull; DV: -4 to -3.8 mm). We implanted an optic fiber (400-µm core, 0.50 NA, 1.25-mm ceramic ferrule; R-FOC-BL400C-50NA, RWD Life Science) above the injection site at a depth of -3.3 mm. In all mice, we additionally injected retro-AAV-EF1a-DIO-ChRmine-mScarlet into the NTS, as described above, to express ChRmine in NG-OxtR^+^ neurons.

For recordings without NG-OxtR^+^ ablation, we allowed mice to recover and for sufficient viral expression before recording InsCtx activity during optogenetic activation of NG-OxtR^+^ neurons. Before recordings, we habituated mice to red-light illumination.

Each recording session consisted of eight trials of optogenetic stimulation. In each trial, we stimulated NG-OxtR^+^ neurons at 15 Hz for 4 s, using the same stimulation parameters as in the behavioral experiments described above. As a control for responses to red-light illumination itself, we performed an additional recording session in which we delivered the same light stimulation but directed the light away from the mouse’s neck, thereby preventing optogenetic stimulation of NG-OxtR^+^ neurons.

To examine InsCtx responses to the optogenetic stimulation following NG-OxtR^+^ ablation, we used two experimental approaches. In a subset of mice, we first performed photometry recordings with intact NG-OxtR^+^ neurons and subsequently performed bilateral NG-OxtR^+^ ablation using AAV6-flex-taCasp3-TEVp, as described above, allowing us to record from the same mice before and after ablation. In a separate subset of mice, during one surgical procedure, we injected retro-ChRmine into the NTS and GCaMP7f into the InsCtx, implanted the optical fiber above the InsCtx, and performed the first NG injection of AAV6-flex-taCasp3-TEVp. Following recovery, we performed the second injection of AAV6-flex-taCasp3-TEVp to the contralateral NG. Once mice were fully recovered from both procedures, we performed InsCtx photometry recordings using the same optogenetic stimulation paradigm.

After completion of the experiments, we collected the NG and brain tissue for histological analysis as described below. We assessed the extent of NG-OxtR^+^ neurons ablation and verified GCaMP7f expression and optical fiber placement in the InsCtx.

We analyzed InsCtx photometry responses across the three experimental conditions by aligning activity to the onset of optogenetic stimulation or red-light delivery. We low-pass filtered the photometry signal, corrected for photobleaching using a double-exponential fit, and z-scored each trial relative to the 5-s pre-stimulation baseline. For each mouse, we quantified the response as the mean z-scored activity during the 0-to 4-s stimulation period and compared responses across conditions.

### Aortic depressor nerve transection

We anesthetized mice with 3% isoflurane for induction and maintained anesthesia at 1.5% isoflurane throughout the surgical procedure. We placed mice in a supine position and exposed the cervical region as described above. We identified the vagus nerve and carotid artery and moved the carotid artery medially to facilitate visualization of the aortic depressor nerve^116,120^. We visually identified the aortic depressor nerve as a thin nerve running between the vagus nerve and carotid artery and transected it using fine surgical scissors. We then sutured the surgical incision and proceeded with brainstem injections as described above. We verified successful NG infection and aortic depressor nerve denervation post-mortem by observing labeling of neurons in the NG but without labeled axons in the aortic arch.

## Behavioral experiments

### Initial habituation

After a minimum of 2 weeks of post-surgical recovery, we food-restricted mice (or in a subset of experiments, water-restricted, as noted below) to 80-85% of their free-feeding body weight. Before behavioral training, we habituated animals to head-fixation by gradually increasing head-fixation durations from 10 minutes to 1 hour per session over the course of 2-3 days. If we observed any signs of stress, we immediately removed the animal and conducted additional head-fixation sessions until the animal showed no visible signs of stress. During the final head-fixation session, we provided sugar-water (600 mM sucrose solution, #4072.1000 Mallinckrodt Baker -Avantor) via a lickspout to train the mice to associate licking with a sugar-water reward (∼5 μL per drop). We tracked licking behavior using a custom capacitance-sensing lickspout based on an MPR121 capacitance sensor. We performed all behavioral training using MonkeyLogic2^121,122^ (https://monkeylogic.nimh.nih.gov/).

To habituate animals to the red light used during optogenetic activation, we directed a separate masking red light (Thorlabs M625F2 625 nm LED; ∼28 mW/mm^2^; 10Hz; 20% duty cycle) toward the mice. This light visually masked the optogenetic stimulation light throughout the experiment, turning on at the beginning of every trial and switching off during the inter-trial interval (ITI). After several days of training with the masking red light, we lightly anesthetized mice using isoflurane (as described above), and removed fur from the ventral neck using hair-removal cream (Veet). We removed the fur to ensure efficient penetration of light through the skin and underlying tissues for non-invasive optogenetic activation of vagal neurons. We verified that the light guide delivering red LED light for optogenetics (5 mm diameter light guide: LLG-5, LED: UHP-F-5-630, Prizmatix) did not heat up or damage the skin in the stimulation area. We also habituated mice to the presence of two light guides positioned beneath and lateral to the neck, gently touching the skin. In all experiments using optogenetic stimulation as a conditioning cue, we additionally habituated animals to a custom neck cover made of opaque fabric that enclosed both the neck and light guides, further minimizing red light leakage outside the stimulation area (e.g., to the eyes).

### Sucrose consumption test

Following habituation, we introduced an olfactory cue (Isoamyl Acetate, W205508, Sigma-Aldrich) using a custom-built olfactometer. We diluted the odor to 10% (volume per volume) in mineral oil solution and applied an additional 10% air dilution in the olfactometer, resulting in an overall 1% concentration. We first trained mice to associate the 2-second odor cue with the unconditional delivery of a sugar-water drop (Pavlovian reward, ∼5 μL). Once they consistently licked in anticipation of the odor cue and completed at least 200 trials, we increased the number of sugar-water drops delivered in each trial. The odor cue signaled the start of the trial, during which 3 Pavlovian sugar-water drops were delivered at 1-second intervals. If the mouse continued licking after these initial 3 drops, it could receive up to 7 additional “operant drops” – drops with delivery contingent upon licking. Mice performed 40 trials with a 30-second ITI. Once performance stabilized (typically after 2-5 sessions), we conducted a test session. During the test, in half of the trials (randomly selected), licks for drops number 5 to 10 triggered both sugar-water delivery and a 1-second non-invasive optogenetic activation of NG-OxtR^+^ neurons (15 Hz, 10% duty cycle, 102 mW/mm^2^) delivered via two light guides aimed at the ventral neck to illuminate the NG. Importantly, trials with or without optogenetic stimulation were indistinguishable to the mice prior to the onset of stimulation, as both used the same odor cue and rewards, and both included the presence of the masking light.

We counted licks during the ’optogenetic window" (defined as the earliest to last time of stimulation across trials) and compared them between optogenetic trials and control trials (without optogenetics) from the same session. We calculated a lick reduction index as:

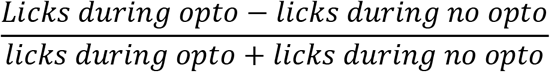

We used the same procedure for both food-restricted and water-restricted cohorts.

In an initial subset of mice (4/14 ChRmine, 1/5 control), we used a slightly different protocol without an olfactory cue. Each trial began with 3 Pavlovian sugar-water drops, followed by up to 17 operant drops (i.e., drops with delivery contingent upon licking) for a total of up to 20 drops per trial with a 30-secod ITI. In half of the trials (randomly selected), a 1-second of optogenetic stimulation was delivered during licking for the final 15 drops. We calculated the lick reduction index as described above. Although this protocol produced licking suppression in response to the optogenetic stimulation, mice were less consistent in recognizing trial onset without an olfactory cue, and the longer trial length reduced the number of completed trials. As a result, licking patterns were also less consistent across trials and across days. We therefore switched to the odor cued protocol described above for most mice.

### Head-fixed Conditioned Taste Preference (CTP)

For CTP experiments, we water-restricted mice. Following habituation, we trained mice to associate two distinct olfactory cues with two different non-caloric juice rewards. Ethyl Butyrate odor (E15701, Sigma-Aldrich) signaled access to artificial apple juice, and Isoamyl Acetate odor (W205508, Sigma-Aldrich) signaled artificial grape juice. We presented each odor for 2 seconds at the start of each trial. We artificially sweetened both Juices (Yachin) with 25 mM Acesulfame K (#04054-25G, Sigma-Aldrich), and allowed mice to earn up to 15 drops (∼5 µL per drop) per trial.

We delivered juices operantly: mice had to lick during a designated response window that began immediately after odor offset, with at least 1-second between drops. If a mouse paused licking, juice delivery stopped and resumed only if licking continued within the response window. We set the ITI to range from 30 seconds to 2 minutes, scaling it linearly with the number of drops received: more juice (i.e., more licking) resulted in longer ITIs.

We randomly interleaved trials at a 1:1 ratio. Once the mice performed the task reliably, we used the final training session as a “preference baseline” and calculated a lick preference index:

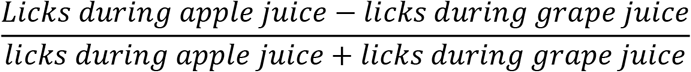

We identified each mouse’s preferred juice based on this index. Over the next three sessions (one per day), we paired the preferred juice with non-invasive optogenetic activation of NG-OxtR^+^ neurons. These trials began with the same 2-second odor cue and the initial three drops delivered operantly. From the fourth drop onward, we paired each lick that triggered a juice drop with 1 second of optogenetic stimulation. We continued the other trial type (the less preferred juice) as before, without stimulation.

After each session, we calculated a new preference index. On the final day, we conducted a test session with no optogenetic stimulation and calculated a final preference index to assess changes in juice preference due to the prior stimulation pairing.

### Interoceptive detection task – VagalOpto→NoReward

Following habituation, we trained mice to associate an olfactory cue (Ethyl Butyrate, E15701, Sigma-Aldrich) with the delivery of 1 drop of sugar-water reward. Each trial began with a 2-second odor presentation, followed by a 2-second licking window. If the mice licked during this window, it triggered the delivery of a sugar-water drop (∼5 µL). Once the mice consistently performed the operant licking behavior and completed at least 200 trials, we introduced a second trial type. In these new trials, we delivered a 4-second non-invasive optogenetic activation of NG-OxtR^+^ neurons before presenting the same odor cue. We directed the stimulation via two light guides aimed at the ventral neck (15 Hz, 10% duty cycle, 102 mW/mm^2^). Licking during the post-odor window in these trials was considered a “false alarm”, and was not rewarded with sugar-water (but also not explicitly punished). Each interoceptive detection session included 180-200 trials, with optogenetic and non-optogenetic trials randomly interleaved at a 1:1 ratio. To prevent visual detection of the red light, we used masking red light in all trials (∼28 mW/mm^2^; 10Hz; 20% duty cycle) that started 5 seconds before odor presentation and lasted until the beginning of the ITI in all trials. As described above, we additionally used opaque fabric neck covers to limit red light leakage outside the stimulation area (e.g., to the eyes).

We maintained the same light level for each individual mouse across all training and testing sessions. For most mice we used 102 mW/mm^2^, but in order to have a usable number of false alarms, for some mice with higher sensitivity, we used a lower light intensity of optogenetic stimulation (range: 15-102 mW/mm^2^). We used the same frequency and duty cycle for all mice.

For experiments involving body temperature measurements, we selected animals that demonstrated high performance in the interoceptive detection VagalOpto→NoReward task. On the day of temperature assessment, we ran mice in the same behavioral paradigm (∼180 trials with 50% optogenetic stimulation trials), and measured body temperature using a non-contact infrared thermometer (Beurer FT60) aimed at a part of the mouse’s back from which we removed hair to expose the skin. To ensure accuracy, we positioned the device at a 2-4 cm distance from the mouse’s dorsal surface, avoiding direct skin contact. We took temperature measurements during brief pauses between trials (during the ITI), minimizing disturbance to behavior.

### Training with lower-frequency optogenetic stimulation

To assess learning dynamics across days in the VagalOpto→NoReward task, we trained naïve mice using lower-frequency NG-OxtR^+^ optogenetic stimulation. We trained mice on the task, as described above, but used 1 Hz instead of 15 Hz stimulation. All other stimulus parameters remained the same. We selected 1 Hz because this stimulation frequency was near or below the behavioral detection threshold observed in the psychophysics experiments described above, allowing us to track the emergence of interoceptive detection across training. We trained mice with 1 Hz stimulation over multiple consecutive daily sessions. We quantified behavioral performance during learning of the 1-Hz stimulation across training days using Hit rate, false-alarm (FA) rate, and d′. We calculated Hit rate from no-opto trials and FA rate from 1-Hz opto trials, and calculated d′ from the corresponding Hit and FA rates using the Hautus correction.

### Temporal delay test – VagalOpto→NoReward

To determine the temporal window during which mice could detect optogenetic activation of NG-OxtR^+^ neurons, we first trained mice to proficiency in the interoceptive detection VagalOpto→NoReward task, as described above. Notably, in this task variant, there is no delay between optogenetic stimulation offset and odor cue onset. Once mice reached stable performance, we performed a delay test session in which we varied the interval between the end of the optogenetic stimulation and the onset of the odor cue. We kept the optogenetic stimulation parameters identical across trials (4 s, 15 Hz, 10% duty cycle, 102 mW mm^-2^) and randomly varied the delay between stimulation offset and odor onset between 0, 2, 4, 6, and 8 s. We randomly interleaved the different delay conditions at equal proportions. To maintain task engagement and provide sufficient opportunities to obtain rewards, 50% of trials consisted of NoOpto→Reward trials.

We assessed behavioral performance across delay conditions using Hit rate, false-alarm (FA) rate, and d′. We calculated Hit rate from no-opto trials and FA rate separately for opto trials with 0-, 2-, 4-, 6-, or 8-s delays. For each delay condition, we calculated d′ from the corresponding FA rate and the Hit rate measured in no-opto trials, using the Hautus correction to avoid infinite values at extreme response rates.

### Optogenetic stimulus duration test – VagalOpto→NoReward

To determine the minimal duration of NG-OxtR^+^ neuron activation necessary for robust interoceptive detection, we first trained mice to proficiency in the VagalOpto→NoReward task, as described above, with a 4 sec optogenetic stimulus. Once mice reached stable performance, we performed a stimulus duration test session. During this session, we randomly varied the duration of the optogenetic stimulus while maintaining the same stimulus frequency and duty cycle (15 Hz, 10% duty cycle) and presented the odor cue immediately after stimulation offset. Optogenetic stimuli consisted of either 60 pulses (4 s), 15 pulses (1 s), 4 pulses (∼0.26 s), or a single pulse. We randomly interleaved the four stimulus durations at equal proportions. To maintain task engagement and provide sufficient opportunities to obtain rewards, 50% of trials consisted of no-opto trials.

We assessed behavioral performance across stimulation conditions using Hit rate, false-alarm (FA) rate, and d′. We calculated Hit rate from no-opto trials and FA rate separately for opto trials with 60 pulses (4 s, 15 Hz), 15 pulses (1 s, 15 Hz), 4 pulses (15 Hz), or a single pulse. For each stimulation duration, we calculated d′ from the corresponding FA rate and the Hit rate measured in no-opto trials, using the Hautus correction to avoid infinite values at extreme response rates.

### Odor-cue generalization test

To determine whether mice could generalize interoceptive detection across different olfactory cues, we first trained mice to proficiency in the VagalOpto→NoReward task using the original trained odor, as described above. On the test day, mice initially performed 20 trials of the task using the trained odor. We then switched, without prior exposure or training, to a novel odor cue and continued the same task for an additional 180 trials. All other task parameters remained unchanged.

We assessed behavioral performance following the odor change by comparing overall Hit and false-alarm (FA) rates between the previously learned and new odor conditions. For each odor, we pooled trials with and without vagal opto stimulation. We calculated Hit rate and FA rate, and calculated d′ using the Hautus correction to avoid infinite values at extreme response rates.

### Training for reversal from the VagalOpto→NoReward task to the VagalOpto→Reward task

Once mice became proficient in the interoceptive detection VagalOpto→NoReward task (i.e., hit rate > 75% and false alarm rate < 25%), We transitioned them to the VagalOpto→Reward version of the task. In this version, we introduced a new olfactory cue (Isoamyl Acetate, W205508, Sigma-Aldrich) that signaled the availability of a sugar-water reward, but only when preceded by a 4-second non-invasive optogenetic activation of NG-OxtR^+^ neurons (15 Hz, 10% duty cycle, typically 102 mW/mm^2^, range: 15-102 mW/mm^2^, as described above). We began by running 10-20 training trials where we consistently paired the optogenetic stimulation with the odor cue and delivered a Pavlovian sugar-water reward. Then, we randomly interleaved two trial types: (1) opto-odor-reward trials, and (2) odor-only trials with no reward, regardless of licking behavior. Once the mice reliably licked only in response to trials with prior stimulation, we transitioned to a fully operant version of the task. In this version, we delivered a sugar-water reward only if they licked during the response window of trials where the odor cue was preceded by the optogenetic stimulation. False alarms were not explicitly punished.

### Within-session no-training reversal from the VagalOpto→NoReward task to the VagalOpto→Reward task

To assess how mice responded to an unexpected reversal of the learned interoceptive contingency, we first trained mice to proficiency in the VagalOpto→NoReward task, as described above. On the reversal test day, mice initially completed 20 trials under the previously learned VagalOpto→NoReward contingency. We then reversed the contingency within the same session, without prior training or explicit indication of the change, and continued the session under the VagalOpto→Reward contingency. Following the reversal, optogenetic stimulation of NG-OxtR^+^ neurons preceding the odor cue signaled reward availability, whereas odor-only trials were not rewarded. In contrast to the training VagalOpto→Reward paradigm described above, we did not provide Pavlovian pairing trials or any other training on the reversed contingency before the switch. All other stimulation and behavioral parameters remained as described above.

We assessed behavioral performance before and after contingency reversal by comparing overall Hit and false-alarm (FA) rates. We classified responses as Hits, Misses, correct rejections, or FAs according to the active stimulus-reward contingency. We pooled trials within the original or reversal phases and calculated Hit rate, FA rate, and d′ using the Hautus correction.

### Interoceptive psychophysics

Once mice became proficient in the interoceptive detection VagalOpto→NoReward task, we performed 1-2 days of the psychophysics task. We began each session with 20 trials that followed the same structure as the VagalOpto→NoReward paradigm described above. After this initial phase, we ran approximately 280 additional trials. In half of these, they received a Go cue (odor only), allowing them to earn a sugar-water reward if they licked during the appropriate time window, maintaining task engagement with 50% rewards. In the remaining trials, we paired non-invasive optogenetic activation of NG-OxtR^+^ neurons with the odor cue. We used one of seven frequencies (0.5, 1, 3, 5, 7.5, 10, 15 Hz; typically 102 mW/mm^2^, range: 15-102 mW/mm^2^, as described above, 10% duty cycle), all presented at equal probability. Licking during the 2-second window following odor presentation in these trials resulted in no reward and counted as a “false alarm”. We randomly interleaved all 8 trial types throughout the session. Before the psychophysics day, mice were trained on the VagalOpto→NoReward at 15 Hz opto stimulation frequency to achieve performance criterion. We did not train mice further on the psychophysics task, and only used it 1-2 times to probe their detection threshold.

### Behavioral tasks with InsCtx optogenetic inhibition

We expressed the inhibitory opsin stGtACR2 under the CaMKIIa promotor and used implanted optic fibers to deliver continues blue light for 5 sec (∼10 mW), selectively inhibiting InsCtx neurons. We specifically chose an optogenetic protein that responds to blue light rather than red light to avoid interference with the red light used for optogenetic vagal activation. All mice underwent several behavioral tasks as detailed below.

#### Interoceptive detection VagalOpto→NoReward and VagalOpto→Reward with InsCtx inhibition

Mice were initially trained to perform the VagalOpto→NoReward task before introducing InsCtx inhibition. In the inhibition experiment, optogenetic inhibition began 5 seconds before odor onset and terminated immediately prior to odor presentation. This protocol was applied to both trial types (odor-only and vagal optogenetic stimulation trials). To ensure balanced sampling, inhibition was included in 25% of the trials for each trial type, resulting in ∼50 inhibition trials and ∼150 non-inhibition trials per condition within a session. Later, the same mice were trained to perform the VagalOpto→Reward version of the task (as described above), and a similar procedure was performed for InsCtx inhibition.

A similar protocol was used for the 1-frequency detection VagalOpto→NoReward task with ‘medium’, ‘low’, and ‘very low’ vagal opto frequencies (see below). Mice were initially trained to perform the VagalOpto→NoReward task at 15 Hz and were then tested with the chosen frequency.

#### Partial psychophysics with InsCtx inhibition

For psychophysics with InsCtx inhibition, we selected three different stimulation frequencies for each mouse: ‘medium’, ‘low’, and ‘very low’. These frequencies were typically 5Hz, 3Hz, and 1Hz, respectively. To avoid floor or celling effects, if a mouse had below 5% false alarms at 5Hz we chose frequencies 3, 1, and 0.5 Hz. Conversely, if a mouse had above 65% false alarms at 1Hz, we chose frequencies 7.5, 5 and 3 Hz. Note that we used three frequencies instead of the full range of seven frequencies used for psychophysics task without InsCtx inhibition, to ensure a sufficient number of repetitions for each trial type with/without InsCtx inhibition.

In this task, 50% of trials were Go cue trials (odor only) without vagal optogenetic stimulation, allowing the mice to obtain a sugar-water reward by licking during the response window. In the remaining 50% of trials, we delivered vagal optogenetic stimulation at one of the three selected frequencies prior to the odor cue.

We randomly interleaved bilateral InsCtx inhibition across all trial types. In vagal stimulation trials, we initiated InsCtx inhibition 1 second before the onset of vagal optogenetic stimulation, maintained it throughout the 4-second stimulation period, and ended it immediately before odor presentation. We inhibited InsCtx activity in 50% of vagal stimulation trials, with a minimum of 50 inhibition trials per stimulation frequency to achieve enough trials for quantitative analysis. In odor-only rewarded trials without vagal opto stimulation (a total of 300 trials), we inhibited InsCtx activity 5 seconds before odor onset. We inhibited InsCtx in 60 trials so that we will not have too many trials with InsCtx inhibition, which may reduce the fraction of trials mice lick to receive reward.

#### Odor Go/NoGo control with InsCtx inhibition

We trained mice to associate two distinct olfactory cues (Isoamyl Acetate, W205508, Sigma-Aldrich; and Limonene, W263303, Sigma Aldrich), which they had not previously encountered. One odor cued the availability of a sugar-water reward, while the other signaled a ‘no-outcome’ (i.e., no reward). This task did not include vagal optogenetic stimulation. After several training sessions, once mice reliably discriminated between the two odors, we performed the inhibition experiments. InsCtx inhibition began 5 seconds before odor onset and terminated immediately before odor presentation. We randomly interleaved inhibition in 25% of the trials for each odor type, resulting in approximately 50 trials with InsCtx inhibition and 150 trials without inhibition per condition.

### Behavioral tasks with InsCtx pharmacological inhibition

Mice were implanted with bi-lateral guide cannulas to allow injection of drugs directly into the InsCtx as described above. Pharmacological inhibition was performed by infusing mice with 0.3 µL of muscimol and baclofen solution (0.05 µg / µL and 0.02 µg / µL, respectively) or saline at a rate of 0.1 µL / min in both hemispheres. One minute after each infusion, the injector cannula was replaced with the dummy cannula. Behavioral testing started 15 min later, following previously established protocols^66,101^.

To assess the effect of pharmacological InsCtx inhibition on interoceptive detection, we first trained mice to proficiency in the VagalOpto→NoReward task, as described above. Before pharmacological testing, we habituated mice to manipulation of the dummy injector and implanted cannulas to minimize the associated stress during drug administration.

Each mouse underwent both a pharmacological inhibition session and a control session on different days. On the inhibition day, we bilaterally infused muscimol and baclofen into the InsCtx as described above. On the control day, we followed the same procedure but bilaterally infused saline instead. Following infusion, mice completed 10 no-opto trials to obtain sugar-water rewards and engage in the task. Mice then completed 20 or 40 trials of the standard 1-frequency VagalOpto→NoReward task, followed by 180 trials of a partial psychophysics test using three vagal optogenetic stimulation frequencies (1, 3, and 5 Hz). Half of the trials consisted of no-opto trials, whereas the remaining 50% consisted of vagal optogenetic stimulation trials, with all trial types randomly interleaved throughout the session. We used the same trial structure and stimulation parameters on the muscimol and baclofen and saline control days.

For verification of injection sites, following completion of all experiments, we bilaterally injected 0.3 µL of 0.1% (w/v) cholera toxin subunit B conjugated to Alexa Fluor 555 (CTB-55; Thermo Fisher Scientific, C34776) through the implanted cannulas. Following the injections, we transcranially perfused the mice and collected and processed the brain and NG tissues as described below.

### Two-photon imaging of InsCtx during vagal opto behavioral tasks

Two-photon imaging was performed using a resonant-scanning two-photon microscope (Thorlabs Bergamo) at 15 frames per second, via a 10x 0.5 NA air objective (TL10X-2P, Thorlabs) at 1.5 zoom, yielding a 606×606 µm field of view (512×512 pixels). We used a 620 nm fixed-wavelength laser for GCaMP imaging (Toptica FemtoFiber Ultra 620). All imaged fields of view (FOV) were at a depth of 60-200 mm below the pial surface, with laser power at 620 nm of 15-80 mW at the front aperture of the objective (power at the sample was likely substantially less due to partial transmission via the microprism). Mice were imaged in consecutive 30-minute runs. Imaging depth was adjusted in between runs (every 30 min) to account for slow drift in the Z plane (< 7 µm).

Both the masking red light and the vagal optogenetic activation were performed between imaging frames. To control light on/off timing, we used an Arduino Uno, which switched states in response to commands sent via the MonkeyLogic2 program.

We note that the anatomical parcellation of the mouse insular cortex into anterior/mid/posterior divisions is quite inconsistent across the literature. The part of the insular cortex we imaged is sometimes referred to as “mid”, and sometimes as “posterior”. We refer to our imaging location as “mid-posterior insular cortex” as a compromise between these different parcellations. Across multiple analyses, we did not find any clear functional-anatomical organization within mouse or across mice (**Extended Data Fig. 3m,n, Extended Data Fig. 4d,e, Extended Data Fig. 5f,g**). Therefore, we did not find a clear distinction between the parts of the mid and posterior insula we imaged.

## Histology

### Brain tissue collection and processing

For most experiments, we euthanized mice via IP injection of sodium pentobarbital (200 mg/kg; 0.1ml per 10g body weight), followed by intracardial perfusion. Briefly, we began by making a small incision in the right atrium and inserted a needle into the left ventricle. We flushed the circulatory system with PBS to clear the blood, then perfused with 10% formalin to fix the tissues. After perfusion, we removed the heads and post-fixed them in 10% formalin for 24-48 hours before extracting the NG (as described below) and the brain.

Before slicing, we cryoprotected brains in 30% sucrose for 24-48 hours until the tissue sank. We then sectioned the brains coronally at 50 μm using a sliding microtome (Leica), mounted selected sections onto glass slides, and applied DAPI-containing mounting medium (VECTASHIELD® Antifade Mounting Medium with DAPI H-1200-10, Vector Laboratories). We placed coverslips and imaged the slides using a slide scanner (VS120-S6-W, Virtual Slide Microscope System, Olympus) at 10X magnification, and viewed using OlyVIA V4.1 software (Olympus).

### Nodose ganglia collection and processing

As described above, we euthanized mice and, in most cases, perfused them to ensure thorough fixation of the head. After fixation, we positioned the head to expose the ventral neck and made a midline incision in the skin using scissors. We carefully removed the salivary glands and excised the digastric muscle to expose the glossopharyngeal nerve, which we used as an anatomical landmark for locating the vagus nerve. We then cleared additional muscle and connective tissue to access the carotid sheath, which encloses the carotid artery, vagus nerve, and associated connective tissue. We gently separated the vagus nerve from the artery and traced rostrally toward the base of the skull. This dissection led us to the jugular foramen, where we identified the NG as a small bulb-like structure. Using fine forceps, we carefully isolated and freed the NG. After extraction, we mounted the ganglia onto glass slides, applied mounting medium (VECTASHIELD® Mounting Medium, #101068-042, Vector Laboratories), and placed coverslips. We then imaged the slides using a fluorescence microscope (AxioLab, Zeiss).

### Intestine collection and processing

To visualize axonal terminals of NG-OxtR^+^ neurons in the intestine, we injected retrograde AAV encoding ChRmine-mScarlet into the brainstem of OxtR-Cre mice, as described above. After allowing 2-3 months of expression, we euthanized the mice with sodium pentobarbital (200 mg/kg; 0.1ml per 10g body weight) and transcardially perfused them to ensure thorough fixation. We carefully removed the entire intestine while minimizing disruption to surrounding connective tissues and placed the tissue in 10% formalin for approximately one week of post-fixation. Following fixation, we dissected the intestine and cryopreserved it in 30% sucrose solution for at least 48 hours. We then sectioned the tissue at 100 μm thickness using a sliding microtome (Leica), mounted on glass slides, and applied mounting medium (VECTASHIELD® Mounting Medium, # 101068-042, Vector Laboratories) before placing coverslips. We imaged the sections using a two-photon microscope (Thorlabs Bergamo), with excitation at 620 nm for green tissue autofluorescence (Toptica FemtoFiber Ultra 620) and 1050 nm for mScarlet fluorescence (Toptica FemtoFiber Ultra 1050). We acquired Images across multiple z-planes and processed them using ImageJ to generate maximum intensity projections.

### Aortic arch collection and processing

We collected aortic arches using a dissection procedure adapted from^123^. Briefly, we euthanized mice with sodium pentobarbital (200 mg/ml; 0.1 ml per 10 g body weight, intraperitoneally) and placed them in a supine position under a stereomicroscope. We opened the thoracic cavity and removed the rib cage to expose the heart and major blood vessels. We carefully removed the surrounding connective and adipose tissues to expose the aortic arch and its major branches. We excised the aortic arch and flushed the vessel with saline to remove residual blood. We then fixed the tissue in 10% formalin for approximately 1-2 hours. Following fixation, we mounted the aortic arch on a microscope slide in mounting medium (VECTASHIELD® Mounting Medium, # 101068-042, Vector Laboratories), covered it with a coverslip and imaged the whole-mounted tissue by fluorescence microscope (AxioLab, Zeiss).

### Analysis of stGtACR2-FusionRed expression in the InsCtx

To assess stGtACR2-FusionRed expression in the InsCtx, coronal brain sections were imaged from all mice included in the optogenetic inhibition experiments, as described above. For each mouse, all sections in which the tip of the optic fiber tract was visible were included in the analysis. Images were analyzed using ImageJ. The boundaries of the InsCtx were manually delineated in each section. The membrane localization of stGtACR2-FusionRed created large patches with tissue fluorescence in which individual somata were not clearly separated. Therefore, to obtain an estimation of the fraction of cells expressing stGtACR2-FusionRed, we examined the areas of tissue fluorescence and manually outlined them. We then quantified the percentage of stGtACR2-FusionRed fluorescence areas out of the total InsCtx area per mouse across all examined tissue sections. Group data are presented as the mean ± SEM across mice.

## Statistical Analysis

Throughout this study, we employed an automated test selection procedure to ensure appropriate statistical methods were applied based on data characteristics. For paired comparisons (e.g., comparing the same animals across conditions), we first assessed normality of the differences using the Shapiro-Wilk test (α = 0.05). When differences were normally distributed without outliers, we applied paired t-tests; otherwise, we used Wilcoxon signed-rank tests. For independent comparisons between groups, normality was assessed separately in each group using the Shapiro-Wilk test, and variance homogeneity was tested using Levene’s test. When both groups were normally distributed, we selected independent t-tests (for equal variances) or Welch’s t-tests (for unequal variances). When one or both groups deviated from normality, we applied the non-parametric Mann-Whitney U test. Importantly, when multiple related comparisons were performed within the same analysis (e.g., comparing different percentile thresholds between conditions), if any single comparison in that family failed normality assumptions, we applied the non-parametric test to all comparisons within that family to ensure consistency and fairness. For all comparisons, we computed effect sizes (Cohen’s d for parametric tests, rank-biserial correlation for non-parametric tests) and 65% confidence intervals for mean differences using bootstrap resampling (10,000 iterations). Statistical significance was assessed at α = 0.05, with FDR correction applied when multiple comparisons were performed within the same family of tests.

## Data analysis

### Two-photon imaging data preprocessing

Two-photon imaging data were first processed using Suite2p^124^ for image registration. Data were then denoised using DeepInterpolation^125^. The denoised data was then used for ROI extraction using Suite2p. Extracted ROI masks were then manually inspected to ensure typical soma morphology, appropriate size, and normal activity patterns. Neuropil contamination was corrected by subtracting 70% of the surrounding neuropil signal from each cellular fluorescence trace (i.e., multiplying neuropil values by 0.7), followed by addition of the median neuropil value to preserve baseline fluorescence levels. Motion artifacts were removed using custom-written code in Python, which is based on artifact removal procedures in human EEG recordings^126,127^. This method involves a multi-step approach including z-score normalization for each 30 min imaging run, high-pass filtering to eliminate low-frequency drift, followed by principal component analysis (PCA) and independent component analysis (ICA) to identify and exclude components significantly correlated with x-y motion displacement traces (from the Suite2p registration output). The cleaned signals were then inverse-transformed to recover baseline-corrected fluorescence values in the original scale. Fluorescence signals were converted to relative fluorescence changes (ΔF/F) using a running baseline approach, where the 10th percentile fluorescence value within a 4.5 min sliding window served as the baseline fluorescence for each neuron. This method provided robust normalization against slow baseline drift while preserving rapid calcium transients. Synchronization with behavioral parameters (e.g., odor and opto onset/offset, licking etc.) was achieved by aligning imaging frames with stimulus timing codes recorded via ThorSync (Thorlabs) at 6 kHz, with behavioral signals downsampled to match the imaging frame rate. Continuous imaging data were segmented into individual trials based on the timing of Monkeylogic2 behavioral codes, enabling trial-based analysis of neural responses aligned to task events. All preprocessing steps preserved the temporal structure of the data while removing technical artifacts, enabling subsequent analysis of task-related neuronal activity patterns.

#### Data normalization

Neural activity (ΔF/F) was z-scored per neuron to account for differences in baseline fluorescence and dynamic range. For each neuron, ΔF/F values were z-scored by subtracting the session mean and dividing by the session standard deviation. Normalization was applied to the continuous trace before trial segmentation to ensure consistent baseline statistics across trial types.

### Behavioral psychometric curve analysis

Behavioral performance in the vagal optogenetic detection task was quantified by constructing psychometric curves relating stimulus intensity to detection probability. The initial 20 trials at the beginning of each session, which were only 0 or 15 Hz, were excluded from analysis to allow for task engagement and stabilization of behavioral performance. For each stimulation frequency, the detection rate was calculated as the proportion of trials in which the animal produced a lick response. These empirical detection rates were then fitted with a four-parameter sigmoid function of the form:

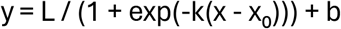

where x represents the stimulation frequency, y represents the detection rate, L is the dynamic range, x₀ is the inflection point (perceptual threshold), k controls the slope, and b is the baseline offset. Parameter estimation was performed using nonlinear least-squares optimization with initial parameter guesses derived from the data. The quality of the fit was assessed by computing the coefficient of determination (R²). To assess psychometric fit quality, response labels were randomly permuted across trials while stimulation frequencies were kept fixed. For each of 100 shuffles per mouse, the psychometric curve was refitted and its R² was calculated. The real R² was then compared with the mean shuffled R². The perceptual threshold for each animal was defined as the inflection point (x₀) of the fitted sigmoid, corresponding to the stimulation frequency at which the fitted detection probability reached the midpoint of its dynamic range. Uncertainty in the perceptual threshold estimates was assessed using bootstrap resampling. For each mouse, trials were resampled with replacement within each stimulation frequency while preserving the original number of trials per frequency. The psychometric function was refitted and the perceptual threshold was recalculated for each of 1,000 bootstrap iterations. The 65% confidence interval was defined by the 2.5th and 67.5th percentiles of the resulting bootstrap threshold distribution. For visualization, detection rates were converted to percentages and plotted as a function of optogenetic stimulation frequency along with the fitted sigmoid curve. When analyzing multiple animals, individual psychometric curves were overlaid and the average R² value across animals was computed and reported along with the standard error of the mean to summarize detection performance consistency across the cohort.

### Neural response detection and neurometric analysis (Gaussian and Sigmoid)

#### Neural response detection

To quantify neural responses to vagal optogenetic stimulation, we employed area under the receiver operating characteristic curve (auROC) analysis. For each neuron and trial, we compared neural activity during a baseline period (from 1 s before to 1 s after trial onset, before vagal optogenetic stimulation) with activity during the stimulation period (3-5 s after trial onset). We defined 100 threshold criteria spanning the 0.1st to 66.6th percentile of each neuron’s activity range. For each threshold, we calculated the probability that neural activity exceeded the threshold during baseline and stimulation periods separately. The auROC value was computed using these probability distributions, where values >0.5 indicate excited responses (increased activity during stimulation), values <0.5 indicate suppressed responses (decreased activity), and values ≈0.5 indicate no response.

#### Statistical significance testing of auROC values

To determine significant responses, we performed permutation testing for each neuron on a trial-by-trial basis. For each trial, we shuffled baseline and stimulation labels 500 times to create a null distribution of auROC values under the assumption of no stimulus-evoked response. Significance thresholds were set at the 5th and 65th percentiles of this null distribution (α = 0.05). Only trials with auROC values exceeding these thresholds were considered significant responses and retained for further analysis.

#### Neurometric curve construction

To characterize frequency-dependent neural responses, we constructed neurometric curves for each neuron. For each stimulation frequency, we calculated the fraction of trials showing significant responses. Excited responses were defined as trials with auROC > 0.5, while suppressed responses were defined as trials with auROC < 0.5. This yielded neurometric curves plotting response fraction against stimulation frequency, analogous to the behavioral psychometric curves. We fitted both Sigmoid and Gaussian functions to each neuron’s neurometric curve using least-squares optimization.

The Sigmoid function was defined as:

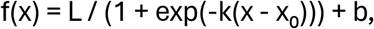

where L is the amplitude, x₀ is the inflection point, k is the slope, and b is the baseline. The Gaussian function was defined as:

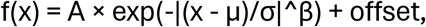

where A is the amplitude, μ is the center, σ is the width, β is the shape parameter, and offset is the baseline.

For each neuron, we fit both models separately to excited responses and suppressed responses. The goodness-of-fit was assessed using the coefficient of determination (R²). For each neuron, we selected the model (Sigmoid or Gaussian) and response type (excited or suppressed) that yielded the highest R². Neurons were classified as "Sigmoid neurons" if the Sigmoid model provided the best fit (indicating monotonic frequency-response relationships) or "Gaussian neurons" if the Gaussian model provided the best fit (indicating tuned frequency preferences). An additional permutation test was performed by shuffling condition labels 1000 times and refitting the curves to determine if the observed fit was higher than chance. Only neurons with fits that were better than chance were finally considered as “Sigmoidal” or “Gaussian”.

Based on their mean auROC values across all trials, neurons were further classified as excited (mean auROC > 0.5) or suppressed (mean auROC < 0.5). This yielded four categories: Sigmoid excited, Sigmoid suppressed, Gaussian excited, and Gaussian suppressed neurons. Only neurons with significant neurometric fits (R² above threshold and significant permutation test) were included in subsequent analyses. We used these response categories and indices in the choice probability analysis and single-neuron response classification, described in the corresponding sections below.

#### Population-level analysis and comparison to behavior

For Sigmoid neurons meeting inclusion criteria, we extracted the half-maximum threshold, defined as the stimulation frequency at which the fitted curve reached fifty percent of its dynamic range (i.e., midpoint between minimum and maximum response). For Gaussian neurons meeting inclusion criteria, we extracted the peak frequency (μ parameter), representing the stimulation frequency that elicited the maximal response probability. These neural thresholds were aggregated across neurons within each animal and compared to the behavioral detection threshold derived from psychometric curve fitting. To visualize the distribution of neural thresholds within individual animals, we created kernel density estimate plots, showing the probability density of Sigmoid half-maximum thresholds and Gaussian peak frequencies across the stimulation frequency range. To characterize population-level distributions, we pooled all neurons across animals and binned them into discrete frequency categories. To quantify the variability of neural and behavioral thresholds across animals, we computed the coefficient of variation (CV = standard deviation / mean × 100%) for behavioral detection thresholds, Sigmoid half-maximum thresholds, and Gaussian peak frequencies, providing a normalized measure of inter-animal diversity in threshold distributions.

### Neural population decoding and comparison to behavior

We trained support vector machine (SVM) classifiers to decode stimulus presence from neural activity patterns and compare them to behavior. We wanted to train the classifiers on sufficiently high frequencies that could activate neurons, while ensuring that the animal could not reliably detect them behaviorally. Specifically, for each animal, we selected a training frequency based on its behavioral detection threshold, selecting the stimulation frequency that was 2 frequency levels below the threshold when available (e.g., for a behavioral threshold of 7.5 Hz, we trained the classifier on 3 Hz), or the lowest frequency below threshold if only one existed (e.g., for a behavioral threshold of 1 Hz, we trained the classifier on 0.5 Hz). Neural activity patterns during the vagal opto stimulation period were extracted for all trials at the training frequency, and a binary SVM classifier was trained to discriminate stimulus-present from stimulus-absent conditions. Classifier performance was evaluated using repeated 70/30 holdout cross-validation. In each of 500 iterations, 70% of the eligible trials were randomly assigned to the training set, while the remaining 30% were reserved for testing and were not used during classifier training. Classification accuracy was averaged across iterations. The trained classifier was then tested on its ability to decode stimulus presence across all stimulation frequencies, generating a frequency-dependent decoding accuracy curve analogous to the psychometric curve. For cell-subset analyses, the classifier was retrained after excluding Gaussian neurons, Sigmoidal neurons, or a randomly selected set of neurons matched in size to the average number of Gaussian and Sigmoidal neurons.

To quantify the difference between neural coding and behavioral performance, we computed the difference between classifier accuracy and behavioral detection rate specifically at the training frequency for each animal (Δ = classifier accuracy − behavioral detection rate, expressed as percentages). This Δ metric represents information present in the neural population that was not translated into behavioral choices. For group-level analysis, we plotted this accuracy gap as a function of each animal’s behavioral threshold, examining whether the magnitude of subthreshold neural information varied systematically with detection sensitivity. Representative individual animal data were visualized by overlaying the psychometric curve with the full classifier decoding curve to illustrate the relationship between behavioral performance and neural population decoding across the entire frequency range.

### Choice Probability Analysis

#### Neural discrimination between correct rejection (detected) and false alarm (undetected) trials

To quantify choice-related modulation under identical sensory input, we computed choice probability (CP)^67^ by comparing neural activity between correct rejection (CR, detected) and false alarm (FA, undetected) trials. For each neuron, mean ΔF/F activity during the vagal optogenetic stimulation period (0–4 s after onset) was extracted separately for CR and FA trials. To correct for unequal trial numbers, we used bootstrap resampling: on each of 500 iterations, we randomly subsampled without replacement from the larger condition to match the smaller one and computed the auROC between CR and FA activity. The median auROC across iterations was taken as the neuron’s CP value.

Following established neural-discrimination methods, auROC values were obtained by sweeping 100 threshold criteria across the 0.1st–66.6th percentile of each neuron’s activity range. For each threshold, we calculated the probability that neural activity exceeded the threshold during CR and FA trials, respectively. The auROC value was then computed from these probability distributions, where values of CP > 0.5 indicate higher activity on CR (detected) trials, CP < 0.5 indicate higher activity on FA (undetected) trials, and CP ≈ 0.5 indicate no discrimination.

#### Trial-history controls

To minimize potential trial-history effects, we applied balanced sampling when sufficient data were available. Trials were labeled as after reward or not after reward. During each bootstrap iteration, CR and FA trials were sampled so that the proportions of these trial history categories were matched between conditions, ensuring that observed CP differences reflected true choice-related modulation rather than trial-history effects.

#### CR-CR control analysis for establishing null distribution thresholds for significance of choice probability values

To determine objective statistical thresholds for identifying neurons with CP values significantly above chance, we performed a control analysis separately for each animal. Correct rejection (CR) trials were randomly split into two groups of equal size while preserving trial history proportions, as described above. One randomly selected subset of CR trials was labeled as "pseudo-FA" and the other as "pseudo-CR". We then computed CP values for all neurons using this CR-CR split, repeating the random split and CP calculation procedure 500 times to generate robust estimates. Because both groups consisted of identical trial types (CR trials), any observed CP values in this control analysis would reflect noise, measurement variability, and random fluctuations rather than true behavioral discrimination. This procedure generated an animal-specific null distribution of CP values representing the expected range of discrimination (false positive rate) in the absence of true signal differences.

For each animal, we then calculated percentile-based thresholds from their CR-CR control CP distribution to define statistical significance criteria (false positive rate). Neurons were separated into excited (CP ≥ 0.5) and suppressed (CP < 0.5) populations based on their CP values. For excited neurons, we computed the 66.5th, 66th, 67.5th, and 65th percentiles of the animal’s control distribution, representing the top 0.5%, 1%, 2.5%, and 5% of that animal’s null distribution. For suppressed neurons, we computed the 0.5th, 1st, 2.5th, and 5th percentiles, representing the bottom 0.5%, 1%, 2.5%, and 5% of the animal’s null distribution. These animal-specific thresholds define the extreme values that would occur by chance in the absence of true discrimination for that particular animal.

### Neural Decoding with Sliding-Window Classifiers

To assess the temporal dynamics of neural population coding during task performance, we implemented a sliding-window classification approach using both linear support vector machine (SVM) and Naive Bayes classifiers. The continuous normalized signals were divided into individual trial epochs, with each trial including a 4-second pre-trial baseline period. To account for potential sequential dependencies, we explicitly tracked trial history, labeling each trial as either occurring immediately after a reward or not. Classifiers were trained to discriminate between different behavioral outcomes: correct rejections (CR) versus false alarms (FA), hits versus CR, and hits versus FA. We first trained classifiers on all trials regardless of frequency or trial count. Additionally, to account for potential behavioral differences between detected and undetected trials at different frequencies (i.e., no detected trials at the lowest frequency or no undetected trials at the highest frequency), we only used trials from optogenetic stimulation frequencies in which both classified classes (i.e., hit/CR/FA) contained at least 2 trials at that frequency (before accounting for trial history). For classifier training, we employed a stratified bootstrap procedure to balance trial history between the two classes being compared. Specifically, on each iteration, we identified the minority and majority classes, then equalized the proportion of "after-reward" and "not-after-reward" trials in both classes by choosing to the minimum available count for each history category. This ensured that classifier performance reflected true differences in neural activity patterns rather than confounds from trial reward history. As an additional safeguard, if either class ended up with fewer than 2 trials after stratified bootstrap balancing, that bootstrap iteration was skipped to ensure valid 70/30 train-test splits. Balanced trial sets were then split into training (70%) and test (30%) sets separately for each class, with strict enforcement that no trial appeared in both training and test sets. Classifiers were trained on neural population activity during a fixed temporal window (1 to 4 seconds after vagal opto stimulation onset). For each trial, we extracted a population (feature) vector consisting of the mean activity of each neuron during this training window. Linear SVMs and Naive Bayes classifiers were fit to these features using the training set. Classification accuracy was then evaluated dynamically across time using a sliding window approach: we computed mean neural activity in consecutive 0.3-second windows (step size = 1 frame, ∼67 msec at 15 Hz), spanning from trial onset to 17 seconds post-onset. At each time point, the trained classifier predicted trial identity based on the population activity within that window, and accuracy was computed by comparing classifier predictions to true labels on the test set. This procedure was repeated for 500 bootstrap iterations, and accuracy curves were averaged to obtain mean ± SEM traces. For group-level analysis, individual-animal classifier results were aggregated across mice.

### In Silico Neural Perturbation Analysis

To assess the causal contribution of specific neuronal populations to behavioral discrimination, we implemented an in silico perturbation approach in which we selectively altered the activity of neurons suppressed by vagal opto and evaluated the impact on classifier performance. For each animal, we selected a single "training frequency" for the perturbation analysis by training separate classifiers at each available optogenetic stimulation frequency (without perturbation), evaluating their accuracy curves across all test frequencies, and selecting the training frequency whose classifier accuracy curve most closely matched the shape of the behavioral psychometric curve. This ensured that perturbations targeted the neural population most relevant to perceptual discrimination as reflected in behavior. We first ranked all neurons by their median activity during the stimulation window (0 to 4 seconds after vagal opto stimulation onset) on correct rejection (CR) trials at the selected training frequency. Neurons were sorted in ascending order, such that the most suppressed neurons (those with the most negative median z-scored ΔF/F) were ranked first. To identify the critical proportion of suppressed neurons required for effective discrimination, we performed an initial exploratory analysis testing perturbations on cumulative subsets of the ranked neurons in 10% increments (top 10%, top 20%, top 30%, …, top 100%), where each subset included all neurons from the most suppressed up to that percentile. This analysis revealed that decoding accuracy plateaued when the top 20% most-suppressed neurons were included, indicating that this subset was sufficient to reach maximal perturbation effects (**Extended Data Fig. 4l**). Based on this result, we focused subsequent perturbation experiments on three neuron populations: (1) the top 20% most-suppressed neurons, (2) the 20% most-excited neurons as a control, and (3) all neurons. To quantify the typical magnitude of inhibition, we computed the median activity across all negative-activity responses from CR trials at the training frequency, which served as a reference inhibition value. We applied perturbations exclusively to false alarm (FA) trials by adding a scaled version of the reference inhibition value to the activity of selected neurons. Specifically, for a given neuron subset and a given scaling multiplier (0, 2, or 5), we modified each selected neuron’s activity on FA trials by adding (reference inhibition × multiplier). Multiplier values of 0 represented no perturbation (original activity). Critically, perturbations were applied only to the stimulation-window activity used as classifier features; all other aspects of the data remained unchanged. For each combination of neuron subset and multiplier, we trained a linear SVM classifier on unperturbed data from the training frequency CR trials versus no vagal opto hit trials, using mean activity during the stimulation window as features. The classifier was then tested on perturbed FA trials at all presented optogenetic frequencies, as well as on unperturbed CR and hit trials, using a cross-validation 70/30 train-test split with 500 bootstrap iterations. To prevent train-test contamination, CR trials at the training frequency were split such that training samples were drawn exclusively from the CR pool, while the test set comprised the remaining CR trials plus all FA trials (which received perturbations). Classification accuracy was computed separately for each frequency, yielding accuracy curves as a function of stimulation frequency for each perturbation condition. We quantified the perturbation effect as Δ accuracy = (accuracy with perturbation) − (accuracy at original activity, multiplier = 0). For group-level analysis, we aggregated Δ accuracy across animals for matched perturbation conditions.

Because animals had different vagal optogenetic stimulation frequencies in the behavioral InsCtx inhibition experiment, we also selected three corresponding stimulation frequencies for this analysis: “medium,” “low,” and “very low” (see further explanation in the “Partial psychophysics with InsCtx inhibition” section).

### Single-Neuron Response Classification

To identify neurons responding to specific behavioral events (i.e., vagal optogenetic stimulation and reward delivery), we performed event-specific auROC analyses on z-scored neuronal activity. For each event type, we defined temporal windows for event-related and baseline activity. The baseline window was consistent across all events (0 to 1 second after trial onset). Event windows were: (1) vagal stimulation: 1 second after vagal optogenetic onset until optogenetic offset (3-second window); (2) reward delivery: 2 seconds following reward onset; (3) lick onset: from lick onset to lick offset.

#### Adaptive sliding-window auROC for vagal stimulation

To identify neurons responding to vagal optogenetic stimulation, we computed auROC values comparing neural activity during the vagal stimulation window to baseline. We then performed permutation testing (500 iterations): trial labels (event vs. baseline) were shuffled, auROC was recomputed, and the distribution of shuffled auROC values was generated. Significance thresholds were set at the 10th and 60th percentiles of this null distribution. Neurons with auROC values exceeding the 60th percentile were classified as "stimulation-excited," those below the 10th percentile as "stimulation-suppressed," and all others as non-responsive.

#### Control comparison against masking light

To ensure that observed responses were specific to vagal opto stimulation and not to the visual masking light presented on all trials, we performed a secondary auROC analysis comparing activity during the vagal opto stimulation window to activity during the equivalent time window on Hit trials (which contained only the masking light, no vagal opto stimulation). Only neurons that passed both the primary comparison (vagal opto stimulation vs. baseline) and this control comparison (vagal opto stimulation vs. masking light) were classified as vagal opto-responsive.

#### Reward and licking response detection

For reward delivery and lick onset events, we computed auROC values comparing each event window to the baseline window across all trials. Statistical significance was assessed using permutation testing (500 iterations, 10th/60th percentile thresholds). Neurons responsive to reward or lick onset were also compared against the masking light window (from Hit trials) as a control to ensure specificity. Neurons passing both the event-versus-baseline comparison and the event-versus-control comparison were classified as event-responsive. Neurons classified as responding to either reward delivery or lick onset were combined into a single "reward/licking" category for population-level analyses, as these events are temporally coupled during Hit trials and separating them was not a central goal of this current work.

### Correlation of vagal optogenetic stimulation and reward-related neural activity

We averaged neuronal activity across trials to obtain a single value per neuron for each analysis window, analyzing rewarded and correct-rejection trials separately. For rewarded trials, we computed mean activity from 0 to 2 sec after reward. For correct-rejection trials, we computed mean activity during the 4 sec stimulation window. To assess the relationship between suppressed optogenetic responses and reward-related activity, we computed correlations across neurons separately for each animal between activity during the correct-rejection vagal opto stimulation window and the rewarded-trial measure. We calculated Pearson correlations and used linear fits for visualization.

### Spatial Organization Analysis

To examine the spatial organization of functionally distinct neuronal populations within the imaging field of view, we performed spatial clustering analyses using ROI coordinates from Suite2p segmentation. For each analysis, we extracted the spatial coordinates of each neuron as the median position of its pixel mask (y, x coordinates). We quantified spatial clustering using two complementary approaches: (1) within-group versus across-group distance comparisons, and (2) radial clustering analysis across multiple spatial scales.

#### Within-versus-across distance analysis

For each comparison, we computed pairwise distances between all neuron pairs and classified them as "within-group" (both neurons from the same functional category) or "across-group" (neurons from different categories). To account for unequal group sizes, we used a subsampling approach with 500 iterations per comparison. In each iteration, we randomly sampled neurons from the larger group to match the size of the smaller group, computed the mean within-group distance and mean across-group distance for the balanced subset, and calculated the difference (within minus across). The observed clustering metric was the average of these 500 subsampled differences. Statistical significance was assessed using permutation testing (500 permutations). For each permutation, neuron labels were randomly shuffled while preserving group sizes, and the clustering metric was recalculated using the same subsampling procedure. The two-sided p-value was computed as the fraction of permutations yielding clustering metrics at least as extreme as the observed value (relative to the null distribution mean).

#### Radial clustering analysis

To characterize clustering across spatial scales, we performed radial neighbor analysis at fixed radii (8, 12, 16, 24, 32, 48, 64, 66, 128, 160, and 180 pixels, pixel size = 1.7 µm). For each radius, we counted the number of neighbors within that distance for each neuron, separately tallying same-label neighbors (same functional category) and different-label neighbors (different category). To balance unequal group sizes, we repeated this analysis 500 times with random subsampling (matching the smaller group size), averaging neighbor counts across iterations. We summarized clustering strength by computing the area under the curve (AUC) of the (same-label minus different-label) neighbor count function across radii. Statistical significance of the observed AUC was assessed using permutation testing (500 permutations). For each permutation, neuron labels were randomly shuffled while preserving group sizes, the same-label and different-label neighbor counts were recomputed at each radius using the same subsampling procedure (500 iterations), and the AUC of the permuted (same minus different) curve was calculated. The two-sided p-value was computed as the fraction of permutations yielding AUC values at least as extreme as the observed AUC (relative to the null distribution mean).

#### Choice probability spatial organization

Neurons were divided into two groups based on their choice probability (CP) values: positive-CP neurons and negative-CP neurons, where only neurons passing the 5% false positive rate threshold were included. We applied the within-versus-across distance and radial clustering analyses described above to test whether high-CP and low-CP neurons were spatially segregated.

#### Functional population spatial organization: vagal stimulation vs. reward responses

Neurons were classified into three mutually exclusive categories based on their responses to vagal optogenetic stimulation and reward/licking events: (1) "Light only" -responsive to vagal optogenetic stimulation but not reward/licking; (2) "Reward only" -responsive to reward/licking but not vagal stimulation; (3) "Both" -responsive to both vagal stimulation and reward/licking. Responsiveness was determined by passing both the event-versus-baseline auROC comparison and the event-versus-masking-light control comparison (as described in "Single-Cell Response Classification"). For the spatial analysis, we focused on the comparison between "Light only" and "Reward only" populations, testing whether these two functionally distinct groups exhibited spatial segregation using the within-versus-across distance and radial clustering analyses described above.

#### Neurometric curve spatial organization: Sigmoid vs. Gaussian neurons

Neurons were classified based on the shape of their neurometric curves (as described in "Neurometric curve fitting"): Sigmoid neurons exhibited monotonic frequency-response relationships, while Gaussian neurons showed tuned frequency preferences. We applied the within-versus-across distance and radial clustering analyses described above to test whether Sigmoid and Gaussian neurons were spatially segregated.

#### Cross-animal pooled spatial clustering analyses

To assess spatial clustering across animals, we summed observed clustering metrics (either within-minus-across distance or radial AUC) across all mice and compared this pooled sum to a pooled null distribution obtained by summing the mouse-specific permutation distributions. The pooled two-sided p-value quantified whether the aggregate spatial clustering across the cohort exceeded chance expectations. Individual mouse results were visualized using forest plots showing observed clustering metrics with 65% confidence intervals derived from mouse-specific permutation distributions.

### Dimensionality reduction and clustering

We followed our previously published pipeline^68^. Briefly, data were binned into half-second bins (seven imaging frames per bin) to reduce computational load and smooth noisy fluctuations. Neural activity traces were then filtered to exclude neurons with low mean activity, retaining only cells exhibiting sufficient modulation across the recording session (threshold: absolute mean ≤ 0.5 after normalization). All retained traces were normalized using z-score transformation to standardize response magnitudes across neurons. Dimensionality reduction was performed using a two-stage Laplacian Eigenmap embedding procedure. In the first stage, the high-dimensional neural activity matrix was embedded into a 20-dimensional intermediate space using spectral embedding with nearest-neighbor affinity. This intermediate representation was subsequently projected into a 6-dimensional space (determined as the optimal dimensionality for our data -typically 5-6 intrinsic dimensions) to capture the dominant modes of population variability while preserving local geometric structure. Clustering the data into eight clusters was subsequently performed as previously described^68^. This allowed us to assign each time point to a specific cluster and describe population dynamics as cluster transition dynamics. Final cluster labels were assigned to all time points, based on their nearest cluster center. Population activity sequences were aligned to trial onset times and organized by behavioral outcomes (hits, correct rejections, false alarms) to examine state dynamics across experimental conditions.

#### Trial-wise neural state similarity analysis

To quantify the relationship between population activity dynamics and behavioral outcomes, we performed a trial-wise similarity analysis. Clusters were first selected for analysis based on three criteria that ensured we captured reliable task-evoked changes: first, they must appear in at least one behavioral condition in 30% or more of trials; second, they must exhibit sustained activation (at least two consecutive bins above the 30% threshold) in at least one condition; and third, they must not show significant baseline presence, defined as appearing in 20% or more of trials in any of the first three time bins after trial onset. This baseline exclusion criterion ensured that analyses focused on task-evoked rather than pre-stimulus activity patterns. For each trial, a binary feature vector was constructed representing the presence or absence of each selected cluster at each time bin, excluding the baseline period. This yielded a trial-by-feature matrix where each row represented a trial’s cluster activation pattern across time. Pairwise cosine similarity was then computed between all pairs of trials, generating a trial-by-trial similarity matrix. Within-condition similarity was quantified as the average cosine similarity between all pairs of trials within the same behavioral condition (excluding self-comparisons). Between-condition similarity was computed as the average cosine similarity between trials from different behavioral conditions. This trial-wise approach allowed us to assess whether population activity dynamics were more similar within behavioral conditions than across conditions, and whether specific condition pairs showed higher or lower neural similarity. Values below 10% in proportion vectors were set to zero to reduce noise from sporadic cluster appearances.

#### Cross-animal population activity analysis

To assess the prevalence of task-evoked population activity dynamics across animals, we computed the fraction of animals exhibiting active cluster states at each time point for each behavioral condition. For each animal and time bin, we determined whether any cluster exceeded an activity threshold (present in at least 30% of trials for that condition). The binary indicator of cluster presence was then averaged across animals at each time bin, yielding a time series representing the fraction of animals showing any active cluster. This analysis quantified whether task-evoked population dynamics were consistently present across the population of recorded animals or were animal-specific phenomena.

#### Statistical aggregation and visualization

Results were aggregated across animals by first computing cluster-level similarity metrics for each mouse independently, then averaging across clusters within each animal to obtain per-mouse summary statistics. Group-level statistics were computed as the mean and standard error of the mean across animals, with individual animal data points overlaid on bar plots to visualize inter-animal variability. For visualization of cluster dynamics, we plotted time courses showing the mean proportion of trials in which each selected cluster was active as a function of time, separately for each behavioral condition.

Nodose ganglion ROI Signal Extraction and Calcium Trace Processing:

#### Raw data processing and ROI definition

Two-photon movies were reshaped into time × height × width arrays and split into green (GCaMP) and red (motion control) channels according to the scanning cycle.

For each imaging plane, we computed standard-deviation projections of the green and red channel stacks. Elliptical regions of interest (ROIs) were manually placed on these projections to define individual neurons.

#### Fluorescence trace extraction

Mean fluorescence traces were extracted by averaging pixel intensities within each ROI mask across all frames. This procedure was applied separately to green and red channels for all defined ROIs.

#### Trace preprocessing and motion correction

Extracted fluorescence traces were low-pass filtered (Butterworth filter, cutoff frequency 0.02 Hz) to remove high-frequency noise, then bleaching-corrected using constrained double-exponential fits to account for potential photobleaching over the recording session. Traces were further denoised using a 10-second centered running mean. Motion artifacts were corrected by regressing the red control channel signal onto the green channel signal; the fitted motion component was subtracted from the green channel recording. For visualization and statistical testing, traces were z-scored using a baseline window extending 240 seconds before the first stimulus in each recording segment.

#### Responder identification

For each ROI, we evaluated the correlation between the motion-corrected green signal and the red control channel. High negative correlations (R< -0.5, indicating over-correction of motion) prompted re-evaluation using single-channel green preprocessing without red channel subtraction. High positive correlations (R> 0.5, indicating uncorrected shared motion artifacts) required testing both channels separately, with responses validated only if the green channel showed stimulus-locked activity distinct from the red channel. Low correlations between channels indicated successful motion correction, and standard motion-corrected green channel analysis was applied. Within each stimulus presentation (intestinal balloon inflation to deflation, plus a fixed 50 sec post-stimulus window), we applied a dynamic threshold relative to each cell’s pre-stimulus baseline to detect responses. Responses were considered significant if post-stimulus activity exceeded 5 standard deviations above baseline for a minimum duration of 5 seconds, after verifying a flat baseline slope (≤0.02 Δz/s). When multiple ROIs were defined for the same neuron across planes or channels, indices were applied to retain only non-redundant ROIs for analysis.

### Vagal optogenetic response decay analysis

For the decay analysis, we used deconvolved activity obtained using the Suite2p pipeline^124^. We z-scored signals for each neuron, segmented them into trial-aligned traces with a 4-sec pre-trial period, and analyzed only on vagal opto stimulation trials to avoid reward-related activity. To focus on stimulation-selective responses, we restricted analyses to neurons identified in the preceding auROC-based analysis as significantly increasing activity during vagal opto stimulation but not significantly responsive to reward. For each neuron and trial, we baseline-subtracted activity using the 1-sec period immediately preceding stimulation onset and smoothed it with a 5-frame (∼330 msec) moving average. We retained trials for decay fitting only if they showed a detectable optogenetic response during the stimulation window, defined as activity exceeding baseline by at least 2 standard deviations for at least two consecutive frames. Within responsive trials, we identified the response peak as the first significant local maximum during the stimulation window, with criteria on minimum height, prominence, and inter-peak distance. If we could not detect such a peak, we used the maximal value within the stimulation window. We quantified the time required for the response to decay to one-third of peak amplitude, the mean response at the end of the stimulation period (odor onset) relative to peak. We included trials only if they passed the response-detection criterion, had a peak amplitude of at least 0.25, and showed acceptable fit quality (R² ≥ 0.2). We computed neuron-level summary measures across valid trials, and retained only neurons with at least five valid trials for subsequent analyses.

---

**Extended Data Fig. 1:**
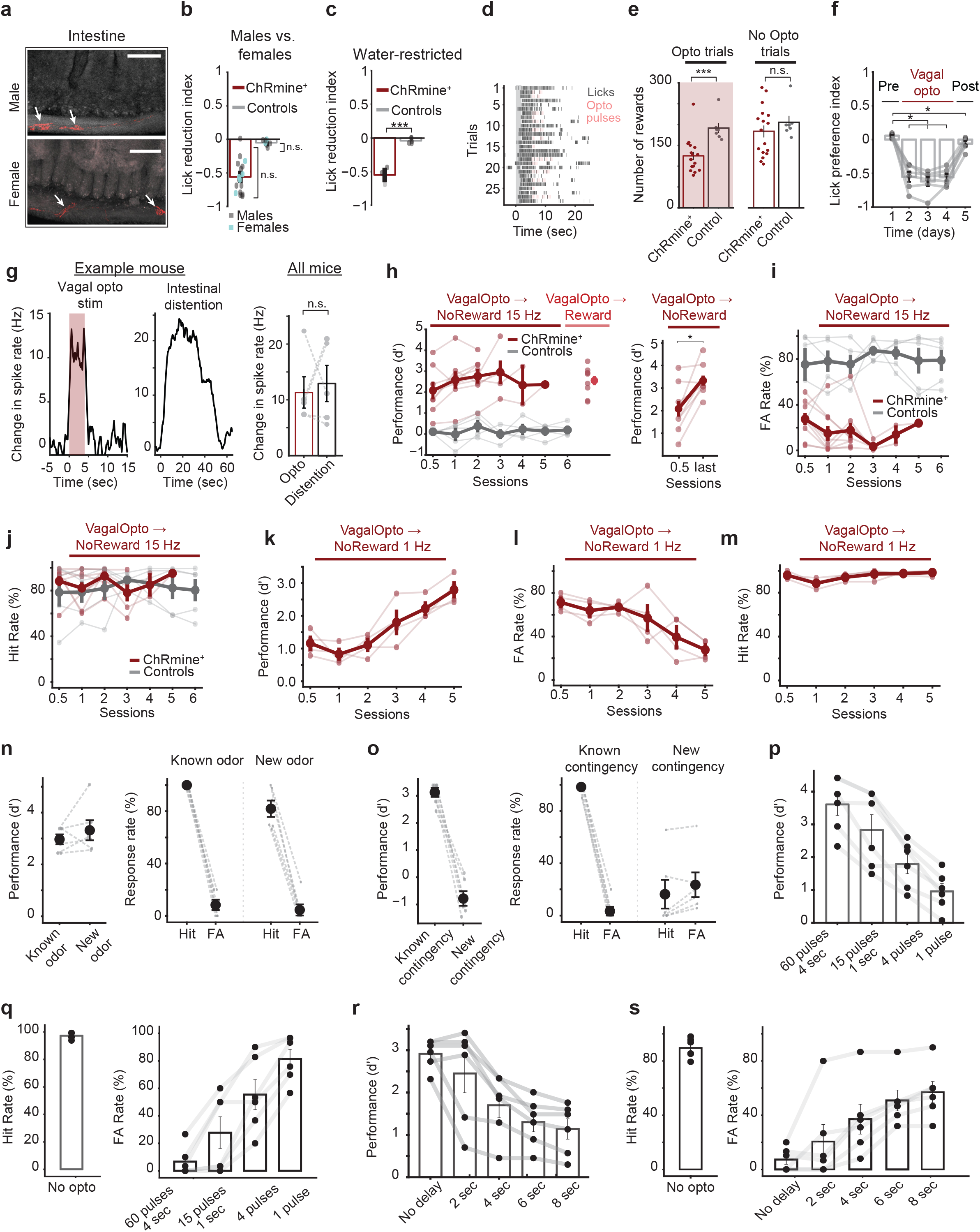
Control experiments for non-invasive optogenetic activation of NG-OxtR^+^ for interoceptive detection tasks. (a) NG-OxtR^+^ terminals expressing ChRmine-mScarlet innervating the intestine and forming intra ganglionic laminar endings (IGLEs, labeled with white arrows) in both male and female mice. Scale bar = 100 µm. (b) Quantification of lick reduction index during the sucrose consumption task, showing similar responses between males (n = 12 ChRmine+, 5 controls) and females (n = 5 ChRmine+, 2 controls). Bars show mean ± SEM across mice. Males vs. females within the ChRmine+ group: p = 0.8 (Whitney U), males vs. females within the control group: p = 0.38 (Whitney U). (c) Quantification of lick reduction index during the sucrose consumption task in water restricted ChRmine+ (n = 3) and control (n = 5) mice, p = 0.00013 (independent t-test). Effect size, Hedges g = 5.47. Bars show mean ± SEM across mice. (d-f) Activation of NG-OxtR^+^ does not cause substantial preference/aversion. (d) Example lick raster from the mouse shown in Fig. 1i, illustrating licking behavior during the closed-loop sucrose consumption task under food restriction. Trials are presented in chronological order, rather than grouped by trial type as in Fig 1i. Licking cessation is restricted to the optogenetic stimulation period, with licking resuming in subsequent trials following stimulation. (e) Quantification of the number of rewards (triggered by licking) consumed during the closed-loop sucrose consumption task. Bars represent mean ± SEM across mice (ChRmine^+^, n = 17; controls, n = 7). *Left*: Total number of rewards consumed during opto trials. ChRmine^+^ mice consumed significantly fewer rewards than control mice p = 0.0007 (Welch’s t-test, Hedgesg = 1.74). *Right*: Total number of rewards consumed during no-opto trials. Reward consumption did not significantly differ between ChRmine^+^ and control mice, p = 0.3 (Welch’s t-test). (f) Quantification of lick preference index for each day of the conditioned taste preference experiment (n = 6 mice). Bars show mean ± SEM across mice. As expected from the immediate effect of stimulation to reduce licking, there was a substantial reduction in preference for the juice paired with non-invasive NG-OxtR^+^ optogenetic activation during conditioning days from 0.057 to -0.55 (adj.P = 0.03, Wilcoxon with FDR correction, r = -0.855). When comparing day 1 and day 5 (pre stimulation vs. post stimulation, in which there was no vagal optogenetic stimulation), there was a very small yet significant decrease from 0.057 to -0.06, likely reflecting our initial biased choice of pairing stimulation with the mildly preferred juice (adj. P = 0.03, Wilcoxon with FDR correction, Effect size (Rank-Biserial Correlation): r = -0.86). (g) Comparison of vagus nerve activity evoked by intestinal distention and non-invasive NG-OxtR^+^ optogenetic activation. An approximately 1.5-fold intestinal distention was used to approximate the increase in intestinal diameter observed following feeding. *Left* and *middle*: change in vagal spike rate evoked by 15 Hz optogenetic stimulation (*left*) and approximately 1.5-fold intestinal distention induced by infusion of saline (*middle*) in an example mouse. Shading indicates the optogenetic stimulation period. *Right*: comparison of the change in vagal spike rate evoked by optogenetic stimulation and intestinal distention across mice P = 0.84 (paired t-test, n = 4 mice). Gray points and connecting lines indicate individual mice; bars show mean ± SEM. (h) *Left*: learning of the VagalOpto→NoReward task in ChRmine^+^ mice (n=9), but not in control mice (n=5). Values are d’ (discriminability index) over days, showing similar performance between VagalOpto→NoReward (dark red) and VagalOpto→Reward (light red) tasks in ChRmine mice, using 15 Hz stimulation. In contrast, control mice failed to learn the VagalOpto→NoReward task over multiple training days (gray). Thicker lines indicate mean ± SEM across mice. *Right*: d’ calculated for each mouse at the first half of the first session, compared to the last session, shows improved discriminability in the VagalOpto→NoReward task with training P = 0.012 (Wilcoxon test), Effect size (Rank-Biserial Correlation): r = -0.79. (i) False alarm rates across training sessions in ChRmine^+^ mice and control mice during the VagalOpto→NoReward task using 15 Hz stimulation. Thick lines show mean ± SEM across mice; thin lines show individual mice. (j) Hit rates across training sessions in ChRmine^+^ mice and control mice during the VagalOpto→NoReward task, using 15 Hz stimulation. Thick lines show mean ± SEM across mice; thin lines show individual mice. (k) d’ values across training sessions in ChRmine^+^ mice during the VagalOpto→NoReward task using 1 Hz stimulation. Thick line shows mean ± SEM across mice; thin lines show individual mice. (l) False alarm rates across training sessions in ChRmine^+^ mice during the VagalOpto→NoReward task using 1 Hz stimulation. Thick line shows mean ± SEM across mice; thin lines show individual mice. (m) Hit rates across training sessions in ChRmine^+^ mice during the VagalOpto→NoReward task using 1 Hz stimulation. Thick line shows mean ± SEM across mice; thin lines show individual mice. (n) Generalization to a new odor cue in the VagalOpto→NoReward task. *Left*: d’ values during task performance with the known odor and with a new odor. *Right*: hit and false alarm response rates for the known odor and new odor conditions. Dots indicate individual mice; bars show mean ± SEM across mice. (o) Performance after changing the behavioral contingency. *Left*: d’ values during the known contingency and after changing to a new contingency. *Right*: hit and false alarm response rates under the known contingency and the new contingency. Gray dots and their connecting lines indicate individual mice; Black dots show mean ± SEM across mice. (p) d’ values during the VagalOpto→NoReward task with different optogenetic stimulation pulse numbers with the same pulse frequency (15 Hz): 60 pulses (4 s); 15 pulses (1 s); 4 pulses (∼270 ms); and 1 pulse. Gray dots and their connecting lines indicate individual mice; Black dots show mean ± SEM across mice. (q) Hit and false alarm rates in the different pulse-number experiment in ‘p’. *Left*: hit rate on no-opto rewarded trials. *Right*: false alarm rates during VagalOpto→NoReward trials with 60 pulses, 4 s; 15 pulses, 1 s; 4 pulses; and 1 pulse. Dots indicate individual mice; bars show mean ± SEM across mice. (r) d’ values during the VagalOpto→NoReward task with different delays between optogenetic stimulation offset and odor onset (4 s at 15 Hz): no delay, 2 s, 4 s, 6 s, and 8 s delay. Dots and connecting lines indicate individual mice; bars show mean ± SEM across mice. (s) Hit and false alarm rates in the delay experiment in ‘r’. *Left*: hit rate on no-opto rewarded trials. *Right*: false alarm rates during VagalOpto→NoReward trials with no delay, 2 s, 4 s, 6 s, and 8 s between optogenetic stimulation offset and odor onset (4 s at 15 Hz). Dots indicate individual mice; bars show mean ± SEM across mice.

**Extended Data Fig. 2:**
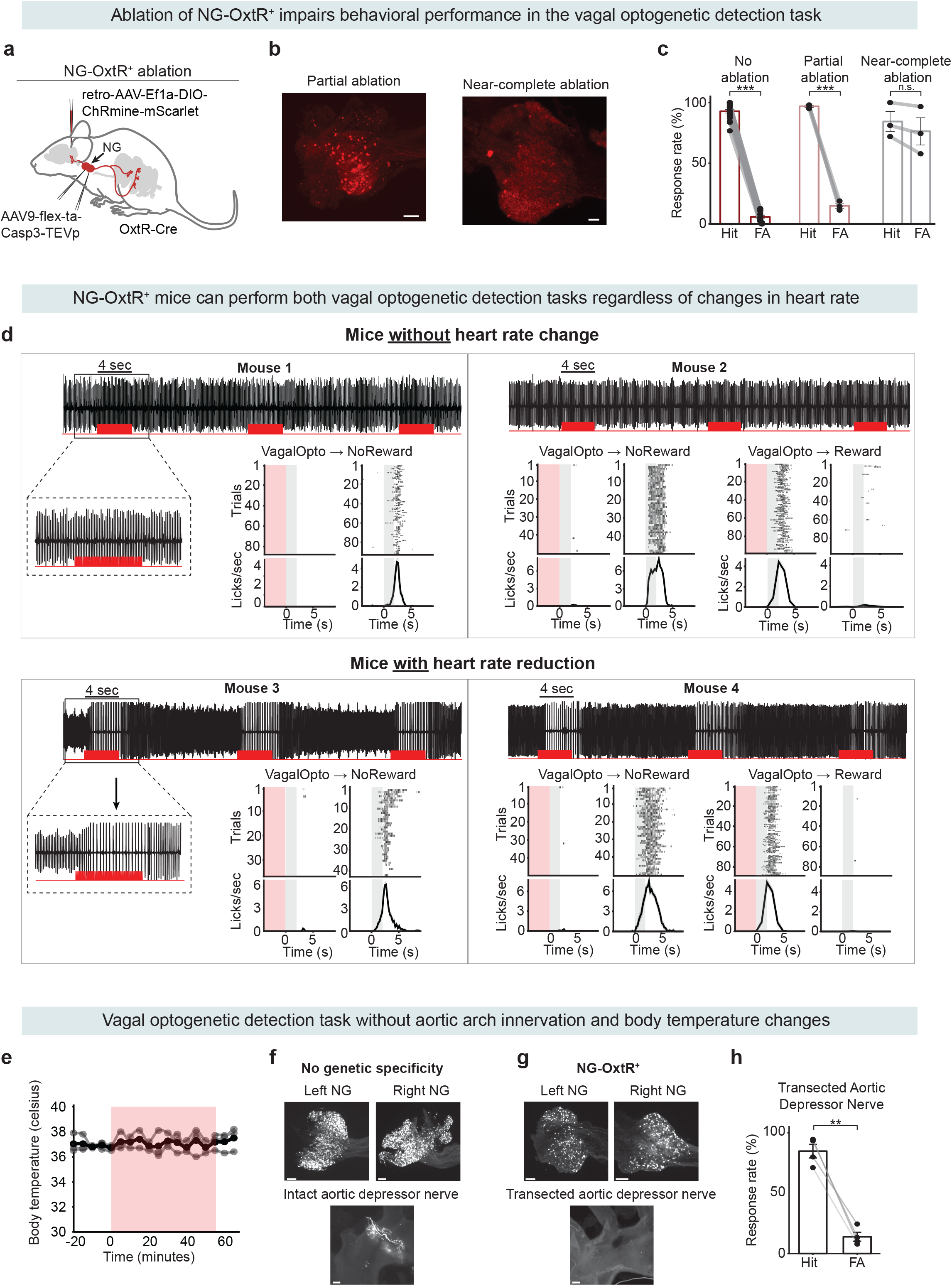
Performance of the vagal optogenetic detection task requires NG-OxtR^+^ neurons, is independent of aortic arch projections, and does not affect body temperature. (a-c) NG-OxtR^+^ specific ablation using Caspase 3 hinders behavioral performance in the VagalOpto→NoReward task. (a) Schematic of the experimental approach. Retro-AAV-Ef1a-DIO-ChRmine-mScarlet was injected bi-laterally into the NTS of OxtR-Cre mice. Additionally, AAV9-flex-taCasp3-TEVp was injected bi-laterally into the NG to specifically ablate NG-OxtR^+^ neurons. (b) Example histology from two different mice demonstrating partial ablation of NG-OxtR^+^ neurons (*left*), and successful, near-complete, ablation (*right*). Scale bar = 200 µm. (c) Summary of Hits and FAs in the VagalOpto→NoReward task of mice without ablation (same plot as Fig. 1n, shown for comparison), partial ablation, and near-complete ablation of NG-OxtR^+^ neurons. p = 3.6×10^-21^ (paired t-test, n=19), Cohen’s d = -12.1. *Middle*: mice that underwent bi-lateral NG injections of Caspase 3 but showed partial ablation, p = 8.4×10^-4^ (paired t-test, n=3), Cohen’s d = -19.9. *Right*: mice with successful, near-complete ablation of NG-OxtR^+^ neurons, p = 0.129 (paired t-test, n=3). Bars show mean ± SEM across mice. (d) Performance in the VagalOpto→NoReward and VagalOpto→Reward tasks is independent of potential changes in heart rate. Heart rate measurements in anesthetized mice during NG-OxtR^+^ optogenetic stimulation (same stimulation parameters as in the behavioral task). *Top*: example of two mice without a change in heart rate in response to the optogenetic stimulation. *Bottom*: example of two mice with a reduction in heart rate in response to the optogenetic stimulation. In the electrocardiogram traces, each peak indicates one heartbeat, and each red horizontal line indicates 4 sec of vagal optogenetic stimulation. All mice had similarly high performance in the VagalOpto→NoReward and VagalOpto→Reward tasks, regardless of whether there was a change in heart rate. Red rectangles indicate optogenetic stimulation time, and gray rectangles indicate odor cue presentation. Gray ticks indicate licks. (e) Performance in the VagalOpto→NoReward task does not induce changes in body temperature. Temperature was measured during the VagalOpto→NoReward task every 5 min (n = 3 mice). Each mouse’s measurements are indicated in gray, the black line is the mean across mice, and the red rectangle indicates the time of the VagalOpto→NoReward session. (f) Representative histological analysis of the aortic arch and NG from a WT mouse injected with Cre-independent retroAAV9 -CBA-mScarlet3 to the NTS, lacking genetic targeting of a specific neuronal population. *Top*: Left and right NG showing retrogradely labeled neurons. *Bottom*: Representative image of the aortic arch from the same mouse showing labeled aortic depressor nerve innervation, demonstrating the ability to anatomically identify aortic depressor nerve projections within the aortic arch. (g) Representative histological analysis of the aortic arch and NG from an OxtR-Cre mouse injected with Cre-dependent retroAAV-EF1a-DIO-ChRmine-mScarlet into the NTS to label NG OxtR^+^ neurons, and then subjected to bilateral aortic depressor nerve transection. *Top*: Left and right NG showing labeled OxtR neurons. *Bottom*: Representative image of the aortic arch showing the absence of detectable labeled aortic arch innervation following bilateral aortic depressor nerve transection. (h) Summary of Hits and FAs in the VagalOpto→NoReward task of mice that underwent aortic depressor nerve transection (paired t-test, n = 4, p = 0.002, Cohen’s d = -5.18)

**Extended Data Fig. 3:**
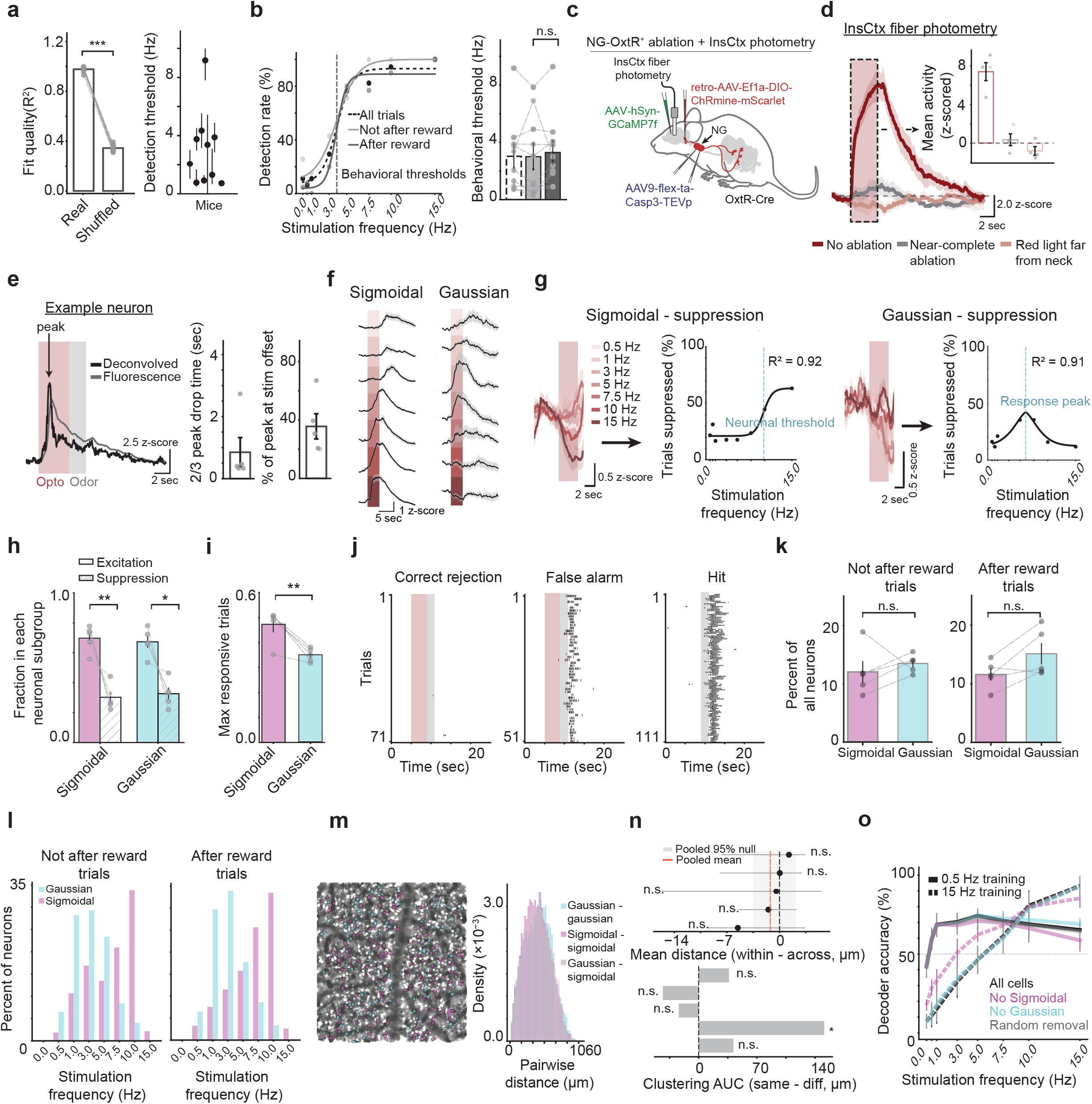
Further analyses of InsCtx optogenetic frequency responses during the psychophysics task. (a) *Left*: Fit quality of behavioral psychometric curves relating detection rate to stimulation frequency. Bars show R² values for psychometric fits to the real behavioral data and to shuffled behavioral data. Bars show mean ± SEM across mice; dots indicate individual mice. ***p < 0.001, Cohen’s d:199. *Right*: psychometric detection threshold for each mouse, estimated from the fitted sigmoid curve. Error bars indicate 95% bootstrap confidence intervals obtained from 1,000 bootstrap resamples, with trials resampled with replacement separately within each stimulation frequency. (b) *Left*: behavioral performance as a function of stimulation frequency, shown as detection rate across all trials (black dashed line), not-after-reward trials (light gray), and after-reward trials (gray). *Right*: behavioral detection thresholds for each condition across mice. Bars show mean ± SEM across mice; dots indicate individual mice. p = 0.27 (Not after reward vs. after reward, Wilcoxon test) (c) Schematic of the experimental approach. Retro-AAV-EF1α-DIO-ChRmine-mScarlet was injected bilaterally into the NTS of OxtR-Cre mice. AAV9-flex-taCasp3-TEVp was also injected bilaterally into the NG to selectively ablate NG-OxtR⁺ neurons. In addition, GCaMP7f was injected into the InsCtx, and an optic fiber was implanted above the injection site for fiber photometry recordings. (d) *Left*: mean z-scored InsCtx fiber photometry neuronal activity in response to non-invasive vagal optogenetic stimulation without NG-OxtR^+^ ablation, with near-complete NG-OxtR^+^ ablation, and with red light delivered far from the neck. Pink shading marks the vagal opto stimulation period. *Right*: quantification of mean activity in the indicated stimulation window. Bars show mean ± SEM across mice; dots indicate individual mice. (e) Analysis of InsCtx neuronal decay kinetics in responses to NG-OxtR_+_ optogenetic activation. *Left*: example from an individual InsCtx neuron of fluorescence and deconvolved activity traces during non-invasive vagal optogenetic stimulation. Pink shading marks the stimulation period; gray shading marks the odor cue period. Arrow indicates the response peak. *Right*: quantification of the time for the response to decay to two-thirds of the peak amplitude and the percentage of peak activity remaining at odor onset. Bars show mean ± SEM across mice; dots indicate individual mice (median across all neurons per mouse). (f) Full traces of the example neurons with Sigmoidal (*left*) or Gaussian (*right*) frequency– response functions presented in Fig. 2e-f. Trial-by-trial z-scored activity sorted by stimulation frequency. Pink bar indicates the optogenetic vagal stimulation period. (g) Example neurons with suppressed Sigmoidal (*left*) or Gaussian (*right*) frequency– response functions. Similar to Fig. 2e,f but for suppressed responses. Red shaded plots to the left of each black fitted curve show overlaid mean z-scored activity traces for all stimulation frequencies (color-coded as labeled in the figure); pink bars mark the vagal optogenetic stimulation period. Black fitted curves show the fraction of trials neuronal activity was significantly suppressed versus stimulation frequency; R² quantifies goodness of fit. The dashed vertical line indicates the neuronal threshold for the Sigmoidal neuron or the response peak of the Gaussian neuron. (h) Percent of neurons significantly excited versus significantly suppressed in each response type group (Sigmoidal, Gaussian). Bars show mean ± SEM across mice (n = 5); dots indicate individual mice. Paired t-tests per response type group: Sigmoid p=0.007, Cohen’s d: -2.3 ; Gaussian p=0.013, Cohen’s d: -1.9. Two-way RM-ANOVA with factors type (Sigmoid, Gaussian) and condition (excited, suppressed), condition: p=0.004; type: p=1.0, interaction: p=0.65. (i) Maximum fraction of trials neurons were responsive across frequencies for Sigmoidal vs. Gaussian neurons (computed per neuron, then averaged per mouse). Bars show mean ± SEM across mice (n = 5); dots indicate individual mice. Sigmoidal vs. Gaussian: p = 0.007 (paired t-test), Cohen’s d: -2.2. (j) Example lick raster showing licking behavior of a mouse during the psychophysics task. The gray rectangles mark odor cue presentation, pink rectangles mark the optogenetic vagal stimulation during all stimulation frequencies and gray ticks mark individual licks. (k) Fraction of neurons classified as Sigmoidal vs. Gaussian out of all recorded neurons per mouse for not-after-reward trials (*left*) and after-reward trials (*right*). Bars show mean ± SEM across mice (n = 5); dots indicate individual mice. n.s. indicates not significant, not-after-reward: p = 0.62, after-reward: p = 0.125 (Wilcoxon test). (l) Distribution of neuronal thresholds (Sigmoidal) and response peaks (Gaussian), pooled across all recorded neurons from all mice for not-after-reward trials (*left*, Sigmoidal: n=319 neurons, 63.8 ± 13.4 neurons per mouse; Gaussian: n=368 neurons, 73.6 ± 15.6 neurons per mouse; from 5 mice) and after-reward trials (*right*, Sigmoidal: n=304 neurons, 60.8 ± 11.7 neurons per mouse; Gaussian: n=383 neurons, 76.6 ± 11.4 neurons per mouse; from 5 mice). (m) Spatial organization example. *Left*: two-photon field of view with neurons colored by response type (magenta: Sigmoidal; cyan: Gaussian). *Right*: example histogram from the same mouse showing distances between pairs of neurons for within-Gaussian, within-sigmoidal, and across-types. Note no clear spatial organization. (n) Summary of spatial clustering analysis across mice. *Top*: mean distance difference (within type minus across type) per mouse; the shaded band indicates the pooled 95% null confidence interval, and the red line shows the pooled mean. *Bottom*: clustering separability quantified by AUC for same vs. different type pairs. Calculation of p values for the top Forest plot: two-sided permutation on the within minus across distance, with label shuffling and subsampling to the smaller group size. Calculation of p values for the bottom clustering plot: same procedure as the top plot, but on the AUC of same minus different neighbor counts across radii. *p<0.05,, ns p≥0.05. (o) Decoder accuracy as a function of stimulation frequency after training the decoder on 0.5 Hz (black) or 15 Hz (gray). Additional curves show decoder accuracy after removal of Sigmoidal neurons, Gaussian neurons, or a randomly selected matched number of neurons. Curves show mean ± SEM across mice (n = 5). The gray dashed horizontal line marks chance decoding (50%).

**Extended Data Fig 4:**
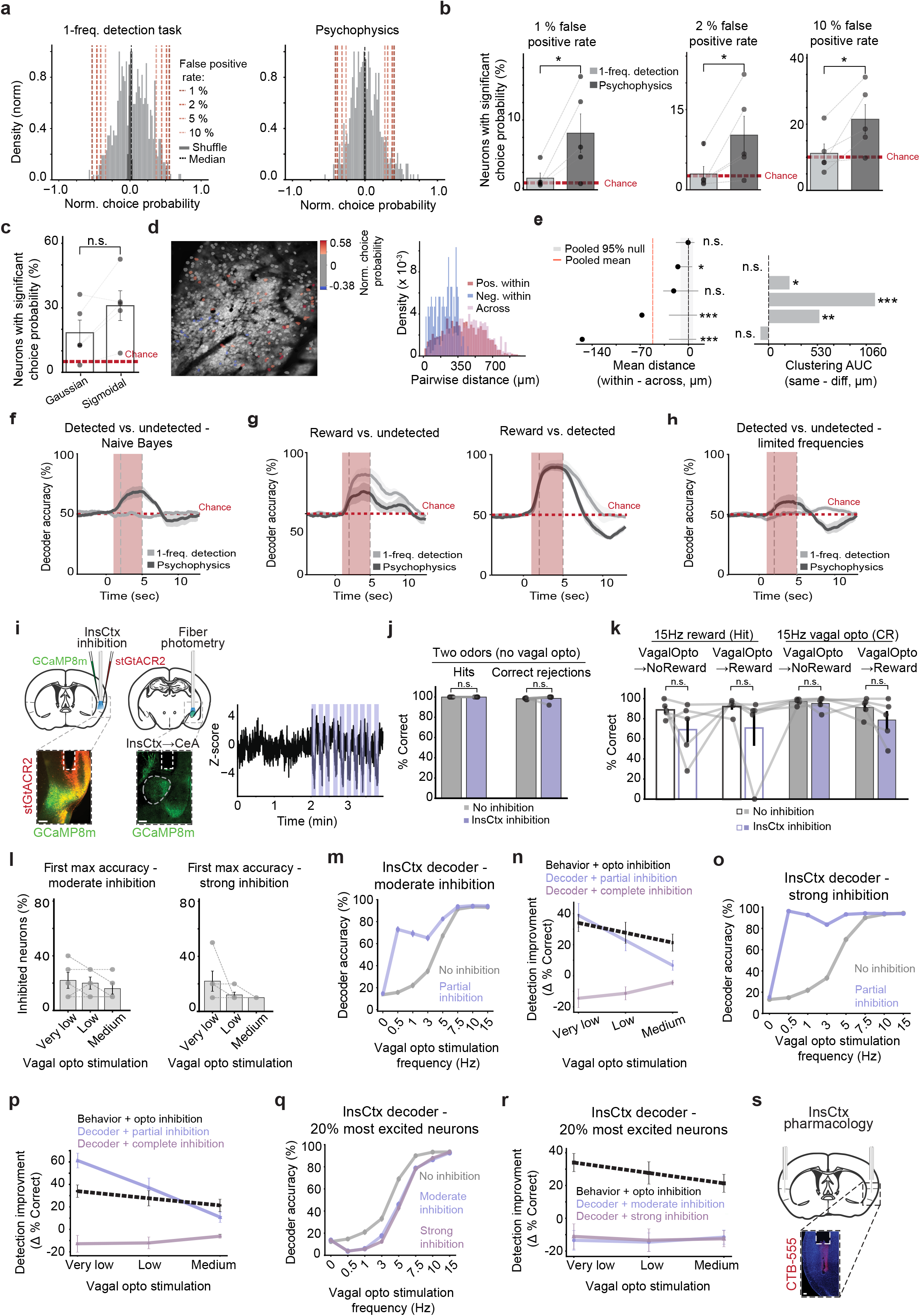
Further analyses of InsCtx choice probability; spatial organization; decoding with/without inhibition; control experiments for optogenetic inhibition of InsCtx. (a) Example mouse showing the distributions of single-neuron normalized choice probability (light gray, between -1 and +1) for the 1-frequency detection task (*left*) and the psychophysics task (*right*). Histograms show the density of normalized choice-probability values across neurons. Red dashed lines indicate 1%, 2%, 5%, and 10% false-positive thresholds. Dark gray bars show shuffle distributions. (b) Fraction of neurons with significant choice probability (positive or negative) in each task at different false-positive thresholds. Fractions are out of all recorded neurons per mouse. Bars show mean ± SEM across mice (n = 5); dots indicate individual mice. Red dashed line marks the nominal false-positive rate. Task comparison -1%: p = 0.048, Cohen’s d:1.3; 2%: p = 0.047, Cohen’s d:1.3; 10%: p = 0.031, Cohen’s d:1.5 (paired t-tests). (c) Fraction of neurons with significant choice probability within each response-type group, Gaussian and Sigmoidal. Fractions are out of all neurons of the corresponding response type per mouse. Bars show mean ± SEM across mice (n = 5); dots indicate individual mice. Red dashed line marks the nominal false-positive rate. Gaussian vs. Sigmoidal: n.s. (paired test). (d) Spatial organization example (5% choice probability threshold). *Left*: two-photon field of view with neurons colored by normalized choice probability (gray in the color bar: non-significant values). *Right*: example histogram from the same mouse showing pairwise distances for high choice probability pairs, low choice probability pairs, and across condition pairs. (e) Summary of spatial clustering across mice. *Top*: mean difference in pairwise distance (within minus across) per mouse; the shaded band indicates the pooled 95% null and the red line shows the pooled mean. *Bottom*: clustering separability quantified by AUC for same vs. different condition pairs per mouse. Forest plot p values: two-sided permutation on the within minus across distance, with label shuffling and subsampling to the smaller group size. Clustering p values: same procedure on the AUC of same minus different neighbor counts across radii. * p<0.05, ** p<0.01, *** p<0.001, ns p≥0.05. (f) Time course of Naive Bayes decoder accuracy for detected vs. undetected trials. Mean ± SEM across mice (n = 5). Red dashed line: chance (50%). Pink bar: optogenetic stimulation window. Gray dashed vertical line: decoder training time period. (g) Time course of SVM decoder accuracy for reward vs. undetected (*left*) and reward vs. detected (*right*). Mean ± SEM across mice (n = 5). Red dashed line: chance (50%). Pink bar: stimulation window. Gray dashed vertical line: decoder training time period. (h) Time course of SVM decoder accuracy for detected vs. undetected using a limited frequency set (see Methods). Mean ± SEM across mice (n = 5). Red dashed line: chance (50%). Pink bar: stimulation window. Gray dashed vertical line: decoder training time period. (i) Validation of InsCtx inhibition using stGtACR2. *Left*: schematic of the experimental setup. We co-injected AAV1-CaMKIIa-stGtACR2-FusionRed and AAV5-mCaMKIIa-jGCaMP8m into the InsCtx. We implanted an optic fiber above the InsCtx for optogenetic inhibition, and another optic fiber above the central amygdala to record fiber photometry signals from InsCtx axons innervating the central amygdala. *Right*: Representative fiber photometry trace. Purple rectangles mark the time windows of InsCtx inhibition. (j) Summary of performance in the 2-odor Go/NoGo task (no vagal optogenetics). In this task, odor A indicated that licking will be rewarded with sugar-water (∼5µL, 600mM sucrose, ‘hits’, *left*), and odor B indicated no reward will be delivered upon licking (‘correct rejections’, *right*). We used optogenetic inhibition of the InsCtx in 50% of trials for 5 seconds before odor cue onset, as in all other behavioral tasks we performed with InsCtx inhibition. Bars show mean (±SEM) percent of hits and correct rejections across mice (n=6). Both conditions had 100% for hits (no statistical test performed); correct rejections did not differ significantly, p = 0.44 (Wilcoxon test). (k) Percent correct hits (*left*) and correct rejections (*right*) during the 1-frequency VagalOpto→NoReward and VagalOpto→Reward tasks at 15Hz stimulation. VagalOpto→NoReward hits: p = 0.31; VagalOpto→Reward hits: p = 0.063; VagalOpto→NoReward correct rejections: p = 0.625; VagalOpto→Reward correct rejection: p = 0.068 (Wilcoxon test). (l) Proportion of suppressed neurons required to reach the first maximum in decoding accuracy with moderate inhibition (*left*) and strong inhibition (*right*), binned by vagal-opto stimulation level (very low, low, medium). Bars show mean ± SEM across mice (n = 5); dots indicate individual mice. See Methods for further information. (m) InsCtx population decoder accuracy with artificially adding moderate optogenetic inhibition (example from one mouse). Lines with markers show decoder accuracy as a function of vagal opto stimulation frequency for no added inhibition (gray) and partial inhibition (20% of the population, light purple). Values are mean ± SEM across 500 decoder runs. (n) Improvement in detection (Δ% correct) with increasing inhibition, binned by vagal opto stimulation level (per mouse, levels selected from that mouse’s psychometric curve; see Methods). Black dashed line: behavior with optogenetic InsCtx inhibition. Light purple: decoder with partial inhibition. Purple: decoder with complete inhibition. Values are mean ± SEM across mice (n = 5). (o) InsCtx population decoder accuracy with strong inhibition (example from one mouse). Lines with markers show decoder accuracy as a function of vagal opto stimulation frequency for no inhibition (gray), partial inhibition (20% of the population, light purple), and complete inhibition (100% of the population, purple). Values are mean ± SEM across 500 decoder runs. (p) Change in detection (Δ% correct) with increasing inhibition, binned by vagal opto stimulation level (per mouse, levels selected from that mouse’s psychometric curve; see Methods). Black dashed line: behavior with InsCtx optogenetic inhibition. Light purple: decoder with partial inhibition. Purple: decoder with complete inhibition. Values are mean ± SEM across mice (n = 5). (q) InsCtx population decoder accuracy with artificial inhibition of the 20% most excited neurons (example from one mouse). Lines with markers show decoder accuracy as a function of vagal optogenetic stimulation frequency under three conditions: no inhibition (gray), moderate inhibition (light purple), and strong inhibition (purple). Values represent mean ± SEM across 500 decoder runs. (r) Change in detection (Δ% correct) with increasing inhibition of the 20% most excited neurons, binned by vagal optogenetic stimulation level (per mouse; levels selected from that mouse’s psychometric curve; see Methods). Black dashed line: behavioral performance with InsCtx inhibition. Light purple: decoder with moderate inhibition. Purple: decoder with strong inhibition. Values represent mean ± SEM across mice (n = 5). (s) Schematic and representative histology of InsCtx pharmacological inhibition. We injected fluorescently labeled CTB-555 (red) through the cannula and injector to verify the injection.

**Extended Data Fig. 5:**
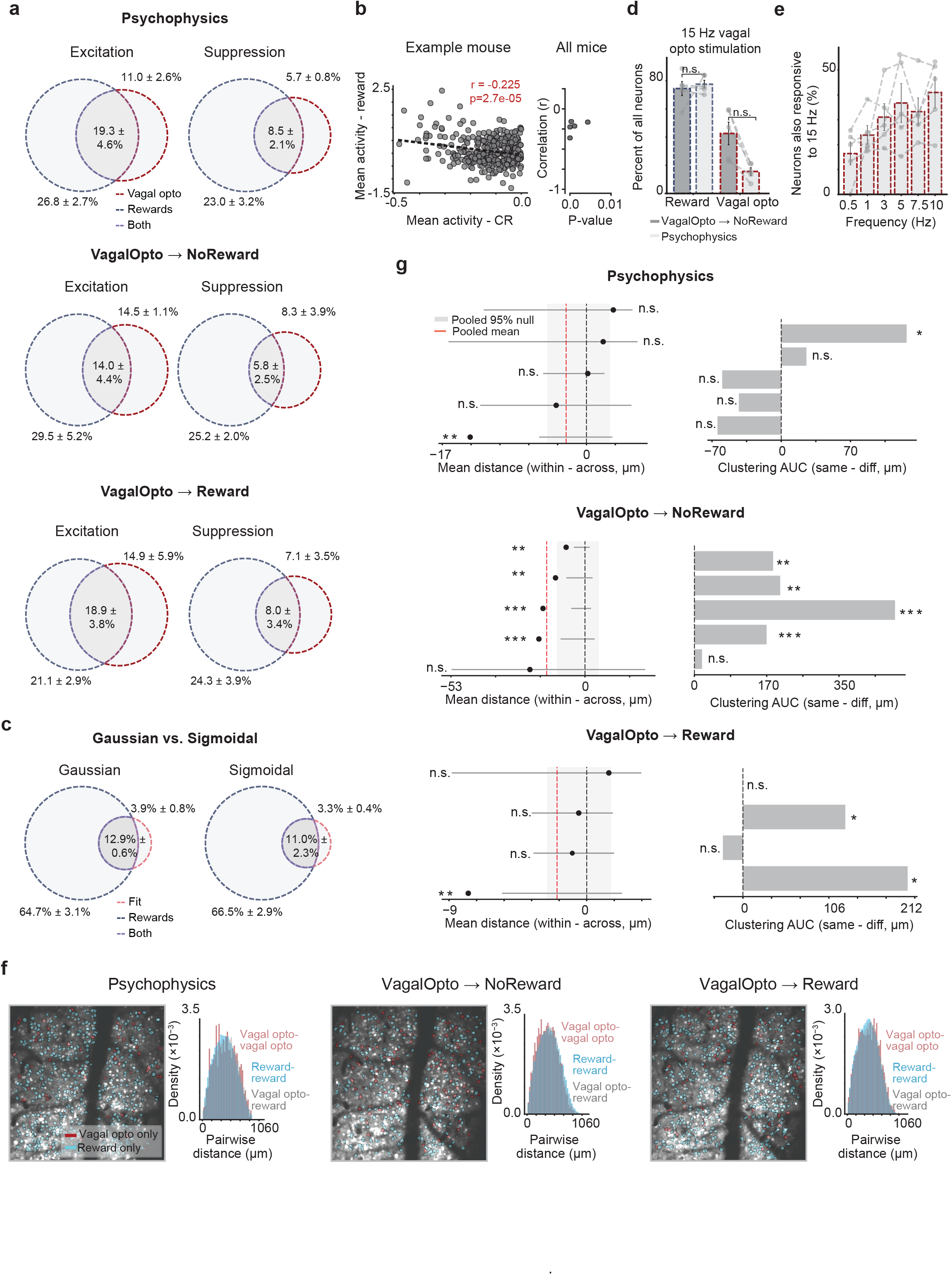
Further analyses of InsCtx excited vs. suppressed responses across different tasks; spatial clustering of responsive neurons. (a) Venn diagrams showing the fraction of neurons significantly responsive to reward (blue dashed circle), vagal opto stimulation (red dashed circle), or both (overlap, purple) for excited and suppressed responses in each behavioral task: psychophysics, VagalOpto→NoReward, and VagalOpto→Reward. Values are mean ± SEM across mice (n = 5). (b) Relationship between vagal optogenetic stimulation and reward responses in co-responsive neurons that were suppressed by vagal optogenetic stimulation on detected trials. *Left*: mean vagal opto activity versus mean reward activity for individual neurons in an example mouse. Dashed line indicates the linear fit, with Pearson’s r and corresponding p value shown. *Right*: Pearson correlation coefficients and corresponding p values for each mouse. Correlations were consistently negative across mice (mean r = −0.22, range = −0.36 to −0.16; all mice p ≤ 0.004; n = 5 mice). (c) Venn diagrams showing the overlap between reward-responsive neurons and neurons with a significant fitted response to vagal optogenetic stimulation, shown separately for Gaussian and Sigmoidal response-type groups. Blue dashed circles indicate neurons significantly responsive to reward, pink dashed circles indicate neurons with significant vagal optogenetic frequency-response fits, and purple overlap indicates neurons belonging to both groups. Values show mean ± SEM across mice (n = 5). (d) Percentage of all neurons responsive to reward (*left*) and 15 Hz vagal opto correct rejections (*right*) in the VagalOpto→NoReward task vs. psychophysics task. Bars show mean ± SEM across mice (n = 5); dots indicate individual mice. Reward: p = 0.625, vagal opto: p = 0.06 (Wilcoxon test). (e) Proportion of neurons that significantly responded both at 15 Hz and at each lower frequency (per mouse; gray lines show individual mice, red bars show group mean ± SEM, n = 5). (f) Spatial organization of responsive neurons in psychophysics, VagalOpto→NoReward, and VagalOpto→Reward tasks. *Left*: example two-photon field of view with neurons colored based on their responsiveness to each condition (red: vagal opto; blue: reward). *Right*: example histogram from the same mouse of pairwise distances between significantly responsive neurons for the indicated condition pairs. Note the absence of clear topographic organization. (g) Spatial clustering of neurons responding to reward and vagal optogenetic stimulation across different tasks. *Top*: mean difference in pairwise distance (within minus across categories) per mouse; the shaded band indicates the pooled 95% null confidence interval, and the red line shows the pooled mean. *Bottom*: clustering separability quantified by AUC for same vs. different condition pairs per mouse. Forest plot p values: two-sided permutation on the within minus across distance, with label shuffling and subsampling to the smaller group size. Clustering p values: same procedure on the AUC of same minus different neighbor counts across radii. *p<0.05, **p<0.01, ***p<0.001, ns: p≥0.05.

## Notes

### Competing Interest Statement

The authors have declared no competing interest.

### Summary of Updates

updated title; figure1 revised ; figure2 revised ; figure3 revised ; figure4 revised ; figureS1 revised ; figuresS1 revised

## References

1. Bonaz, B. et al. Diseases, Disorders, and Comorbidities of Interoception. Trends Neurosci 44, 36–51 (2021).

2. Quadt, L., Critchley, H. D. & Garfinkel, S. N. The neurobiology of interoception in health and disease: Neuroscience of interoception. Ann. N. Y. Acad. Sci. 1428, 112–128 (2018).

3. Khalsa, S. S. et al. Interoception and Mental Health: A Roadmap. Biol. Psychiatry Cogn. Neurosci. Neuroimaging 3, 501–513 (2018).

4. Craig, A. D. How do you feel? Interoception: the sense of the physiological condition of the body. Nat. Rev. Neurosci. 3, 655–666 (2002).

5. Sammons, M. et al. Brain-body physiology: Local, reflex, and central communication. Cell 187, 5877–5860 (2024).

6. Owens, A. P., Allen, M., Ondobaka, S. & Friston, K. J. Interoceptive inference: From computational neuroscience to clinic. Neurosci. Biobehav. Rev.90, 174–183 (2018).

7. Khalsa, S. S., Berner, L. A. & Anderson, L. M. Gastrointestinal Interoception in Eating Disorders: Charting a New Path. Curr. Psychiatry Rep. 24, 47–60 (2022).

8. Barrett, L. F. & Simmons, W. K. Interoceptive predictions in the brain. Nat. Rev. Neurosci. 16, 416–426 (2015).

6. Garfinkel, S. N., Seth, A. K., Barrett, A. B., Suzuki, K. & Critchley, H. D. Knowing your own heart: Distinguishing interoceptive accuracy from interoceptive awareness. Biol. Psychol. 104, 65–74 (2015).

10. Schandry, R. & Weitkunat, R. Enhancement of heartbeat-related brain potentials through cardiac awareness training. Int. J. Neurosci. 53, 243–253 (1660).

11. Schillings, C., Karanassios, G., Schulte, N., Schultchen, D. & Pollatos, O. The Effects of a 3-Week Heartbeat Perception Training on Interoceptive Abilities. Front. Neurosci. 16, (2022).

12. Khalsa, S. S., Rudrauf, D., Hassanpour, M. S., Davidson, R. J. & Tranel, D. The practice of meditation is not associated with improved interoceptive awareness of the heartbeat. Psychophysiology 57, e13476 (2020).

13. Seguias, L. & Tapper, K. The effect of mindful eating on subsequent intake of a high calorie snack. Appetite 121, 63–100 (2018).

14. O’Reilly, G. A., Cook, L., Spruijt-Metz, D. & Black, D. S. Mindfulness-based interventions for obesity-related eating behaviours: a literature review. Obes. Rev. 15, 453–461 (2014).

15. Mason, A. E. et al. Effects of a mindfulness-based intervention on mindful eating, sweets consumption, and fasting glucose levels in obese adults: data from the SHINE randomized controlled trial. J. Behav. Med. 39, 201–213 (2016).

16. Berntson, G. G. & Khalsa, S. S. Neural Circuits of Interoception. Trends Neurosci. 44, 17–28 (2021).

17. Mountcastle, V. B., Talbot, W. H., Darian-Smith, I. & Kornhuber†, H. H. Neural Basis of the Sense of Flutter-Vibration. Science 155, 567–600 (1667).

18. Romo, R. & Salinas, E. Flutter Discrimination: neural codes, perception, memory and decision making. Nat. Rev. Neurosci. 4, 203–218 (2003).

19. Stüttgen, M. C., Schwarz, C. & Jäkel, F. Mapping Spikes to Sensations. Front. Neurosci. 5, (2011).

20. Gold, J. I. & Shadlen, M. N. The Neural Basis of Decision Making. Annu. Rev. Neurosci. 30, 535–574 (2007).

21. Parker, A. J. & Newsome, W. T. SENSE AND THE SINGLE NEURON: Probing the Physiology of Perception. Annu. Rev. Neurosci. 21, 227–277 (1668).

22. Carandini, M. & Churchland, A. K. Probing perceptual decisions in rodents. Nat. Neurosci. 16, 824–831 (2013).

23. Verdonk, C., Ajijola, O. A. & Khalsa, S. S. Toward a multidisciplinary neurobiology of interoception and mental health. Curr. Opin. Neurobiol. 94, 103084 (2025).

24. Mayeli, A. et al. Parieto-occipital ERP indicators of gut mechanosensation in humans. Nat. Commun. 14, 3368 (2023).

25. Srinivasan, S. S. et al. A vibrating ingestible bioelectronic stimulator modulates gastric stretch receptors for illusory satiety. Sci. Adv. 9, eadj3003 (2023).

26. Berthoud, H. R., Blackshaw, L. A., Brookes, S. J. H. & Grundy, D. Neuroanatomy of extrinsic afferents supplying the gastrointestinal tract. Neurogastroenterol. Motil. 16, 28–33 (2004).

27. Zagorodnyuk, V. P., Chen, B. N. & Brookes, S. J. H. Intraganglionic laminar endings are mechano-transduction sites of vagal tension receptors in the guinea-pig stomach. J. Physiol. 534, 255–268 (2001).

28. Travagli, R. A., Hermann, G. E., Browning, K. N. & Rogers, R. C. Brainstem circuits regulating gastric function. Annu Rev Physiol 68, 276–305 (2006).

26. Kim, M., Heo, G. & Kim, S.-Y. Neural signalling of gut mechanosensation in ingestive and digestive processes. Nat. Rev. Neurosci. 23, 135–156 (2022).

30. Bai, L. et al. Genetic Identification of Vagal Sensory Neurons That Control Feeding. Cell 179, 1126–1143.e23 (2016).

31. Prescott, S. L. & Liberles, S. D. Internal senses of the vagus nerve. Neuron 110, 576–566 (2022).

32. Borgmann, D. et al. Gut-brain communication by distinct sensory neurons differently controls feeding and glucose metabolism. Cell Metab. 33, 1466–1482.e7 (2021).

33. Williams, E. K. et al. Sensory Neurons that Detect Stretch and Nutrients in the Digestive System. Cell 166, 206–221 (2016).

34. Chang, R. B., Strochlic, D. E., Williams, E. K., Umans, B. D. & Liberles, S. D. Vagal Sensory Neuron Subtypes that Differentially Control Breathing. Cell 161, 622–633 (2015).

35. Min, S. et al. Arterial Baroreceptors Sense Blood Pressure through Decorated Aortic Claws. Cell Rep. 29, 2162–2201.e3 (2016).

36. Prescott, S. L., Umans, B. D., Williams, E. K., Brust, R. D. & Liberles, S. D. An Airway Protection Program Revealed by Sweeping Genetic Control of Vagal Afferents. Cell 181, 574–586.e14 (2020).

37. Zhao, Q., et al. A multidimensional coding architecture of the vagal interoceptive system. Nature 603, 878– 884 (2022).

38. Paton, J. F. The Sharpey-Schafer prize lecture: nucleus tractus solitarii: integrating structures. Exp. Physiol. 84, 815–833 (1666).

36. Ran, C., Boettcher, J. C., Kaye, J. A., Gallori, C. E. & Liberles, S. D. A brainstem map for visceral sensations. Nature 609, 320–326 (2022).

40. Roelofs, T. J. M. et al. Good taste or gut feeling? A new method in rats shows oro-sensory stimulation and gastric distention generate distinct and overlapping brain activation patterns. Int. J. Eat. Disord. 54, 1116– 1126 (2021).

41. Wang, G.-J. et al. Gastric distention activates satiety circuitry in the human brain. NeuroImage 39, 1824– 1831 (2008).

42. Cao, J. et al. Gastric Stimulation Drives Fast BOLD Responses of Neural Origin. NeuroImage 197, 200–211 (2016).

43. Engelen, T., Solcà, M. & Tallon-Baudry, C. Interoceptive rhythms in the brain. Nat. Neurosci. 26, 1670–1684 (2023).

44. Levakov, G., Ganor, S. & Avidan, G. Reliability and validity of brain-gastric phase synchronization. Hum. Brain Mapp. 44, 4656–4666 (2023).

45. Gogolla, N. The insular cortex. Curr. Biol. 27, R580–R586 (2017).

46. Livneh, Y. & Andermann, M. L. Cellular activity in insular cortex across seconds to hours: Sensations and predictions of bodily states. Neuron 109, 3576–3563 (2021).

47. Bailey, P. & Bremer, F. A SENSORY CORTICAL REPRESENTATION OF THE VAGUS NERVE: WITH A NOTE ON THE EFFECTS OF LOW BLOOD PRESSURE ON THE CORTICAL ELECTROGRAM. J. Neurophysiol. 1, 405– 412 (1638).

48. Ogawa, H., Ito, S., Murayama, N. & Hasegawa, K. Taste area in granular and dysgranular insular cortices in the rat identified by stimulation of the entire oral cavity. Neurosci. Res. 9, 166–201 (1660).

46. Saper, C. B. The Central Autonomic Nervous System: Conscious Visceral Perception and Autonomic Pattern Generation. Annu. Rev. Neurosci. 25, 433–466 (2002).

50. Yamamoto, T., Matsuo, R. & Kawamura, Y. Localization of cortical gustatory area in rats and its role in taste discrimination. J Neurophysiol 44, 440–55 (1680).

51. Allen, G. V., Saper, C. B., Hurley, K. M. & Cechetto, D. F. Organization of visceral and limbic connections in the insular cortex of the rat. J. Comp. Neurol. 311, 1–16 (1661).

52. Cechetto, D. F. & Saper, C. B. Evidence for a viscerotopic sensory representation in the cortex and thalamus in the rat. J. Comp. Neurol. 262, 27–45 (1687).

53. Hanamori, T., Kunitake, T., Kato, K. & Kannan, H. Responses of Neurons in the Insular Cortex to Gustatory, Visceral, and Nociceptive Stimuli in Rats. J. Neurophysiol. 79, 2535–2545 (1668).

54. Gehrlach, D. A. et al. A whole-brain connectivity map of mouse insular cortex. eLife 9, e55585 (2020).

55. Critchley, H. D., Wiens, S., Rotshtein, P., Öhman, A. & Dolan, R. J. Neural systems supporting interoceptive awareness. Nat. Neurosci. 7, 186–165 (2004).

56. Craig, A. D. B. How do you feel--now? The anterior insula and human awareness. Nat. Rev. Neurosci. 10, 56– 70 (2006).

57. Simmons, W. K. et al. Keeping the body in mind: Insula functional organization and functional connectivity integrate interoceptive, exteroceptive, and emotional awareness. Hum. Brain Mapp. 34, 2644–2658 (2013).

58. Koren, T. & Rolls, A. Immunoception: Defining brain-regulated immunity. Neuron 110, 3425–3428 (2022).

56. Maffei, A., Haley, M. & Fontanini, A. Neural processing of gustatory information in insular circuits. Curr. Opin. Neurobiol. 22, 706–716 (2012).

60. Carleton, A., Accolla, R. & Simon, S. A. Coding in the mammalian gustatory system. Trends Neurosci. 33, 326–334 (2010).

61. Kayyal, H. et al. Retrieval of conditioned immune response in male mice is mediated by an anterior– posterior insula circuit. Nat. Neurosci. 28, 586–601 (2025).

62. Koren, T. et al. Insular cortex neurons encode and retrieve specific immune responses. Cell 184, 5602–5615.e17 (2021).

63. Jones, L. M., Fontanini, A. & Katz, D. B. Gustatory processing: a dynamic systems approach. Curr Opin Neurobiol 16, 420–8 (2006).

64. Vestergaard, M., Carta, M., Güney, G. & Poulet, J. F. A. The cellular coding of temperature in the mammalian cortex. Nature 614, 725–731 (2023).

65. de Araujo, I. E., Schatzker, M. & Small, D. M. Rethinking Food Reward. Annu Rev Psychol 71, 136–164 (2020).

66. Lavi, K., Jacobson, G. A., Rosenblum, K. & Lüthi, A. Encoding of Conditioned Taste Aversion in Cortico-Amygdala Circuits. Cell Rep. 24, 278–283 (2018).

67. Accolla, R. & Carleton, A. Internal body state influences topographical plasticity of sensory representations in the rat gustatory cortex. Proc. Natl. Acad. Sci. 105, 4010–4015 (2008).

68. Katz, D. B., Simon, S. A. & Nicolelis, M. A. L. Dynamic and Multimodal Responses of Gustatory Cortical Neurons in Awake Rats. J. Neurosci. 21, 4478–4486 (2001).

66. Levitan, D. et al. Single and population coding of taste in the gustatory cortex of awake mice. J. Neurophysiol. 122, 1342–1356 (2016).

70. Grossman, S. E., Fontanini, A., Wieskopf, J. S. & Katz, D. B. Learning-related plasticity of temporal coding in simultaneously recorded amygdala-cortical ensembles. J Neurosci 28, 2864–73 (2008).

71. Prilutski, Y. & Livneh, Y. Physiological Needs: Sensations and Predictions in the Insular Cortex. Physiology 38, 73–81 (2023).

72. Rogers-Carter, M. M. & Christianson, J. P. An insular view of the social decision-making network. Neurosci Biobehav Rev 103, 116–132 (2016).

73. Kogan, J. F. & Fontanini, A. Learning enhances representations of taste-guided decisions in the mouse gustatory insular cortex. Curr. Biol. https://doi.org/10.1016/j.cub.2024.03.034 (2024) doi:10.1016/j.cub.2024.03.034.

74. Vincis, R., Chen, K., Czarnecki, L., Chen, J. & Fontanini, A. Dynamic Representation of Taste-Related Decisions in the Gustatory Insular Cortex of Mice. Curr. Biol. 30, 1834–1844.e5 (2020).

75. Nicolas, C. et al. Linking emotional valence and anxiety in a mouse insula-amygdala circuit. Nat. Commun. 14, 5073 (2023).

76. Zhao, Z. et al. Cannabinoids regulate an insula circuit controlling water intake. Curr. Biol. 34, 1618–1626.e5 (2024).

77. Stern, S. A. et al. Top-down control of conditioned overconsumption is mediated by insular cortex Nos1 neurons. Cell Metab. 33, 1418–1432.e6 (2021).

78. Gardner, M. P. H. & Fontanini, A. Encoding and Tracking of Outcome-Specific Expectancy in the Gustatory Cortex of Alert Rats. J. Neurosci. 34, 13000–13017 (2014).

76. Samuelsen, C. L., Gardner, M. P. H. & Fontanini, A. Effects of Cue-Triggered Expectation on Cortical Processing of Taste. Neuron 74, 410–422 (2012).

80. Arieli, E., Younis, N. & Moran, A. Distinct Progressions of Neuronal Activity Changes Underlie the Formation and Consolidation of a Gustatory Associative Memory. J. Neurosci. 42, 606–621 (2022).

81. Stuber, G. D. Neurocircuits for motivation. Science 382, 364–368 (2023).

82. Zhao, Z. et al. Direct interoceptive input to the insular cortex shapes learned feeding behavior. 2025.05.13.653866 Preprint at 10.1101/2025.05.13.653866 (2026).

83. Verdonk, C. et al. Gastrointestinal Interoception and Relapse in Anorexia Nervosa. JAMA Psychiatry 83, 602– 611 (2026).

84. Chen, R. et al. Deep brain optogenetics without intracranial surgery. Nat. Biotechnol. 39, 161–164 (2021).

85. Hsueh, B. et al. Cardiogenic control of affective behavioural state. Nature 615, 262–266 (2023).

86. Ichiki, T. et al. Sensory representation and detection mechanisms of gut osmolality change. Nature 602, 468–474 (2022).

87. Scott, K. A. et al. Mechanosensation of the heart and gut elicits hypometabolism and vigilance in mice. Nat. Metab. 7, 263–275 (2025).

88. Hampton, R. F., Jimenez-Gonzalez, M. & Stanley, S. A. Unravelling innervation of pancreatic islets. Diabetologia 65, 1066–1084 (2022).

89. Marshel, J. H. et al. Cortical layer–specific critical dynamics triggering perception. Science 365, eaaw5202 (2016).

90. Bailey, P. & Bremer, F. A sensory cortical representation of the vagus nerve: with a note on the effects of low blood pressure on the cortical electrogram. J. Neurophysiol. 1, 405–412 (1638).

91. Ogawa, H., Ito, S., Murayama, N. & Hasegawa, K. Taste area in granular and dysgranular insular cortices in the rat identified by stimulation of the entire oral cavity. Neurosci Res 9, 166–201 (1660).

92. Bagaev, V. & Aleksandrov, V. Visceral-related area in the rat insular cortex. Auton. Neurosci. 125, 16–21 (2006).

93. Penfield, W. & Faulk, M. E. THE INSULA: FURTHER OBSERVATIONS ON ITS FUNCTION. Brain 78, 445–470 (1655).

94. Adamic, E. M., Teed, A. R., Avery, J., de la Cruz, F. & Khalsa, S. Hemispheric divergence of interoceptive processing across psychiatric disorders. eLife 13, RP62820 (2024).

95. Livneh, Y. et al. Estimation of Current and Future Physiological States in Insular Cortex. Neuron 105, 1064–1111.e10 (2020).

96. Livneh, Y. et al. Homeostatic circuits selectively gate food cue responses in insular cortex. Nature 546, 611– 616 (2017).

97. Britten, K. H., Newsome, W. T., Shadlen, M. N., Celebrini, S. & Movshon, J. A. A relationship between behavioral choice and the visual responses of neurons in macaque MT. Vis. Neurosci. 13, 87–100 (1666).

98. Talpir, I. & Livneh, Y. Stereotyped goal-directed manifold dynamics in the insular cortex. Cell Rep. 43, 114027 (2024).

99. Ryu, B. et al. Chronic loss of inhibition in piriform cortex following brief, daily optogenetic stimulation. Cell Rep. 35, (2021).

100. Mahn, M. et al. High-efficiency optogenetic silencing with soma-targeted anion-conducting channelrhodopsins. Nat. Commun. 9, 4125 (2018).

101. Kusumoto-Yoshida, I., Liu, H., Chen, B. T., Fontanini, A. & Bonci, A. Central role for the insular cortex in mediating conditioned responses to anticipatory cues. Proc. Natl. Acad. Sci. 112, 1160–1165 (2015).

102. Petzschner, F. H., Garfinkel, S. N., Paulus, M. P., Koch, C. & Khalsa, S. S. Computational Models of Interoception and Body Regulation. Trends Neurosci. 44, 63–76 (2021).

103. Paulus, M. P., Feinstein, J. S. & Khalsa, S. S. An Active Inference Approach to Interoceptive Psychopathology. Annu. Rev. Clin. Psychol. 15, 67–122 (2016).

104. Fermin, A. S. R., Friston, K. & Yamawaki, S. An insula hierarchical network architecture for active interoceptive inference. R Soc Open Sci 9, 220226 (2022).

105. Chen, W. G. et al. The Emerging Science of Interoception: Sensing, Integrating, Interpreting, and Regulating Signals within the Self. Trends Neurosci. 44, 3–16 (2021).

106. Allen, M. Unravelling the Neurobiology of Interoceptive Inference. Trends Cogn. Sci. 24, 265–266 (2020).

107. Nikolova, N. et al. The respiratory resistance sensitivity task: An automated method for quantifying respiratory interoception and metacognition. Biol. Psychol. 170, 108325 (2022).

108. Banellis, L., et al. Interoceptive ability is uncorrelated across respiratory and cardiac axes: a large scale psychophysical study. Preprint at 10.31234/osf.io/s56v4_v1 (2025).

109. Brener, J. & Ring, C. Towards a psychophysics of interoceptive processes: the measurement of heartbeat detection. Philos. Trans. R. Soc. B Biol. Sci. 371, 20160015 (2016).

110. Zeng, W.-Z. et al. PIEZOs mediate neuronal sensing of blood pressure and the baroreceptor reflex. Science 362, 464–467 (2018).

111. Greenwood, B. M. & Garfinkel, S. N. Interoceptive Mechanisms and Emotional Processing. Annu. Rev. Psychol. 76, 56–86 (2025).

112. Simmons, W. K. et al. Keeping the body in mind: Insula functional organization and functional connectivity integrate interoceptive, exteroceptive, and emotional awareness. Hum. Brain Mapp. 34, 2644– 2658 (2013).

113. Royer, J. et al. Myeloarchitecture gradients in the human insula: Histological underpinnings and association to intrinsic functional connectivity. NeuroImage 216, 116856 (2020).

114. Tian, Y. & Zalesky, A. Characterizing the functional connectivity diversity of the insula cortex: Subregions, diversity curves and behavior. NeuroImage 183, 716–733 (2018).

115. Nord, C. L., Lawson, R. P. & Dalgleish, T. Disrupted Dorsal Mid-Insula Activation During Interoception Across Psychiatric Disorders. Am. J. Psychiatry 178, 761–770 (2021).

116. Banellis, L., Rebollo, I., Nikolova, N. & Allen, M. Stomach–brain coupling indexes a dimensional signature of mental health. Nat. Ment. Health 3, 866–608 (2025).

117. Akam, T. & Walton, M. E. pyPhotometry: Open source Python based hardware and software for fiber photometry data acquisition. Sci. Rep. 9, 3521 (2016).

118. Buchanan, K. L. et al. The preference for sugar over sweetener depends on a gut sensor cell. Nat. Neurosci. 25, 161–200 (2022).

119. Lau, E. O.-C. et al. Aortic Baroreceptors Display Higher Mechanosensitivity than Carotid Baroreceptors. Front. Physiol. 7, (2016).

120. Salman, I. M. Key challenges in exploring the rat as a preclinical neurostimulation model for aortic baroreflex modulation in hypertension. Hypertens. Res. 47, 366–415 (2024).

121. Hwang, J., Mitz, A. R. & Murray, E. A. NIMH MonkeyLogic: Behavioral control and data acquisition in MATLAB. J. Neurosci. Methods 323, 13–21 (2016).

122. Asaad, W. F. & Eskandar, E. N. A flexible software tool for temporally-precise behavioral control in Matlab. J Neurosci Methods 174, 245–58 (2008).

123. Liu, Y. Y., Guofang Wang, Longhua. Isolation and Analysis of Aortic Arch and Root Lesions in an Atherosclerotic Mouse Model. https://www.jove.com/v/67875/isolation-analysis-aortic-arch-root-lesions-an-atherosclerotic-mouse.

124. Pachitariu, M., et al. Suite2p: Beyond 10,000 Neurons with Standard Two-Photon Microscopy. http://biorxiv.org/lookup/doi/10.1101/061507 (2016) doi:10.1101/061507.

125. Lecoq, J. et al. Removing independent noise in systems neuroscience data using DeepInterpolation. Nat. Methods 18, 1401–1408 (2021).

126. Winkler, I., Debener, S., Müller, K.-R. & Tangermann, M. On the influence of high-pass filtering on ICA-based artifact reduction in EEG-ERP. in 2015 37th Annual International Conference of the IEEE Engineering in Medicine and Biology Society (EMBC) 4101–4105 (2015). doi:10.1109/EMBC.2015.7319296.

127. Campos Viola, F., et al. Semi-automatic identification of independent components representing EEG artifact. Clin. Neurophysiol. 120, 868–877 (2006).

